# Covalent Leader Peptide Probes Enable Proteome-Level Mapping of RiPP Enzymes beyond Biosynthetic Gene Clusters

**DOI:** 10.64898/2026.01.17.700038

**Authors:** Lan Wang, Boning Wang, Ying Wang, Yundan Zheng, Yinzheng Xia, Xiangqian Xie, Ciji Wang, Tian Tian, Jing Zhao, Huijie Pan, Huan Wang

## Abstract

Genome-based analyses have enabled widespread discovery and biosynthetic annotation of ribosomally synthesized and post-translationally modified peptides (RiPPs) pathways, yet often fail to capture all participating enzymes, particularly those encoded outside canonical biosynthetic gene clusters. In this study, leveraging the central role of leader peptides (LPs) in RiPP enzyme–substrate recognition, we developed a substrate-guided chemical–proteomic strategy using covalent LP probes to directly identify LP-binding enzymes from native proteomes. Using the LctA-LctM and PatE-LynD systems, we demonstrate that LP probes carrying a photocrosslinker engage enzymes at their authentic LP-recognition sites while preserving full catalytic competence. Application of this strategy to a model strain *Streptomyces sparsogenes* uncovered lanthipeptide synthetases encoded outside canonical biosynthetic gene clusters and revealed graded cross-cluster activities among multiple RiPP pathways. This work establishes a proof-of-concept model platform for proteome-level discovery of LP-binding RiPP enzymes and highlights cross-cluster enzymatic crosstalk as a latent mechanism for structural diversification.

## INTRODUCTION

Ribosomally synthesized and post-translationally modified peptides (RiPPs) represent a broad class of natural products distinguished by exceptional structural diversity.^1^ This diversity arises from a simple but powerful biosynthetic logic in which a precursor peptide—typically composed of an N-terminal leader peptide (LP) and a C-terminal core peptide (CP)—is processed by LP-recognizing tailoring enzymes that install modifications such as dehydration, cyclization, and thioether or ester bond formation (Figure 1A).^2^ The LP usually serves as a universal recognition module that recruits biosynthetic enzymes, modulates catalytic activation, and enforces pathway fidelity.^3^ This LP-directed biosynthetic logic has been extensively characterized in most RiPP systems and has long served as the conceptual foundation for understanding RiPP biosynthesis and bioengineering.^4, 5^

**Figure 1.**
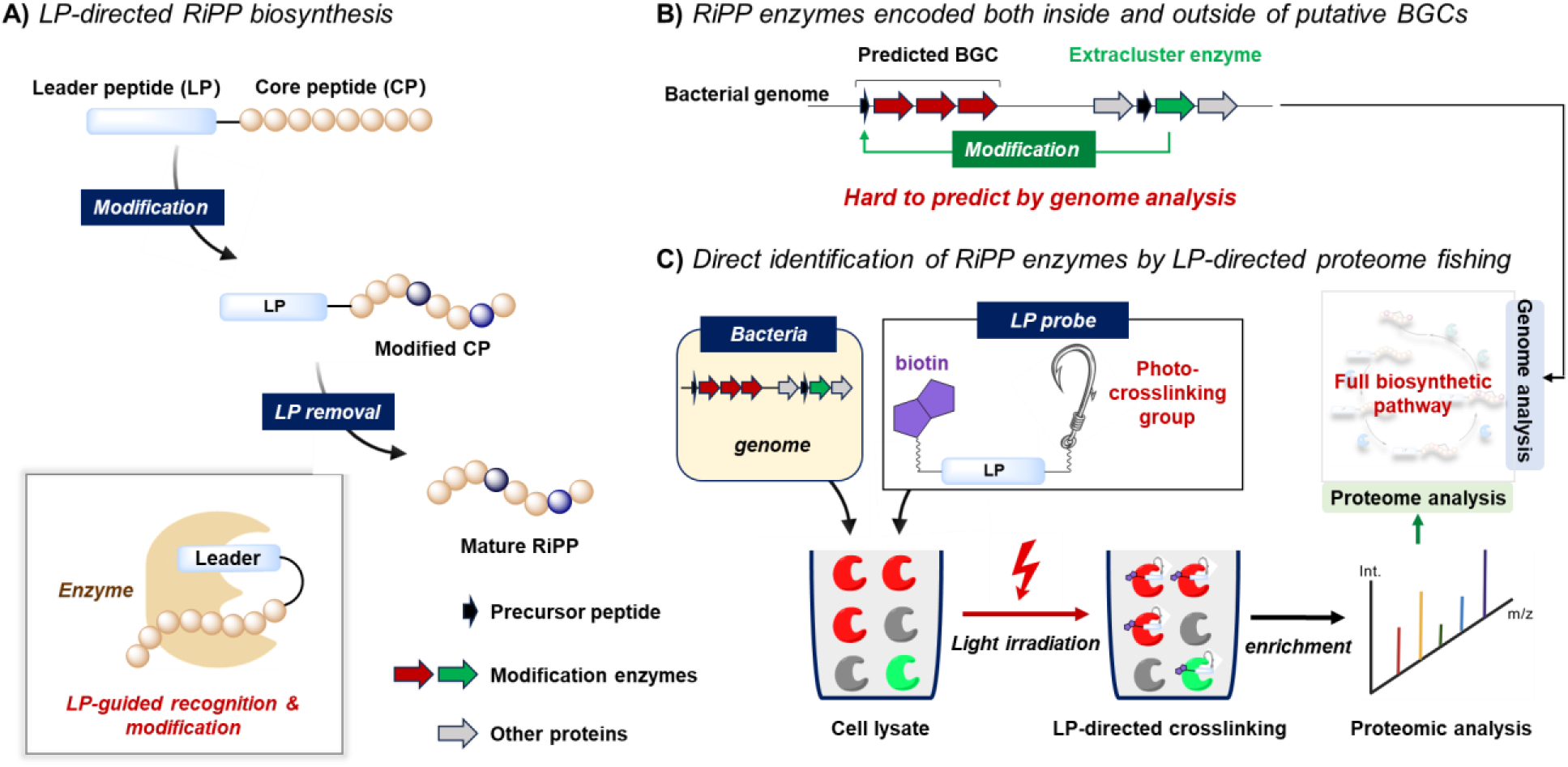
Overview of leader peptide (LP)-mediated recognition in RiPP biosynthesis and the strategy developed in this work to identify LP-binding enzymes. (A) LP-directed enzymatic modification in RiPP biosynthesis. (B) RiPP enzymes encoded both inside and outside of putative BGCs. (C) This work: Direct identification of RiPP enzymes by covalent LP probes.

The discovery and biosynthetic characterization of RiPP natural products have advanced rapidly in recent years, driven largely by the maturation of genome-mining technologies.^6–8^ These computational approaches enable the prediction of precursor peptides and associated tailoring enzymes directly from genomic sequences. Central to these strategies is the assumption that precursor peptides and their cognate modification enzymes are co-localized within the same biosynthetic gene cluster (BGC), allowing genomic context to serve as the primary guide for pathway reconstruction (Figure 1B).

However, emerging genomic and biochemical evidence increasingly suggests that RiPP biosynthetic networks can be far more complex than this classical cluster-centric paradigm implies. For example, the procchlorosin-producing cyanobacterium, encodes more than forty precursor peptide genes dispersed throughout the genome, distant from the sole modification enzyme ProcM.^9^ The biosynthesis of cypepeptins similarly depends on an orphan LanKC enzyme located outside the annotated precursor cluster,^10^ and the maturation of citrulassin A requires a peptidylarginine deiminase encoded ∼20 kb away from its BGC.^8^ Large-scale correlational network analyses further indicate that the processing of many class III lanthipeptides involves “hidden” metallopeptidases encoded distally from their associated clusters.^11, 12^ Together, these examples reveal that RiPP maturation often engages extra-cluster enzymes, distributed auxiliary factors, and cross-cluster crosstalk—features that considerably expand the biochemical landscape beyond what is apparent from BGC organization.

These emerging patterns expose a critical methodological gap: genomic context alone cannot determine which proteins in a native organism physically recognize a given RiPP precursor or functionally act upon it. This challenge is further exemplified by several important RiPP families, such as darobactins and methanobactins, whose BGCs lack identifiable proteases or other key tailoring enzymes, making their biosynthetic logic difficult to reconstruct from genomic information alone and substantially hindering efforts toward rational pathway elucidation and molecule production. Given these limitations at the genomic level, proteome-level interrogation offers a powerful complementary strategy to directly identify the proteins that engage a precursor peptide *in vivo*. Chemical proteomic methods typically rely on functionalized probes to capture interacting proteins in complex lysates, followed by affinity enrichment and mass-spectrometric identification.^13^ Because the leader peptide (LP) functions as the universal binding module that recruits nearly all RiPP biosynthetic enzymes, probes derived from native LP sequences provide a direct and biologically grounded means to map the operative biosynthetic machinery. However, the inherently weak and transient nature of LP-enzyme interactions—typically in the micromolar affinity range—undermines the effectiveness of conventional noncovalent pull-down or affinity-enrichment methods. These challenges underscore the need for a covalent, LP-based probe capable of stabilizing native interactions and enabling robust identification of the full complement of RiPP-modifying enzymes, including those encoded outside canonical BGC boundaries.

Here, we present a covalent LP-probe strategy that addresses these challenges by installing photo-reactive crosslinking groups onto LP sequences to capture LP-binding enzymes under light irradiation (Figure 1C). Using the LctA-LctM from the lacticin 481 biosynthesis and PatE-LynD from the cyanobactin biosynthesis as models,^14^ we demonstrate that these covalent LP probes engage enzymes at their authentic LP-binding sites with high specificity while preserving and promoting catalytic competence. We further apply this strategy to a model strain *Streptomyces sparsogenes* encoding multiple RiPP BGCs, where LP-directed photolabeling reveals cryptic modification enzymes and uncovers previously unrecognized cross-cluster enzymatic interactions. This work establishes a general chemical-proteomic platform for mapping RiPP biosynthetic networks and provides new insights into the distributed and interconnected nature of RiPP maturation across microbial genomes.

## RESULTS AND DISCUSSION

### Validation of covalent leader-peptide probes for specific enzyme capture

To evaluate whether LP probes bearing photoreaction crosslinkers can function as faithful probes, we selected a model system that allows assessment of three key criteria: (i) the modified LP must retain specific binding to its cognate enzyme, (ii) covalent capture should occur only at the authentic LP-binding interface rather than at off-pathway sites, and (iii) the LP-enzyme complex must adopt a functionally productive binding mode that can be experimentally verified. The LctA-LctM system from the biosynthesis of lanthipeptide lacticin 481 fulfills all three requirements (Figure 2A): LctM binds the LctA leader peptide (LctA_LP_) and is activated upon LP engagement to catalyze dehydration and cyclization of the cognate core peptide, enabling a functional readout of correct LP binding.^15, 16^

**Figure 2.**
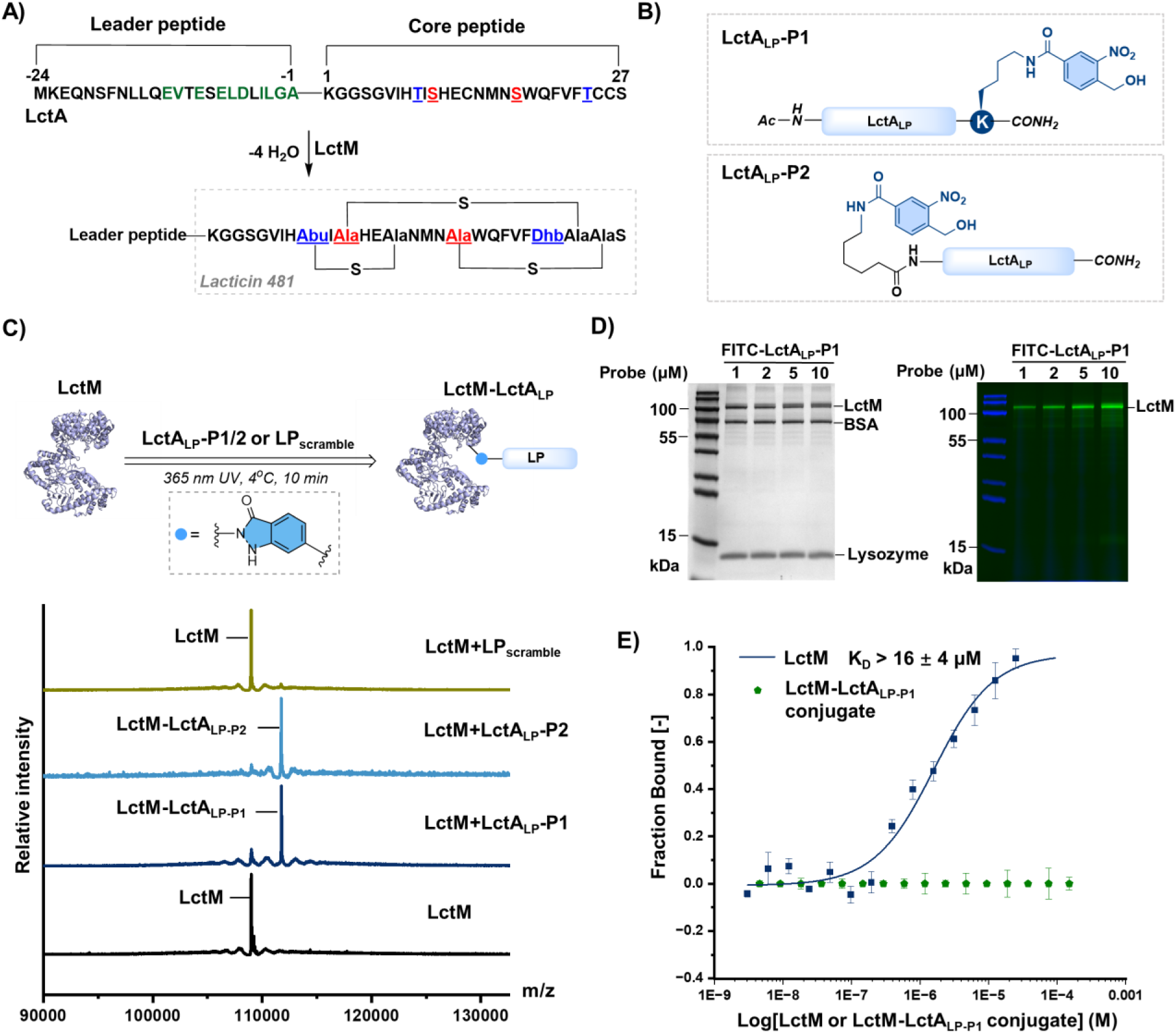
Covalent LctA_LP_ probes capture LctM in an efficient and specific manner. (A) Modification of LctA peptide by LctM. (B) Structures of LctA_LP_ probes. (C) MS analysis of photo-crosslinking between LctM and LctA_LP_-P1, LctA_LP_-P2, or LP_scramble_. LctM: calcd: 108998 Da, obs.: 108996 Da. LctM-LctA_LP-P1_: calcd: 111773 Da, obs.: 111775 Da. LctM-LctA_LP-P2_: calcd: 111714 Da, obs.: 111716 Da. (D) Specific photo-labelling of LctM by FITC-LctA_LP_-P1 in mixed protein samples. (Left) Coomassie-stained SDS–PAGE gel. (Right) Fluorescence scan showing selective, dose-dependent labeling of LctM. (E) MST binding curves for LctM and the LctM–LctA_LP-P1_ conjugate titrated against LctA_LP_ peptide.

To develop an effective covalent LP probe, we first needed to select an appropriate photoreactive crosslinker and determine where such groups should be installed on the leader peptide. Structural predictions of the LctA-LctM complex indicated that the N- and C-terminal regions of the LctA_LP_ are positioned near LctM surface patches containing residues for photochemical coupling (Figure S1), including Lys, Arg and Tyr, suggesting that both termini of LctA_LP_ provide viable positions for installing a photoreactive moiety. We next consider the choice of photo-crosslinking groups commonly used in chemical proteomics. Diazirines and benzophenones offer broad reactivity but generally exhibit low photolabeling efficiencies (often below 30%).^17^ In contrast, *o*-nitrobenzyl azide (*o*-NBA) specifically reacts with nearby Lys residues with high efficiency, frequently approaching near-quantitative crosslinking.^18^

Guided by these considerations, we selected *o*-NBA as the photoreactive crosslinker and installed it at either the N- or C-terminus of LctA_LP_, yielding two probes: LctA_LP_-P1 and LctA_LP_-P2 (Figure 2B). Microscale thermophoresis (MST) analysis confirmed that installation of the *o*-NBA group did not perturb LP-enzyme recognition (Figure S2). In contrast, an NBA-modified scrambled peptide (LP_scramble_), which contains the same amino acid composition as LctA_LP_ but in randomized order, showed negligible interaction with LctM (Figure S2). Upon 365 nm irradiation of LctM in the presence of probe (2.5 equiv.), both LctA_LP_-P1 and LctA_LP_-P2 underwent efficient mono-crosslinking to LctM, whereas LP_scramble_ yielded no detectable modification under the same conditions (Figure 2C). These results confirm that *o*-NBA–mediated photolabeling occurs in an LP-dependent and sequence-specific manner. To assess the protein specificity of the photoreaction, we synthesized a fluorescently labeled probe (FITC-LctA_LP_-P1) and performed competition assays in the presence of excess bovine serum albumin (BSA) and lysozyme. Fluorescence imaging of SDS–PAGE gels showed that LctA_LP_-P1 crosslinked to LctM selectively, with negligible labeling of the competing proteins (Figure 2D), confirming that photocoupling is driven by specific LctM-LctA_LP_ recognition rather than nonspecific surface reactivity.

To verify that crosslinking occurred within the authentic LP-binding pocket, the purified LctM-LctA_LP_ conjugate was subjected to MST titration with free LctA_LP_. No measurable binding was detected (Figure 2E), suggesting that the LP-binding site of LctM is occupied by the covalently tethered probe. LctM is known to require LP engagement to achieve catalytic activation toward the LctA core peptide (LctA_cp_), as established by both in-trans assays and LctM-LP fusion (LctCE) constructs (Figure 3A).^15, 16^ We therefore examined whether the LctM-LctA_LP_ conjugate retains this LP-dependent catalytic behavior. Remarkably, the covalent conjugate modified LctA_cp_ with efficiency comparable to that of the LctM–LP fusion enzyme LctCE, generating fully dehydrated and cyclized products with the characteristic ring topology of lacticin 481 (Figure 3B, C and Figure S3). Collectively, these results confirm that the photo-affinity LctA_LP_ probes bind specifically to the native LP-recognition site of LctM and preserve its full catalytic functionality after covalent crosslinking.

**Figure 3.**
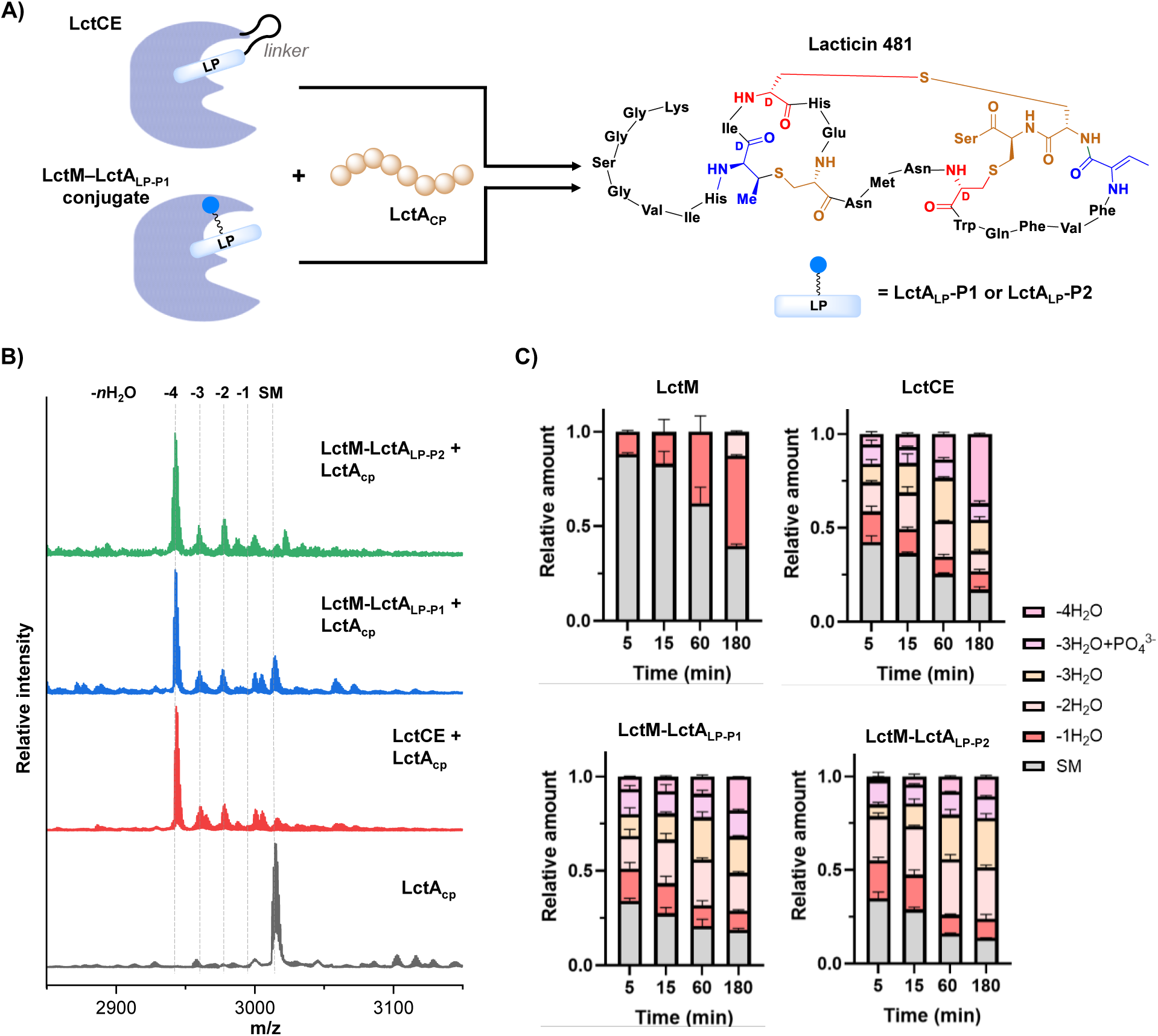
Functional competence of LctM-LctA_LP_ conjugates to modify LctA core peptide. (A) LctCE and LctM-LctA_LP_ conjugates are equally competent in modifying LctA_CP_. (B) MALDI-TOF analysis of LctA_CP_ modified by LctM, LctCE, LctM-LctA_LP-P1_, and LctM-LctA_LP-P2_. Reaction conditions: 20 μM LctA_CP_, 2 μM enzyme, 10 mM MgCl_2_, 2 mM ATP, 25 μg/mL BSA, and 50 mM Tris (pH 7.5) at 25 °C for 3 hours. (C) LC–MS time-course analysis of LctA_CP_ modification by LctM enzymes.

To assess the generality of our covalent LP-probe strategy, we next applied it to the cyanobactin hetero-cyclase LynD and its precursor peptide PatE (Figure 4A).^19^ Like LctM, LynD exhibits only minimal activity toward the PatE core peptide in its free form, but binding of the PatE leader peptide strongly activates the enzyme and restores robust macrocyclization activity. Structural modeling of the LynD-PatE complex indicated that the C-terminal region of the leader peptide inserts into a defined binding pocket containing residues suitable for *o*-NBA-mediated photochemistry (Figure S4). Guided by this model, we installed an *o*-NBA group at the C-terminus of the minimal leader peptide of PatE to generate probe PatE_mLP_-P1. Under optimized irradiation conditions, PatE_mLP_-P1 underwent efficient mono-crosslinking with LynD (Figure 4B, C). The resulting covalent LynD-PatE_LP_ conjugate displayed strong macrocyclization activity toward the PatE core peptide (PatE_CP_), in sharp contrast to the minimal activity of free LynD (Figure 4D), demonstrating that the covalent probe engages LynD at its native LP-recognition pocket and stabilizes its catalytically active state. Taken together with the LctM results, these findings establish covalent LP probes as a general strategy for capturing RiPP modification enzymes directly in their authentic LP-bound state. For LP-activated enzymes, this approach provides a convenient and applicable means to generate catalytically competent, self-activated RiPP enzymes, without the spatial constraints inherent to N- or C-terminal LP-fusion constructs.

**Figure 4.**
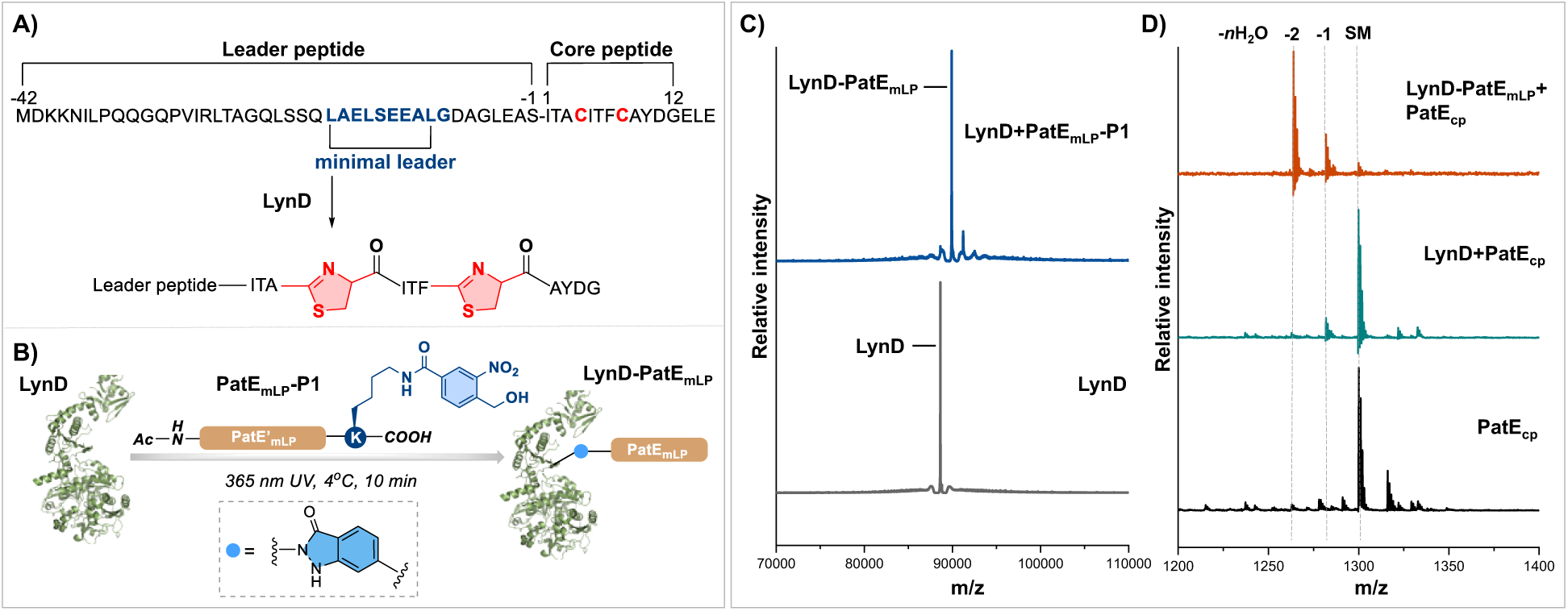
Extension of the covalent LP probe strategy to the cyanobactin macrocyclase LynD. (A) Modification of PatE peptide by cyanobactin hetero-cyclase LynD. (B) Photo-induced conjugation between LynD and the minimal leader peptide probe PatE_mLP_-P1. (C) HRMS analysis of photo-crosslinking between LynD and PatE_mLP_-P1. LynD: calcd: 88636 Da, obs.: 88635 Da. LynD-PatE_mLP_: calcd: 89922 Da, obs.: 89921 Da. (D) MALDI-TOF analysis showing macrocyclization of the PatE_cp_ catalyzed by the crosslinked LynD-PatE_mLP_ complex.

### Covalent LP probes reveal lanthipeptide biosynthetic enzymes beyond canonical gene clusters

Having demonstrated the specificity and functional fidelity of covalent LP probes, we next leveraged this strategy to interrogate the proteome of a native RiPP-producing strain. *S. sparsogenes* ATCC 25498 was selected as a representative system, as its genome encodes multiple RiPP biosynthetic gene clusters as predicted by antiSMASH, including two class III lanthipeptide BGCs (*lan III-1 and lan III-2*), one class IV lanthipeptide BGC (*lan IV*), and one thioamitide BGC (*tat*) (Figure 5A, Figure S5-S8 and Table S3). Among these, the *lan III-1* cluster encodes the lanthipeptide synthetase SpaKC-1 and three precursor peptides SpaA1.1–1.3 (Figure 5B and Figure S6). *In vitro* reconstitution confirmed that SpaKC-1 efficiently modifies SpaA1.1–1.3, generating the corresponding labionin- and lanthionine-containing products (Figure 5B and Figure S10-S15). None of the lanthipeptide BGCs encode detectable peptidases, leaving unresolved the identity of the functional leader-peptide–processing proteases required for maturation of lanthipeptides encoded in this organism.

**Figure 5.**
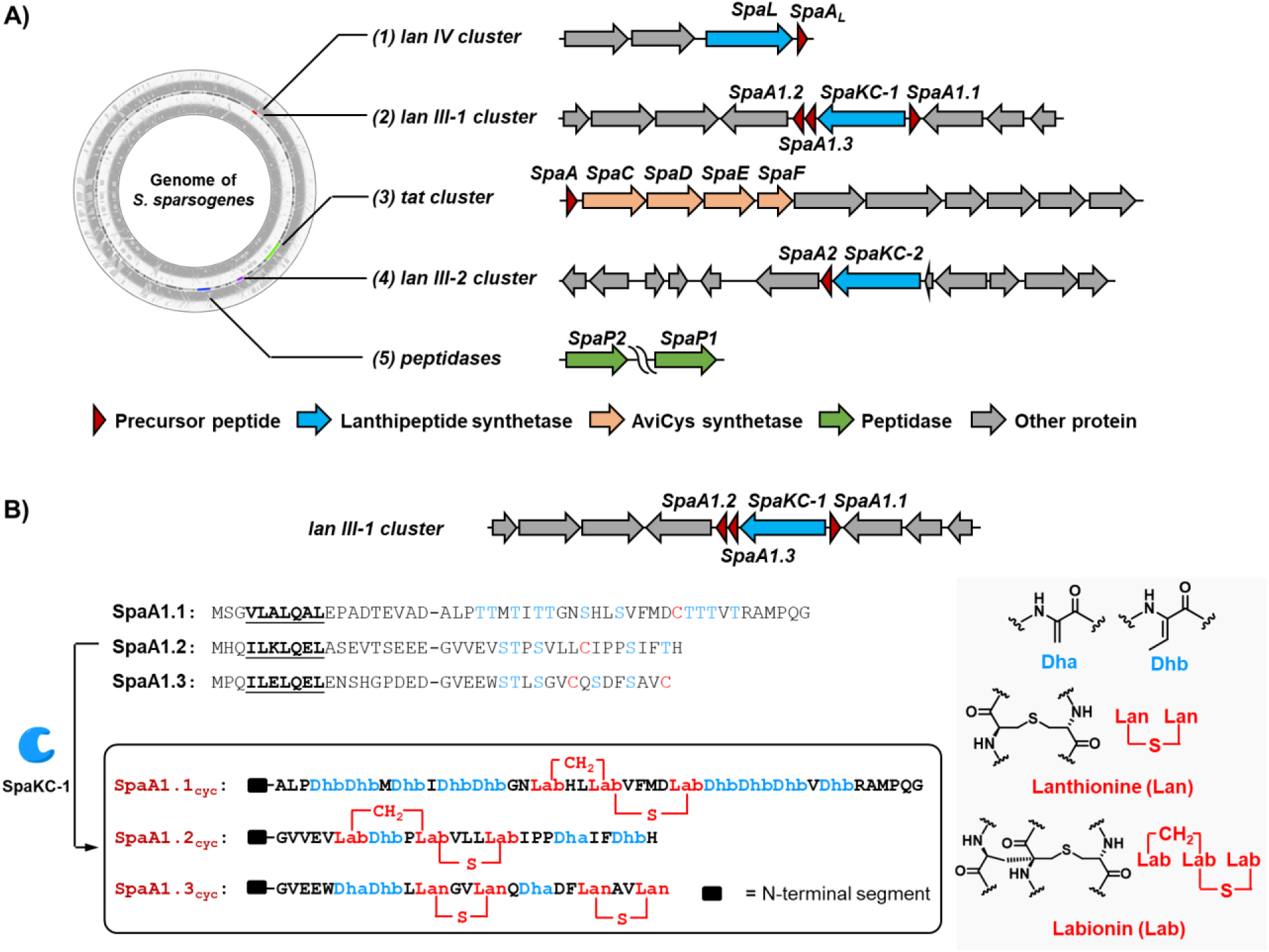
Putative RiPP BGCs encoded in *S. sparsogenes* ATCC 25498 and reconstitution of the activation of SpaKC-1 from the *lan III-1* cluster. (A) Putative RiPP BGCs encoded in S. sparsogenes ATCC 25498. (B) Sequences of the lanthipeptide precursor peptides SpaA1.1, SpaA1.2, and SpaA1.3, together with their corresponding products modified by the cognate lanthipeptide synthetase SpaKC-1 encoded in the *lan III-1* cluster. Conserved recognition sequences within the N-terminal leader peptides are underlined.

To validate the ability of substrate-guided photo-crosslinking probes to retrieve *bona fide* modification enzymes directly from *S. sparsogenes* lysates, we designed two photo-crosslinking probes based on the SpaA1.2 leader peptide: SpaA1.2_LP_-P1, bearing an NBD fluorophore for gel-based detection, and SpaA1.2_LP_-P2, equipped with a biotin handle for affinity enrichment (Figure 6A). When applied to *E. coli* lysates supplemented with SapKC-1 proteins, SpaA1.2_LP_-P1 exhibited selective and concentration-dependent photo-labeling of SpaKC-1 with minimal background from endogenous proteins (Figure 6B), demonstrating that probe reactivity is governed by native LP–enzyme recognition rather than nonspecific adsorption. Having established probe specificity, we next employed the biotinylated probe SpaA1.2_LP_-P2 to *S. sparsogenes* lysates. Following UV-induced crosslinking and streptavidin affinity enrichment, the captured proteins were analyzed by silver staining and LC–MS/MS (Figure 6C and Figure S20A). Ion-mobility MS revealed a dose-dependent enrichment pattern: treatment with 10 μM probe yielded 51 enriched proteins, whereas the 5 μM probe captured 33 proteins. After subtracting proteins present in the no-probe control, these datasets converged on a high-confidence set of 22 specific targets (Figure 6D and Figure S20C). Among the top-ranked hits, four proteins in addition to SpaKC-1 were consistently enriched, including lanthipeptide synthetases, SpaKC-2 from BGC *lan III-2* and SpaL from BGC *lan IV*, and two M1 family metallopeptidases, SpaP1 and SpaP2 (Figure 6E). Notably, *spaP1* and *spaP2* genes are located distally from any annotated RiPP BGC (Figure 5A).

**Figure 6.**
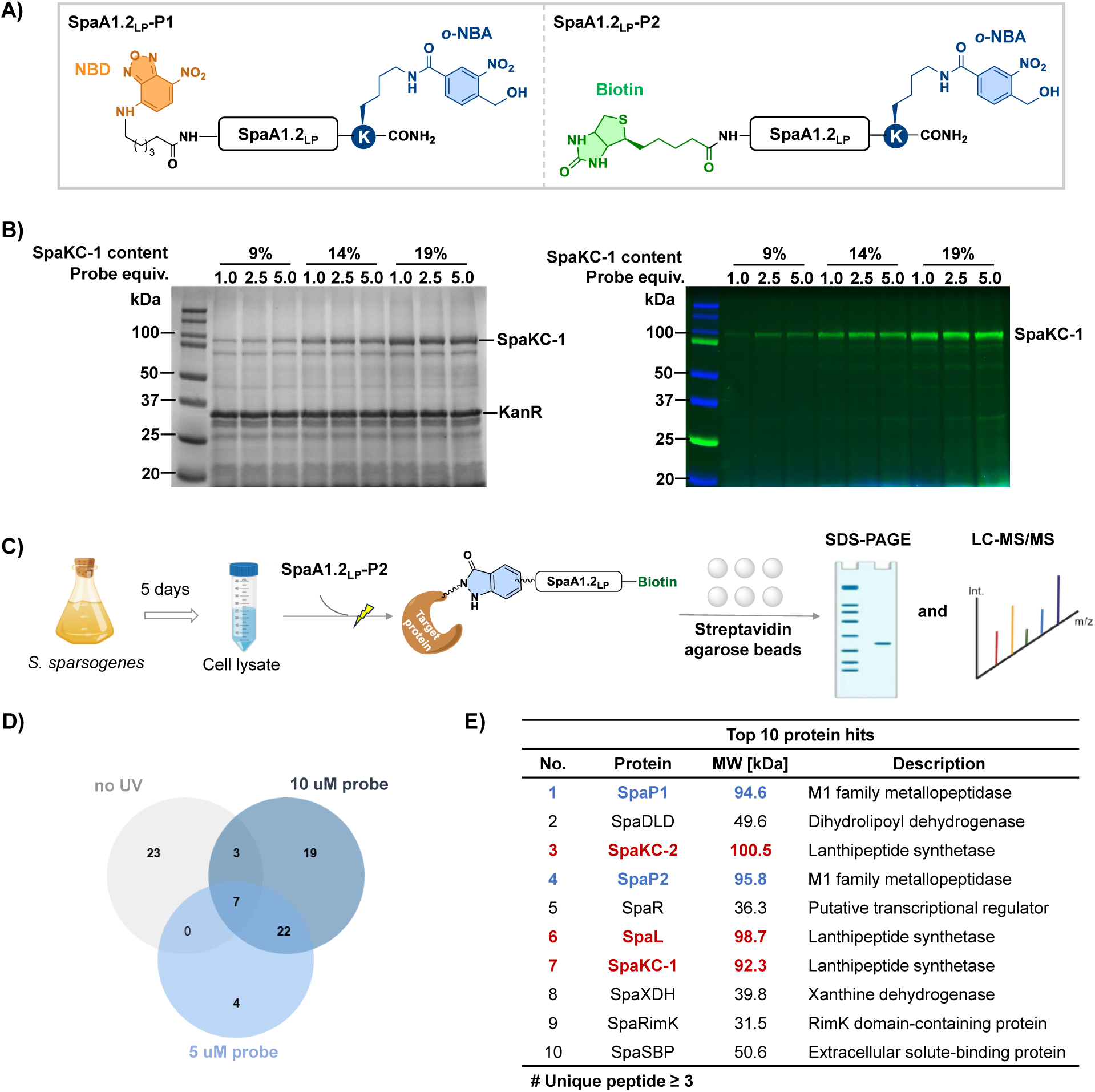
Proteome-level identification of lanthipeptide biosynthetic enzymes beyond canonical BGCs using SpaA1.2-derived covalent LP probes. (A) SpaA1.2-derived probes: NBD-labeled SpaA1.2_LP_-P1 and biotinylated SpaA1.2_LP_-P2. (B) Specific labeling of SpaKC-1 in *E. coli* cell lysates by SpaA1.2_LP_-P1. (Left) Coomassie-stained SDS–PAGE gel. KanR, kanamycin resistance protein. (Right) Alexa 488 fluorescence scan showing minimal nonspecific labeling. (C) Workflow for the identification of RiPP-associated enzymes encoded outside canonical BGCs from the *S. sparsogenes* cell lysate using the SpaA1.2_LP_-P2 probe. (D) Venn diagram showing proteins significantly enriched by SpaA1.2_LP_-P2 in *S. sparsogenes* cell lysate. (E) Top 10 protein hits enriched by SpaA1.2_LP_-P2. Lanthipeptide synthetases are highlighted in red, and M1 family metallopeptidases are highlighted in blue.

To validate the proteomic hits, we assessed whether the lanthipeptide synthetases identified can be efficiently captured by the SpaA1.2_LP_ probes *in vitro*. Using a simplified probe Spa1.2A_LP_-P3, we found that all three enzymes underwent detectable photo-crosslinking under standard conditions, confirming that each can directly engage the SpaA1.2 leader peptide (Figure S21). HRMS further revealed quantitative differences in reactivity: SpaKC-1 and SpaKC-2 displayed high crosslinking efficiency, whereas SpaL showed weaker but still measurable labeling. These results verify that the lanthipeptide synthetases identified in the proteomic screen are LP-binding enzymes and are directly accessible to covalent capture by the SpaA1.2 leader-peptide probe.

We next sought to determine whether the proteases uncovered by LP probe–based proteomics play functional roles in RiPP biosynthesis in *S. sparsogenes*. *In vitro* assays showed that both SpaP1 and SpaP2 efficiently cleaved the leader peptide of (SpaKC-1)–modified SpaA1.2 (SpaA1.2_cyc_). SpaP2 exhibited site-specificity and exclusively targeted the LQ–EL bond, whereas SpaP1 additionally processed the upstream LK–LQ site, indicating a broader cleavage scope (Figure 7A). Beyond this substrate, both proteases were capable of removing N-terminal segments from all lanthipeptide precursor peptides encoded in *S. sparsogenes*, including in SpaA1.1–1.3, SpaA2 and SpaA_L_, demonstrating their roles as promiscuous processing enzymes encoded outside lanthipeptide BGCs (Figure S22-S26). Furthermore, consistent with their binding to SpaA1.2_LP_ probes, both SpaKC-2 and SpaL were able to modify the SpaA1.2 precursor, each generating product profiles distinct from those produced by SpaKC-1 (Figure 7B, C and Figure S37-S40). Taken together, these results demonstrate that covalent LP probes not only recover the cognate lanthipeptide synthetase SpaKC-1 from the *S. sparsogenes* proteome, but also uncover a broader network of biosynthetic enzymes encoded beyond the canonical gene cluster—including the processing proteases SpaP1 and SpaP2—that collectively mediate SpaA1.2 and other precursor peptides maturation. As a proof-of-concept, this system demonstrates the feasibility of using covalent leader-peptide probes to map RiPP biosynthetic interactions and to identify auxiliary enzymes that are not evident from genomic analysis alone.

**Figure 7.**
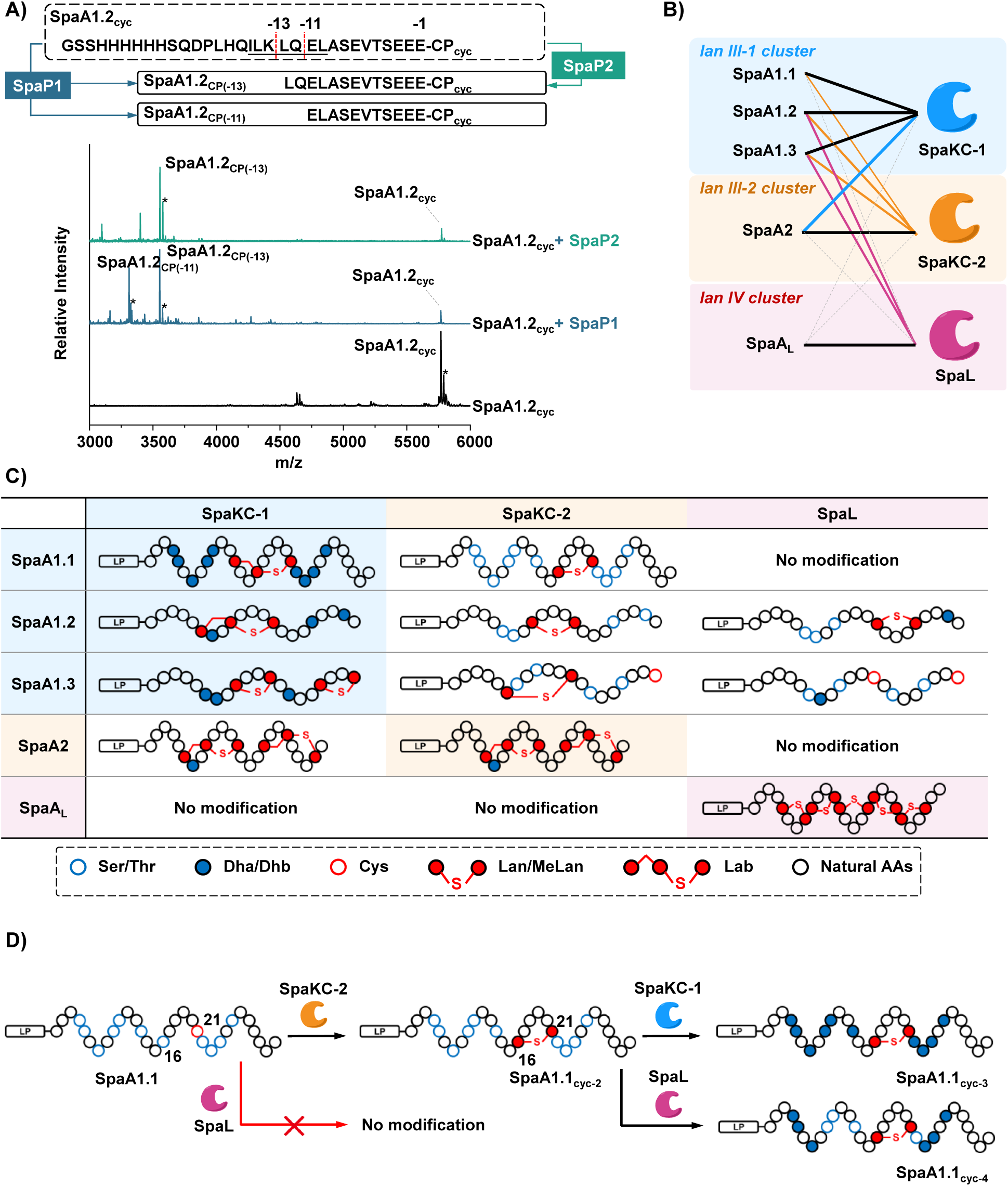
Functional characterization of proteases and lanthipeptide synthetases reveals cross-cluster enzymatic crosstalk and stepwise cascade modification. (A) Endopeptidase activity of metallopeptidases SpaP1 and SpaP2 toward SpaA1.2_cyc_ as analyzed by MALDI MS. [SpaA1.2_cyc_ + H]^+^ Calc. 5762.86 Obs. 5762.94, [SpaA1.2_CP(-13)_ + H]^+^ Calc. 3549.78 Obs. 3549.92, [SpaA1.2_CP(-11)_ + H]^+^ Calc. 3308.64 Obs. 3308.45. (B) Cognate-cluster and cross-cluster modification activities of three lanthipeptide synthetases toward SpaA-derived precursor peptides. Unmodifiable peptides are indicated by dashed lines. (C) Products generated by cognate-cluster and cross-cluster lanthipeptide modification. (D) Proposed stepwise cascade modification of SpaA1.1 mediated by multiple enzymes. Dha/Dhb, dehydroalanine or dehydrobutyrine; Lan/MeLan, lanthionine or methyl lanthionine; Lab, labionin; Natural AAs, natural amino acid residues.

### Cross-cluster crosstalk enables combinatorial biosynthesis of lanthipeptides

Building on the discovery that SpaA1.2 can be modified by lanthipeptide synthetases encoded in different BGCs, we sought to evaluate whether *S. sparsogenes* possesses a broader capacity for cross-cluster RiPP modification. Such behavior parallels well-established phenomena in other classes of natural products,^20–24^ where enzymes originating from distinct BGCs cooperate to expand chemical diversity, yet investigations of this type of crosstalk remain rare within the RiPP family. Our previous study on the *tat* BGC–encoded thioamitide pathway hinted at a similar capability in *S. sparsogenes*: SpaKC-1, the class III lanthipeptide synthetase from the *lan III-1* cluster, was shown to function as a complementary dehydratase in thioamitide biosynthesis, despite not generating new metabolites. This precedent suggests that RiPP enzymes in *S. sparsogenes* may operate beyond their native genetic contexts and may participate in a distributed modification network across multiple RiPP pathways.

To probe whether genes *spaKC-1*, *spaKC-2*, *spaL*, *spaP1*, and *spaP2* identified above might act cooperatively *in vivo*, we first examined their transcriptional profiles during *S. sparsogenes* fermentation. qPCR analysis revealed highly similar temporal expression patterns, with all five genes reaching maximal transcript levels on the fifth day of liquid cultivation (Figure S27). This coordinated expression suggests that these enzymes are co-regulated and supports the hypothesis that they may function within an interconnected biosynthetic framework spanning multiple lanthipeptide gene clusters.

Guided by this evidence of co-expression, we systematically assessed the substrate scope of the three lanthipeptide synthetases toward precursor peptides originating from the three RiPP gene clusters (see Table S4 for precursor peptide sequences). Each enzyme efficiently processed the precursor peptides encoded within its cognate BGC, yielding the labionin- and lanthionine-containing products (Figure 7C and Figure S9-S19). Beyond their native substrates, however, the synthetases displayed markedly different degrees of cross-cluster activity. SpaKC-1 fully dehydrated SpaA2 to form a cyclic product identical to that produced by SpaKC-2, yet it showed no detectable activity toward the class IV precursor SpaA_L_. SpaKC-2 exhibited broader promiscuity, modifying all three precursors from the *lan III-1* cluster and installing a single lanthionine ring in each case, with product profiles distinct from those generated by SpaKC-1. In contrast, SpaL displayed the most stringent specificity: it preferentially modified its native substrate SpaA_L_ and showed only limited activity toward SpaA1.2 and SpaA1.3, introducing 1–2 dehydrations. Notably, SpaL-mediated modification of SpaA1.2 established ring formation in an N-to-C direction, a topology distinct from that produced by SpaKC-1 in its cognate pathway.

Many of the cross-BGC–modified precursor peptides retained unmodified Ser/Thr and Cys residues, raising the possibility that distinct synthetases could act sequentially on the same precursor. For example, SpaKC-2 modified SpaA1.1 by installing only a single dehydration at Ser16, generating a monocyclic intermediate (SpaA1.1_cyc-2_) that remains rich in unmodified Ser/Thr residues. This partially processed peptide remains competent for further editing: SpaKC-1 can further act on SpaA1.1_cyc-2_ to complete dehydration and yield a product (SpaA1.1_cyc-3_) with a ring topology distinct from that generated by SpaKC-1 alone. Notably, although SpaL shows negligible activity toward unmodified SpaA1.1, the SpaKC-2–derived intermediate SpaA1.1_cyc-2_ become an excellent substrate (Figure 7D). SpaL efficiently installs six additional dehydrations to generate a new product SpaA1.1_cyc-4_. The sequential SpaKC-2-to-SpaL modification of SpaA1.1 is further supported by a SpaA1.1-SpaKC-2-SpaL co-expression system in *E. coli* (Figure S35 and S36). Collectively, these results demonstrate that initial processing by one synthetase can prime a precursor for extensive elaboration by another, revealing the latent potential for stepwise, combinatorial RiPP modification across distinct BGCs. Considering intra- and inter-cluster single-and multi-step cross-modification among the three lanthipeptide BGCs in *S. sparsogenes*, a total of 17 distinct products can be generated from a limited set of precursor peptides—far exceeding the classical one-precursor-one-product paradigm and highlighting the substantial combinatorial potential encoded within RiPP biosynthetic networks (Figure S28-S56).

While these activities have been demonstrated only *in vitro* or in heterologous hosts, and therefore do not yet establish that *S. sparsogenes* naturally produces such cross-modified products, they nonetheless uncover an unexpected degree of enzymatic compatibility among its RiPP pathways. Notably, the LP-based photo-crosslinking probes were essential for revealing these noncognate interactions directly from native lysates, allowing us to detect enzyme-substrate relationships that are difficult to infer from genomic context alone. This probe-guided approach thus offers a concise and effective means to expose otherwise hidden biosynthetic connections and to evaluate the combinatorial potential encoded within complex RiPP gene clusters.

## CONCLUSIONS

In this work, we establish covalent LP probes as a substrate-guided chemical-proteomic strategy for discovering RiPP biosynthetic enzymes encoded both inside and outside canonical BGCs, that physically engage precursor peptides in native proteomes. Application of this approach to *S. sparsogenes* uncovered a distributed network of lanthipeptide synthetases and metallopeptidases that participate in precursor peptide maturation. Systematic biochemical reconstitution revealed that these enzymes are not restricted to their cognate gene clusters but instead display graded substrate specificities and the capacity for cross-cluster and sequential modification. These findings uncover a latent layer of combinatorial potential in RiPP biosynthesis that is not readily predictable from genomic context alone.

It is worth noting that, as a proof-of-concept, the present strategy is tailored to LP-binding proteins and enzymes, and does not aim to provide a comprehensive inventory of all factors involved in RiPP maturation. Proteins that act directly on core peptides without LP engagement may therefore escape detection, positioning this approach as complementary to genome-based prediction and functional reconstitution. In addition, the current implementation relies on lysine-reactive photocrosslinking chemistry. Future incorporation of orthogonal photoreactive groups with complementary residue preferences is expected to enable LP-probe cocktails with expanded coverage of LP-binding proteins and enzymes.

Together, this work introduces covalent LP probes as a versatile and extensible tool for mapping substrate-enzyme interactions at the proteome level. Beyond microbial systems, this LP-guided chemical–proteomic framework should be particularly valuable for investigating RiPP pathways in plants and animals, where dispersed gene organization and the absence of well-defined biosynthetic gene clusters often obscure reliable genome-based pathway annotation.

## ASSOCIATED CONTENT

Supporting Information.

Detailed materials and methods for protein expression, purification and biochemical characterization; additional data including sequence alignment, MS and tandem MS analyses.

## AUTHOR INFORMATION

### Author Contributions

H.W. initiated and directed this study. L.W. carried out the design, synthesis of leader peptide probes, photo-crosslinking assays and proteomic analysis. L.W. and B.W. prepared enzymes and peptides and performed biochemical assays. B.W. performed genome mining, gene cluster analysis, cosmid and plasmid construction, fermentation and isolation of peptide natural products. All authors participated in the data analysis and manuscript preparation. All authors have given approval to the final version of the manuscript.

### Notes

The authors declare no competing interest.

## ACKNOWLEDGMENT

This work is supported by NSF of China (Grant 22325702), the Natural Science Foundation of Jiangsu Province (BK20253010, BK20232020 and BG2025036), and Yachen Foundation of Nanjing University. We thank for financial support provided by the State Key Laboratory of Coordination Chemistry. We thank Dr. Hongjuan Chen from the State Key Laboratory of Pharmaceutical Biotechnology for assistance in proteomic analysis.

