## Supporting Information for "Covalent Leader Peptide Probes Enable Proteome-Level Mapping of RiPP Enzymes beyond Biosynthetic Gene Clusters"

### General Information

**Reagents and materials.** Rink amide MBHA resin (0.31 mmol/g) and 2-Cl-Trt resin (0.709 mmol/g) were purchased from GL Biochem (Shanghai, China). All amino acids and other building blocks were obtained from GL Biochem or Bide Pharmatech Ltd. (Shanghai, China) and used without further purification. Coupling reagents (DIC and Oxyma) and trifluoroacetic acid (TFA) were purchased from Adamas-beta (Shanghai, China). MS-grade trypsin was obtained from Beijing Life Proteomic (Beijing, China). All reagents used for proteolytic digestion were LC–MS grade. Unless otherwise noted, general reagents were obtained from Thermo Fisher Scientific (Waltham, MA, USA) or Merck (Rahway, NJ, USA). Oligonucleotides were synthesized by GenScript Biotech (Nanjing, China). Restriction endonucleases were purchased from New England Biolabs (Ipswich, MA, USA). Phanta® Max Master Mix and the ClonExpress II One Step Cloning Kit were purchased from Vazyme Biotech Co., Ltd. (Nanjing, China). *E. coli* DH5α was used for cloning and plasmid propagation, whereas *E. coli* BL21 (DE3) was employed for heterologous protein and peptide expression. Endoprotease GluC was purchased from Roche Biosciences (Basel, Switzerland). Bacterial culture media components were sourced from Thermo Fisher Scientific. *Streptomyces sparsogenes* ATCC 25498 was obtained from the China General Microbiological Culture Collection Center (CGMCC).

**Instruments.** All peptides were synthesized on a CEM Liberty Blue 2.0 automated microwave peptide synthesizer and subsequently purified by HPLC. Semi-preparative HPLC was performed on a Shimadzu system equipped with an LC-20AR semi-preparative solvent delivery unit, a Welch Materials Ultimate® XB-C18 column (5 μm, 10 × 250 mm), and an SPD-20A UV/Vis detector. UPLC-MS analyses were performed with an ACQUITY UPLC I-Class Plus -coupled Xevo G2-XS QTOF MS System (Waters) using ACQUITY Premier Peptide BEH C18 Column (1.7 μm, 2.1 x 150 mm). High-resolution protein mass spectrometry (HRMS) was performed on a Xevo G3 QTof mass spectrometer (Waters) operated in positive electrospray ionization mode (ESI+). Matrix assisted laser desorption/ionization time-of-flight mass spectrometry (MALDI-TOF MS) was carried out on a Bruker UltraFlextreme. Liquid chromatography electrospray ionization tandem mass spectrometry (LC/ESI-MS/MS) was carried out and processed using a Triple TOF 4600 System (AB Sciex) equipped with a Prominence Ultra-Fast Liquid Chromatography (UFLC) system (Shimadzu). Protein–peptide interactions were quantified by microscale thermophoresis (MST) on a NanoTemper MST instrument. Proteomic identification was performed on a Bruker timsTOF Pro2 ion mobility mass spectrometer using data-dependent acquisition in PASEF mode (DDA-PASEF). The acquired datasets were subsequently imported into TASQ™ for quantitative analysis. Polymerase chain reactions (PCR) were carried out on a C1000 Touch™ thermal cycler (Bio-Rad). DNA sequencing was performed by the Tsingke Biotech company, using appropriate primers. The genome map of *Streptomyces sparsogenes* ATCC 25498 was generated by Proksee[1].

### Methods

**Molecular cloning of precursor peptide and enzyme genes.** Enzyme genes were cloned into pET-24b for overexpression and subsequent *in vitro* assays. Precursor peptide genes together with their cognate enzymes were assembled into pRSFDuet-1 for in vivo co-expression. Target genes were synthesized (LctM and LynD) or amplified from the genomic DNA of *S. sparsogenes* using high fidelity Phanta® DNA Polymerase. PCR amplification was confirmed by 1% agarose gel electrophoresis.

pRSFDuet-1 and pET-24b vectors were digested with appropriate restriction enzymes in 1× NEB buffer (New England Biolabs) at 37 °C for 3 h. PCR products and digested vectors were purified using an Omega Biotech Gel Extraction Kit and subsequently ligated by homologous recombination in 5 × CE II buffer with CE II enzyme at 37 °C for 0.5 h. *E. coli* DH5α cells were transformed with the ligation mixture (10 μL) by heat shock. transformants were selected on LB agar plates supplemented with kanamycin and incubated at 37 °C for 15 h. Single colonies were picked and used to inoculate separate 5 mL LB-kanamycin medium and cultured at 37 °C for 12 h. Plasmids were isolated using an Omega Biotech Plasmid Mini Kit. The sequences of the resulting plasmids were confirmed by DNA sequencing.

**Overexpression and purification of peptides.** *E. coli* BL21 (DE3) cells were transformed with the plasmid described above. A single colony was used to inoculate a 30 mL culture of LB supplemented with 50 μg/mL kanamycin. The culture was grown at 37 °C for 12 h and further used to inoculate three 1 L of LB cultures in 2 L flasks supplemented with 50 μg/mL kanamycin. The culture was grown at 37 °C to $OD_{600}$~0.6-0.8, and cooled at 4 °C on ice for 20 min before the addition of IPTG to a final concentration of 0.2 mM. The culture was grown at 18 °C for additional 18 h. Cells were harvested by centrifugation at 12,000 ×g for 15 min at 4 °C. The resulting cell pellet was resuspended in 100 mL of Buffer-1 (6 M guanidine HCl, 20 mM $NaH_2PO_4$, pH 7.5, 500 mM NaCl, 0.5 mM imidazole) and lysed by sonication on ice. Cell debris was removed by centrifugation at 23,700 ×g for 30 min at 4 °C. The supernatant was filtered through a 0.45 μm filter and loaded to a 5 mL HisTrap HP (GE Healthcare Life Sciences) immobilized metal affinity chromatography (IMAC) column charged with $NiSO_4$ and equilibrated in Buffer-1. The column was washed with five column volumes of Buffer-1, followed by five column volumes of Buffer-2

(4 M guanidine, 20 mM $NaH_2PO_4$, pH 7.5, 500 mM NaCl, 30 mM imidazole). Peptides were eluted with 3 column volumes of Elution Buffer (4 M guanidine, 20 mM $NaH_2PO_4$, pH 7.5, 500 mM NaCl, 1 M imidazole). Fractions containing the desired product were combined, desalted using Sep-Pak® C18 Cartridges and analyzed by MALDI-TOF MS. The product was lyophilized and kept at −80°C for long-term storage.

**Overexpression and purification of enzymes.** Bacterial culture is performed using the same method as above. The pellet was resuspended in 30 mL of start buffer (20 mM Tris buffer, pH 8.0, 500 mM NaCl, 1.0 mM TCEP, 10% glycerol). All protein purification steps were performed at 4 °C. The cell paste was suspended in start buffer, and the cells were lysed using a high-pressure homogenizer (Avestin, Inc.). Cell debris was removed via centrifugation at 23,700 ×g for 20 min at 4 °C. The supernatant was loaded onto a 5 mL HisTrap HP IMAC column charged with $NiSO_4$ and equilibrated with start buffer. The column was washed with 50 mL of buffer A (30 mM imidazole, 20 mM Tris, pH 7.5, 300 mM NaCl), and the protein was eluted using a linear gradient of 0-100% buffer B (200 mM imidazole, 20 mM Tris, pH 7.5, 300 mM NaCl) over 40 min at a 2 mL/min flow rate. Fractions containing proteins were collected and analyzed by SDS-PAGE. Fractions containing target proteins were combined and concentrated using an Amicon Ultra-50 Centrifugal Filter Unit (30 kDa MWCO, Millipore). Eluted fractions containing desired proteins were combined, desalted using PD-10 desalting columns (GE Healthcare) and concentrated by centrifugation. The resulting protein sample was stored at −80 °C. Protein concentration was determined using a Bradford Assay Kit (Pierce).

**Procedure for photo-crosslinking of LctM using LctA leader peptide probes.** LctM (2 μM) and the leader peptide probe (2.5 equiv. relative to the enzyme) were dissolved in 50 mM Tris buffer (pH 8.0) and mixed thoroughly. The mixture was irradiated at 365 nm under 4 °C for 10 min to induce photo-crosslinking. The resulting reaction mixture was purified by gel filtration. *(Note: Excessive irradiation can cause protein precipitation.)*

**Competition assays for the specific photo-crosslinking of LctM by FITC–LctA$_{LP}$–P1 in mixed protein samples.** Purified LctM, BSA and lysozyme were combined in equimolar amounts in 50 mM Tris buffer (pH 8.0) and mixed thoroughly. The mixture was then aliquoted into four equal portions and supplemented with 1, 2, 5 or 10 μM FITC-LctA$_{LP}$-P1 probe. After gentle mixing, samples were irradiated at 365 nm for 10 min at 4 °C. The reaction mixtures were analyzed by 12% SDS-PAGE, followed by in-gel fluorescence imaging and subsequent Coomassie staining.

**Catalytic efficiency of LctM–$LctA_{LP}$ conjugate toward $LctA_{CP}$.** For all LctM-catalyzed reactions, 20 μM $LctA_{cp}$ was incubated with 2 μM purified enzyme (LctM, LctCE, LctM-$LctA_{LP}$-P1 or LctM-$LctA_{LP}$-P2) in the presence of 10 mM $MgCl_2$, 2 mM ATP, 25 μg $mL^{-1}$ bovine serum albumin, and 50 mM Tris buffer (pH 7.5) at 25 °C. Reaction aliquots were collected at 5, 15, 60, and 180 min and quenched by addition of trifluoroacetic acid to a final concentration of 0.5% (pH 1-2). Following centrifugation at high speed for 10 min, clarified supernatants were subjected to LC–MS analysis to quantify the distribution of reaction intermediates and products.

**Procedure for photo-crosslinking of LynD using PatE mini-leader peptide probe.** LynD (2 μM) and PatE mini-leader peptide probe ($PatE_{mLP}$-P1) (10 equiv relative to the enzyme) were dissolved in 50 mM Tris buffer (pH 8.0) and mixed thoroughly. The mixture was irradiated at 365 nm under 4 °C for 10 min to induce photo-crosslinking. The resulting reaction mixture was purified by gel filtration.

**Catalytic efficiency of LynD–$PatE_{mLP}$ conjugate toward $PatE_{CP}$.** For all LynD-catalyzed reactions, 100 μM $PatE_{cp}$ was incubated with 5 μM purified enzyme (LynD or LynD-$PatE_{mLP}$) in the presence of 150 mM NaCl, 10 mM HEPES, pH 7.4, 1 mM TCEP, 5 mM ATP and 5 mM $MgCl_2$ for 16 h at 37 °C. The extent of reaction was monitored by periodically removing aliquots of the reaction, which were quenched by the addition of TFA to 0.5% final concentration (pH 1.0-2.0), desalted by ZipTipC18 and analyzed by MALDI-TOF MS.

**Specific labeling of SpaKC-1 in *E. coli* cell lysates.** An empty pET-28a vector was transformed into *E. coli* BL21 cells for protein expression, and cell lysates were prepared by cell disruption. Purified SpaKC-1 was then spiked into the *E. coli* cell lysates at final contents of 9%, 14%, or 19% (w/w). $SpaA1.2_{LP}$-P1 was subsequently added to each lysate at 1.0, 2.5, or 5.0 equiv relative to SpaKC-1. After gentle mixing, the samples were irradiated at 365 nm for 10 min at 4 °C. Reaction mixtures were analyzed by 8% SDS-PAGE, followed by in-gel fluorescence imaging and subsequent Coomassie staining.

**Chemical proteomics for target identification.**

**Step 1: Preparation of cell lysate.** *Streptomyces sparsogenes* ATCC 25498 was cultured by fermentation for 5 days (100 mL). Cells were harvested and washed twice with phosphate-buffered saline (PBS), followed by a single wash with Tris buffer (50 mM Tris, 75 mM NaCl, pH 8.0). The cell pellet from the 100 mL culture was resuspended in 10 mL of Tris buffer with gentle agitation. Cells were lysed by ultrasonication at 65 W for 90 min using a pulse program of 3 s on and 6 s off. The lysate was clarified by centrifugation at 12 000 rpm for 20 min, and the resulting supernatant was collected as the crude cell lysate. Total protein concentration was determined using a BCA assay.

**Step 2: Photo-crosslinking of cell lysates with SpaA1.2$_{LP}$-P2**. Cell lysates were supplemented with SpaA1.2$_{LP}$-P2 probe at final concentrations of 0, 1, 5, or 10 μM. The samples were then subjected to UV irradiation at 365 nm (30 W) for 10 min at 4 °C to induce photo-crosslinking. A parallel control sample lacking UV irradiation was processed under otherwise identical conditions.

**Step 3: Protein precipitation**. Following the photo-crosslinking reaction, five volumes of cold methanol were added to each sample, and the mixtures were incubated at -20 °C for 4 h to allow complete protein precipitation. The precipitated proteins were collected by centrifugation at 21500 ×g for 10 min, and the supernatants were carefully removed. The methanol precipitation and washing procedure was repeated three times to ensure thorough removal of unreacted SpaA1.2$_{LP}$-P2 probe. The resulting protein pellets were air-dried to remove residual methanol.

**Step 4: Blocking of Streptavidin–agarose beads**. For each sample, 200 μL of streptavidin–agarose beads were washed with 10 mL of PBS (pH 7.4) by gentle agitation at 4 °C for 5 min, followed by centrifugation at 1000 ×g for 1 min. This washing step was repeated three times. The beads were then incubated with 1.8 mL of BSA blocking solution at 4 °C for 4 h with gentle agitation to achieve complete surface passivation. The BSA blocking solution consisted of 5 mg $mL^{-1}$ bovine serum albumin dissolved in a mixture of 4% SDS and 10 mM EDTA (pH 7.4), supplemented with 1% Brij 97, 150 mM NaCl, and 50 mM triethanolamine (pH 7.4).

**Step 5: Protein resolubilization**. Protein pellets were resuspended in a total volume of 4 mL of resolubilization buffer composed of a 1:8 (v/v) mixture of 4% SDS and 10 mM EDTA (pH 7.4) with 1% Brij 97, 150 mM NaCl, and 50 mM triethanolamine (pH 7.4). The suspensions were agitated until complete protein resolubilization was achieved.

**Step 6: Biotin–streptavidin enrichment.** The blocked streptavidin-agarose beads were washed three times with PBS (pH 7.4) and collected by centrifugation at 1000 ×g for 1 min after each wash. The resolubilized protein samples were then added to the beads, and the mixtures were incubated at 4 °C for 12 h with gentle agitation to allow biotin–streptavidin binding.

**Step 7: Removal of nonspecifically bound proteins.** Following enrichment, the streptavidin-agarose beads were pelleted by centrifugation at 1000 ×g for 1 min, and the supernatant was carefully removed. The beads were then sequentially washed with the following solutions to remove nonspecifically bound proteins: (a) 2% SDS in PBS (pH 7.4); (b) 8 M urea in 250 mM ammonium bicarbonate; (c) 2.5 M NaCl in PBS (pH 7.4); (d) 0.5 M ammonium bicarbonate; (e) 0.25 M ammonium bicarbonate; and (f) 0.05 M ammonium bicarbonate. Each wash was performed at room temperature with gentle agitation for 5 min, followed by centrifugation at 1000 ×g for 1 min and removal of the supernatant.

**Step 8: Protein elution.** The enriched streptavidin–agarose beads (200 μL) were divided into two equal portions. One portion was resuspended in SDS–PAGE loading buffer and heated at 100 °C for 10 min, after which the eluate was subjected to pull-down analysis. The second portion was incubated with elution buffer consisting of 8 M guanidine hydrochloride (pH 1.5) and heated at 95 °C for 5 min. This elution step was repeated three times, and the supernatants were collected by centrifugation. The combined eluates were subsequently concentrated using an ultrafiltration device and processed for downstream proteolytic digestion.

**Step 9: Proteolytic digestion.** The concentrated protein samples were adjusted to pH 8.0 using 8 M urea in 250 mM ammonium bicarbonate. Disulfide bonds were reduced with dithiothreitol, followed by alkylation with iodoacetamide. The buffer was subsequently exchanged to 50 mM triethylammonium bicarbonate (TEAB), and on-filter proteolytic digestion was performed using trypsin at an enzyme-to-substrate ratio of 1:30 (w/w) at 37 °C for 16 h. The resulting peptides were collected, desalted, and subjected to LC–MS/MS analysis for protein identification.

**Procedure for photo-crosslinking of SpaKC-1, SpaKC-2 and SpaL using SpaA1.2$_{LP}$-P3.** Enzyme (2 μM) and SpaA1.2$_{LP}$-P3 (2.5 equivalents relative to the enzyme) were dissolved in 50 mM Tris buffer (pH 8.0) and mixed thoroughly. The mixture was irradiated at 365 nm under 4 °C for 10 min to induce photo-crosslinking.

**Procedures of *in vitro* modification assays and derivatization assays.**

***In vitro* modification of the precursor peptides by lanthipeptide synthetases.** Typically, the precursor peptide (50 μM) and lanthipeptide synthetase (10 μM) were incubated with 0.5 mM TCEP, 2.5 mM ATP and 5 mM $MgCl_2$ in 20 mM Tris-HCl, pH 8.0, at 28 °C for 4 h. The reaction was quenched by adding 0.1% HCOOH (final concentration) and then centrifuged at 14,000 ×g for 5 min. The supernatant of the reaction mixture was desalted by a SPE column before analysis by MALDI-TOF-MS and LC-MS/MS.

**Derivatization of free cysteine residues in peptides with NEM.** Peptides (50 μM) were typically modified by 5 mM N-ethylmaleimide (NEM) in 20 mM Tris-HCl (pH 8.0) and 0.5 mM TCEP at 37 °C in the dark for 0.5 h. The reaction was then quenched by the addition of 20 mM DTT. The reaction mixture was desalted by Ziptip and further analyzed by MALDI-TOF-MS.

**Derivatization of Dha/Dhb residues in peptides with βME.** Peptides (50 μM) were typically treated by β-mercaptoethanol (β-Me, 10 mM) in 20 mM Tris-HCl (pH 8.0) at 37 °C for 1 h. The reaction mixture was desalted by Ziptip and further analyzed by MALDI-TOF-MS.

**Procedure for MALDI-TOF MS analysis of the endopeptidase activity of SpaP1 and SpaP2.** 100 μM peptide substrate, 20 μM enzyme in 20 mM Tris-HCl buffer (pH 8.0), incubated at 37°C for 24 h. Reactions were quenched with 0.1% HCOOH (final concentration) and centrifuged; the supernatant was subjected to MALDI-TOF analysis.

**Procedure for the aminopeptidase activity analysis of SpaP1 and SpaP2 by the hydrolysis of amino acid-*para*-nitroanilide (*p*NA) derivatives.** 100 μM *p*NA derivative, 1 μM enzyme in 20 mM Tris-HCl buffer (pH 8.0), incubated at 37°C. Absorbance at 405 nm was recorded every 30 s using a UV-Vis spectrophotometer.

**Procedure of RNA extraction, reverse transcription and qRT-PCR**. Mycelia of *S. sparsogenes* grown in YEME were collected, frozen in liquid nitrogen, and ground to a fine powder. The total mRNA was extracted using Trizol reagent (Vazyme) according to the manufacturer's instructions. Isolated mRNA was reverse-transcribed into cDNA using a cDNA synthesis kit (Vazyme). The qRT-PCR was performed on the Bio-Rad C1000 Touch Thermal Cycler + CFX96 Real Time System Lab using synthetic primers and ChamQ Blue Universal SYBR qPCR Master Mix (Vazyme). The 16S rRNA gene was used as internal control. Relative expression level was calculated by the comparative Ct method. Results were normalized relative to 16S rRNA expression. Each experiment was performed in triplicate.**Chemical synthesis of leader peptide probes**

Peptides were assembled on an automated microwave peptide synthesizer (Liberty Blue$^{TM}$, CEM) using Rink MBHA Amide. Semi-preparative HPLC was performed on a Shimadzu system equipped with an LC-20AR semi-preparative solvent delivery unit and an SPD-20A UV/Vis detector. The mobile phase consisted of buffer A ($H_2O$ with 0.1% formic acid) and buffer B ($CH_3CN$ with 0.1% formic acid). Analytical HPLC was conducted using an ACQUITY Premier Peptide BEH C18 column (1.7 μm, 2.1 × 150 mm, Waters) at a flow rate of 0.4 mL/min with a 2489 UV/Vis detector. All water used in the experiments was distilled and filtered through a MilliporeSigma purification system. LC–MS analysis was performed using a Welch Materials Ultimate® XB-C18 column (5 μm, 10 × 250 mm) with PDA detection and a linear gradient from 10% to 90% acetonitrile containing 0.1% formic acid.

**General procedures for leader peptide probes synthesis:**

**a. Fmoc deprotection.** The Fmoc group was removed using 20% piperidine in DMF (2 x 10 min) or by microwave irradiation at 90 °C for 1 min. The resin was extensively washed with *N,N*-dimethylformamide (DMF, 3 x).

**b. Amino acid coupling.** Fmoc amino acid (5.0 equiv relative to resin loading), ethyl(hydroxyimino)cyanoacetate (oxyma, 5.0 equiv), and N,N′-diisopropylcarbodiimide (DIC, 5.5 equiv) were dissolved in DMF, and the resulting mixture was added to the resin. Coupling was performed either by microwave irradiation at 90 °C for 2 min or by shaking at 25 °C for 1 h, after which the resin was washed with DMF (3 x) before proceeding to the next deprotection cycle.

**c. Capping of the N-terminus with Ac:** N,N-Diisopropylethylamine (DIPEA, 10 equiv) and acetic anhydride (10 equiv) were added to the resin, and the mixture was shaken at 25 °C for 1 h. The resin was then washed with DMF (3 x) followed by DCM (3 x).

**d. Dde deprotection**. The Dde protecting group was removed using 5% hydrazine monohydrate in DMF (v/v) (2 x 10 min). The resin was then washed thoroughly with DMF (3 x) followed by DCM (3 x).

**e. Coupling with 4-(hydroxymethyl)-3-nitrobenzoic acid**. 4-(Hydroxymethyl)-3-nitrobenzoic acid (3.0 equiv), oxyma (3.0 equiv), and DIC (3.3 equiv) were dissolved in N-methylpyrrolidone (NMP), and the resulting mixture was added to the resin. Coupling was carried out by shaking at 25 °C for 3 h, after which the resin was washed with DMF (3 x) followed by DCM (3 x).

**f. Loading of 2-Cl-Trt resin**. Fmoc-AA-OH (2 equiv) and DIPEA (7 equiv) were dissolved in DCM and added to the resin, and the mixture was agitated at room temperature for 2 h.

**g. Cleavage from the resin**. The dried Rink Amide MBHA or 2-Cl-Trt resin was treated with a cleavage cocktail of TFA/TIPS/$H_2O$ (95:2.5:2.5, v/v/v) for 2 h (or 4 h for peptides longer than 15 amino acids). Volatile solvents were removed under a stream of $N_2$ or Ar, and the resulting residue was triturated with cold diethyl ether to precipitate the crude peptide. After centrifugation, the supernatant was discarded, and the crude peptide was purified by semi-preparative HPLC.

**h. Fluorescein 5-isothiocyanate (FITC) and primary amines substitution reaction.** FITC (1.2 equiv) and the peptide substrate (1.0 equiv) were dissolved in DMF to a final concentration of 50 μM, followed by the addition of DIPEA (10 equiv). The reaction mixture was stirred at room temperature for 1 h, concentrated under reduced pressure, and purified by semi-preparative HPLC.

**i. NHS ester–amine substitution reaction.** The NHS ester (10 equiv) and peptide substrate (1.0 equiv) were dissolved in PB buffer (pH 8.0) to a final concentration of 50 μM. The reaction mixture was stirred at 37 °C for 4 h and subsequently purified by semi-preparative HPLC.

### Synthetic routes to leader peptide probes

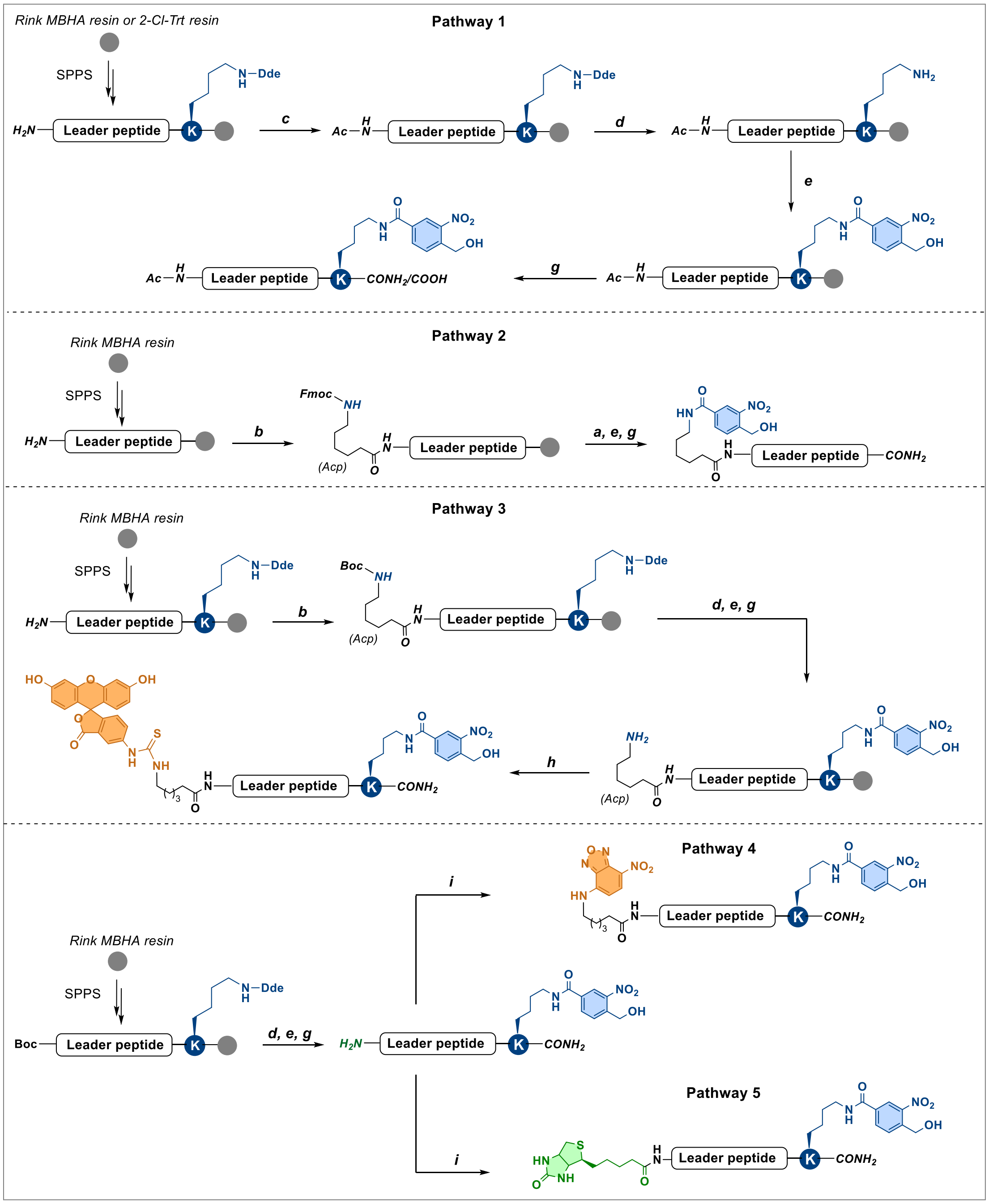

**Scheme S1.** Synthetic routes to the leader peptide probe in this study.

**Pathway 1.** C-terminal NBA leader peptide probes were assembled on either Rink amide MBHA resin (yielding C-terminal amides) or 2-Cl-Trt resin (yielding C-terminal acids) on a 0.1 mmol scale. Fmoc-Lys(Dde)-OH was first coupled to the resin, followed by standard Fmoc solid-phase peptide elongation as described in General Procedures **a** and **b**. After chain assembly, the N terminus was acetylated according to General Procedure ***c***. The Dde protecting group was then removed (General Procedure **d**), and the peptide was coupled with 4-(hydroxymethyl)-3-

nitrobenzoic acid (General Procedure **e**). Finally, the peptides were cleaved from the resin following General Procedure **g** and purified by semi-preparative HPLC.

**Pathway 2.** N-terminal NBA leader peptide probes were synthesized on Rink amide MBHA resin (0.1 mmol scale) by standard Fmoc solid-phase peptide synthesis as described in General Procedures **a** and **b**. Fmoc-6-aminohexanoic acid (Fmoc-Acp-OH) was introduced at the N-terminus as a spacer. The Fmoc group was removed and 4-(hydroxymethyl)-3-nitrobenzoic acid was then coupled (General Procedure **a** and **e**). Peptides were cleaved from the resin according to General Procedure **g**, then purified by semi-preparative HPLC.

**Pathway 3.** Leader peptide probes bearing an N-terminal FITC group and a C-terminal NBA group were synthesized on Rink amide MBHA resin on a 0.1 mmol scale. Fmoc-Lys(Dde)-OH was installed as the initial amino acid, followed by standard Fmoc solid-phase peptide synthesis according to General Procedures **a** and **b**. Boc-6-aminohexanoic acid (Boc-Acp-OH) was incorporated as the final amino acid for orthogonal protection. After removal of the Dde group, the resin was coupled with 4-(hydroxymethyl)-3-nitrobenzoic acid (General Procedure **d** and **e**), and the peptides were cleaved from the resin following Procedure **g** and purified by semi-preparative HPLC. The purified peptides were subsequently reacted with FITC to introduce the N-terminal fluorescent label (General Procedure **h**).

**Pathway 4.** Leader peptide probes bearing an N-terminal NBD group and a C-terminal NBA group were synthesized on Rink amide MBHA resin (0.1 mmol scale). Fmoc-Lys(Dde)-OH was installed as the first amino acid, followed by standard Fmoc solid-phase peptide synthesis according to General Procedures **a** and **b**. Boc-AA-OH was incorporated as the C-terminal residue for orthogonal protection. After Dde removal, the resin was coupled with 4-(hydroxymethyl)-3-nitrobenzoic acid (General Procedure **d** and **e**). Peptides were then cleaved from the resin following Procedure **g** and purified by semi-preparative HPLC. The purified peptides were finally reacted with NBD-NHS ester to introduce the N-terminal fluorescent label (General Procedure **i**).

**Pathway 5.** Leader peptide probes bearing an N-terminal biotin group and a C-terminal NBA group were synthesized on Rink amide MBHA resin (0.1 mmol scale). Fmoc-Lys(Dde)-OH was installed as the first amino acid, followed by standard Fmoc solid-phase peptide synthesis according to General Procedures **a** and **b**. Boc-AA-OH was incorporated as the C-terminal residue for orthogonal protection. After Dde removal, the resin was coupled with 4-(hydroxymethyl)-3-nitrobenzoic acid (General Procedure **d** and **e**). The peptides were then cleaved from the resin following Procedure **g** and purified by semi-preparative HPLC. The purified peptides were finally reacted with biotin-NHS ester to install the N-terminal biotin moiety (General Procedure **i**).

### Supplementary Tables

**Table S1.** Primers for molecular cloning

| Name | Sequence |
|---|---|
| pET24b-SpaKC-1-F | GAAGGAGATATACATATGAGGAAACCGTACGGATTCACTGTCGCC |
| pET24b-SpaKC-1-R | AAAGAGGACTTCAAGGGACCGTGCGGCGATCGC |
| pET24b-SpaKC-2-F | TAAGAAGGAGATATACATATGATGGACAAAAGGTATGAGGTCTTCTG |
| pET24b-SpaKC-2-R | GGGTCCCTGAAAGAGGACCTCGAGATTTCCCTGGGGATGGG |
| pET24b-SpaL-F | TAAGAAGGAGATATACATATGGTGCTCACTTTCCCGTTGCA |
| pET24b-SpaL-R | GTGGTGGTGGTGGTGCTCGAGGGACAGCGGCATCCA |
| pRSF-SpaA1.1-F | CACCACAGCCAGGATCCGATGAGCGGAGTTCTGGCCCT |
| pRSF-SpaA1.1-R | ATTATGCGGCCGCAAGCTTCAGCCCTGAGGCATCGCC |
| pRSF-SpaA1.2-F | ACCACAGCCAGGATCCGTTGCACCAGATTCTGAAACTGCAGGA |
| pRSF-SpaA1.2-R | TATGCGGCCGCAAGCTCTAGTGGGTGAAGATCGAGGGCG |
| pRSF-SpaA1.3-F | TCATCACCACAGCCAGGATCCAATGCCGCAGATCCTGGAAC |
| pRSF-SpaA1.3-R | GCATTATGCGGCCGCAAGCTTCTAGCAGACGGCGCTGAAGT |
| pRSF-SpaA$_L$-F | TCATCACCACAGCCAGGATCCAATGGACACCGACCTCGACG |
| pRSF-SpaA$_L$-R | GCATTATGCGGCCGCAAGCTTTCAGACGTCCGTACGCCCG |

**Table S2.** Primers for qRT-PCR

| Name | Sequence |
|---|---|
| SpaKC-1-F | ACTTCCGGGAATGGGAACAC |
| SpaKC-1-R | TTCTGATGCAGGGGTTGTGG |
| SpaKC-2-F | CAAGGTCTGGGACTACTGCG |
| SpaKC-2-R | GGGTAGATGGTGGCGAACTT |
| SpaL-F | GTGCTGATAGACCCGGAGTG |
| SpaL-R | GAGCGAGTAGAGGTCCGACT |
| SpaP1-F | ACGACGTCCTGGTCAACTTC |
| SpaP1-R | TGGACGCCCTTGAAGAACTC |
| SpaP2-F | GGAAGTACGACCAGGCCTTC |
| SpaP2-R | TCGTGGAGGATCACATTGGC |
| 16S rRNA-F | GGCATCTTACCGGGTGGAAA |
| 16S rRNA-R | GTGTCTCAGTCCCAGTGTGG |

**Table S3.** Relevant modification enzymes and precursor peptides in *Streptomyces sparsogenes* ATCC 25498.

| | | |
|---|---|---|
| | ***Lan III-1* cluster** | |
| **Name** | **NCBI Accession** | **Description** |
| SpaA1.1 | WP_158080302.1 | Class III lanthipeptide precursor peptide |
| SpaA1.2 | WP_281182485.1 | Class III lanthipeptide precursor peptide |
| SpaA1.3 | allorf_32520_32657 | Class III lanthipeptide precursor peptide |
| SpaKC-1 | WP_065963996.1 | Class III lanthipeptide synthetase |
| | ***Lan III-2* cluster** | |
| **Name** | **NCBI Accession** | **Description** |
| SpaA2 | WP_104531210.1 | Class III lanthipeptide precursor peptide |
| SpaKC-2 | WP_065958791.1 | Class III lanthipeptide synthetase |
| | ***Lan IV* cluster** | |
| **Name** | **NCBI Accession** | **Description** |
| SpaA$_{L}$ | WP_211301783.1 | Class IV lanthipeptide precursor peptide |
| SpaL | WP_065963992.1 | Class IV lanthipeptide synthetase |
| | ***Tat* cluster** | |
| **Name** | **NCBI Accession** | **Description** |
| SpaA | WP_158080408.1 | Thioviridamide family precursor peptide |
| SpaC | WP_065960364.1 | Phosphotransferase |
| SpaD | WP_065960366.1 | Lyase |
| SpaE | WP_065960368.1 | Pseudokinase |
| SpaF | WP_065960370.1 | Flavoprotein |
| | **Peptidases beyond BGCs** | |
| **Name** | **NCBI Accession** | **Description** |
| SpaP1 | WP_065959395.1 | M1 family metallopeptidase |
| SpaP2 | WP_065959395.1 | M1 family metallopeptidase |

**Table S4.** Sequence alignment of relevant precursor peptides in *Streptomyces sparsogenes* ATCC 25498. Conserved recognition sequences within the N-terminal leader peptides are underlined.

| BGC | Name | Sequence |
|---|---|---|
| *Lan III-1* | SpaA1.1 | MSGVLALQALEPADTEVAD-ALPTTMTITTGNSHLSVFMDCTTTVTRAMPQG |
| | SpaA1.2 | MHQILKLQELASEVTSEEE-GVVEVSTPSVLLCIPPSIFTH |
| | SpaA1.3 | MPQILELQELENSHGPDED-GVEEWSTLSGVCQSDFSAVC |
| *Lan III-2* | SpaA2 | MTLLDLQTLETEESESFGG-GGGGESSVSLLLCDKNSAVSQLLCL |
| *Lan IV* | SpaA$_L$ | MDTDLDALQLLPSEEE-TAAICWFTCLGTCGGVTCDGTCGRTDV |

### Supplementary Figures

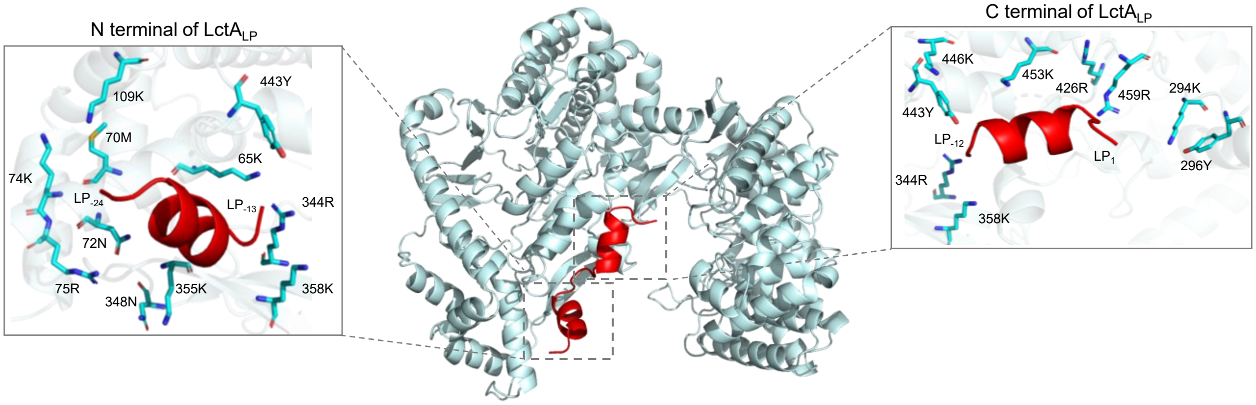

**Figure S1.** The LctM–LctALP docking model reveals the presence of potential photo-crosslinking residues (Lys, Arg, and Tyr) in close proximity to the leader peptide–binding site (LctM, cyan; LctALP, red).

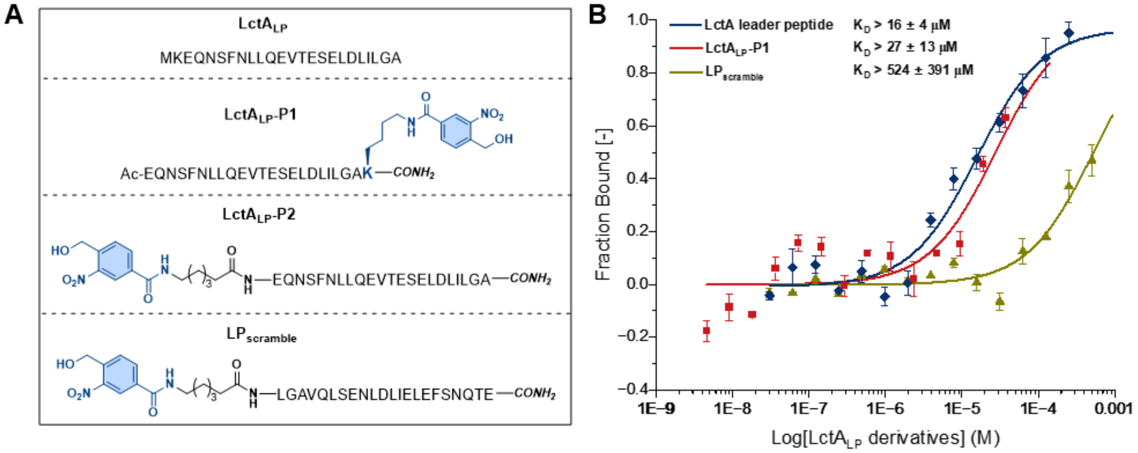

**Figure S2.** (A) Structure of LctALP, LctALP-P1, LctALP-P2 and LPscramble. (B) MST analysis of the binding affinities between LctM and leader peptide derivatives. Error bars represent the SD of the mean (n = 3). The results indicate that installation of the NBA group does not disrupt the interaction between the leader peptides and LctM. The scrambled probe shows negligible binding with LctM. Due to the low solubility of LctALP-P2 in PBS-T (<0.1 mg/mL), its MST affinity could not be accurately determined.

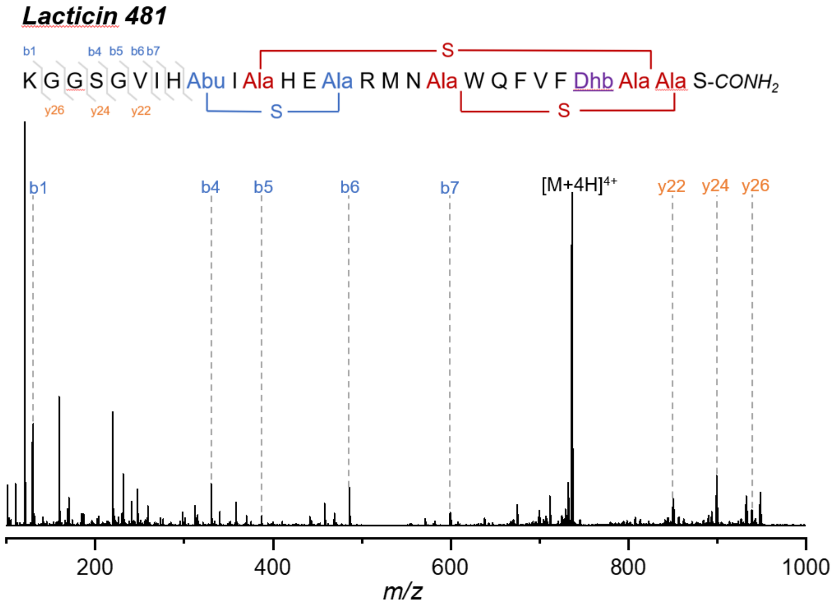

| Species | Calc. mass. (Da) | Obs. mass. (Da) | Err. (ppm) |
|---|---|---|---|
| b7 | 599.3512 | 599.3505 | -1.2 |
| b6 | 486.2671 | 486.267 | -0.2 |
| b5 | 387.1987 | 387.199 | 0.8 |
| b4 | 330.1772 | 330.1784 | 3.6 |
| b1 | 129.1023 | 129.1025 | 1.5 |
| y26 | 938.4154 | 938.4212 | 6.2 |
| y24 | 900.4011 | 900.4013 | 0.2 |
| y22 | 852.3833 | 852.3809 | -2.8 |

**Figure S3.** MS/MS validation of LctA$_{cp}$ cyclization sites catalyzed by LctM-LctA$_{LP\text{-}P1}$.

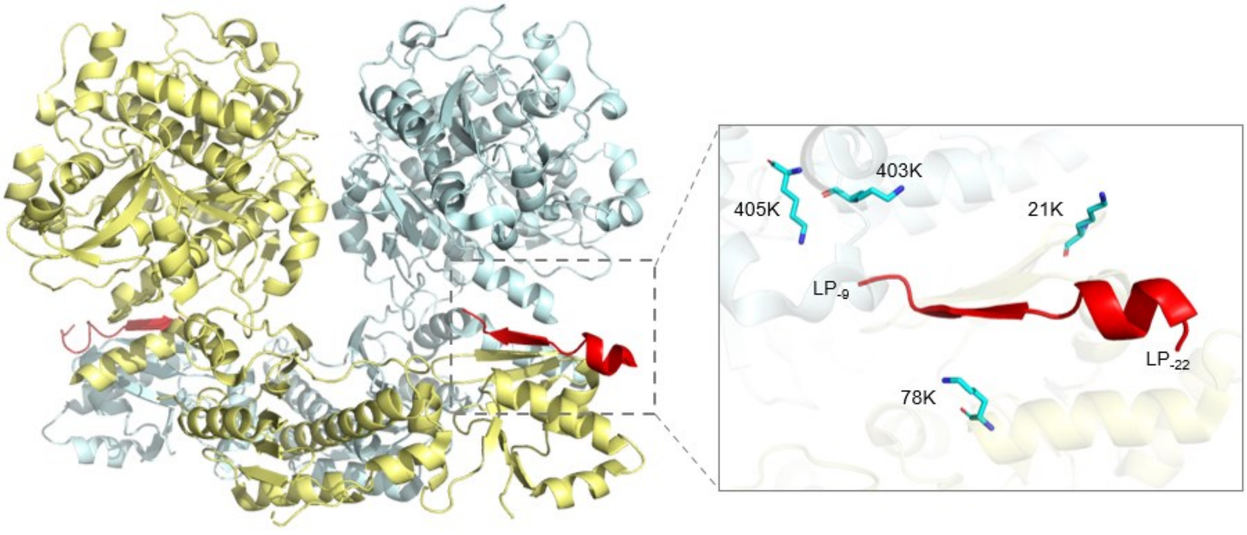

**Figure S4.** LynD-PatE$_{mLP}$ model indicates that the C-terminal region of the leader peptide inserts into a defined binding pocket containing residues suitable for *o*-NBA-mediated photochemistry. Monomers colored pale yellow and cyan. PatE minimal leader peptide (mLP) colored red.

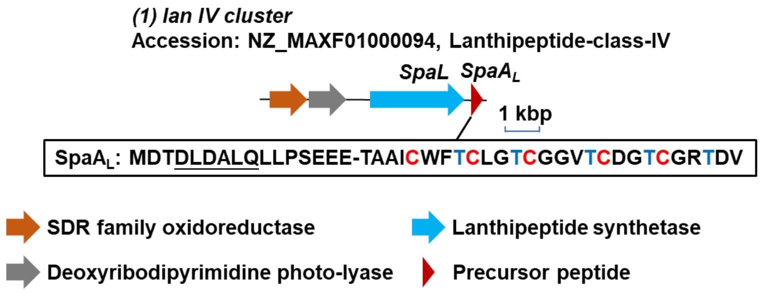

| Relevant precursors and proteins | NCBI Accession | Putative Function |
|---|---|---|
| SpaSDR | WP_079151080.1 | SDR family oxidoreductase |
| SpaDPL | WP_065963990.1 | Deoxyribodipyrimidine photo-lyase |
| SpaL | WP_065963992.1 | Class IV lanthipeptide synthetase |
| $SpaA_L$ | WP_211301783.1 | Class IV lanthipeptide precursor peptide |

**Figure S5.** The *lan IV* cluster and relevant precursors and proteins within the BGC. The precursor peptide sequence is highlighted in the box. AntiSMASH [2] was used to analyze BGC.

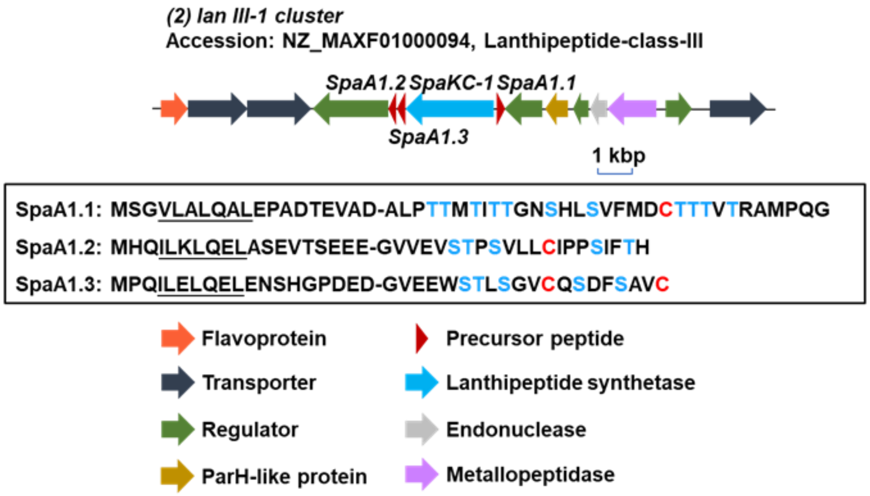

| Relevant precursors and proteins | NCBI Accession | Putative Function |
|---|---|---|
| SpaD1 | WP_065963994.1 | Flavoprotein |
| SpaT1 | WP_141713022.1 | ABC transporter |
| SpaT2 | WP_158080303.1 | ABC transporter |
| SpaR1 | WP_065963995.1 | LuxR family transcriptional regulator |
| SpaA1.2 | WP_281182485.1 | Class III lanthipeptide precursor peptide |
| SpaA1.3 | - | Class III lanthipeptide precursor peptide |
| SpaKC-1 | WP_065963996.1 | Class III lanthipeptide synthetase |
| SpaA1.1 | WP_158080302.1 | Class III lanthipeptide precursor peptide |
| SpaT3 | WP_065964004.1 | MAB_1171c family putative transporter |
| SpaPH | WP_065964005.1 | ParH-like protein |
| SpaR2 | WP_065964006.1 | Helix-turn-helix transcriptional regulator |
| SpaEN | WP_065964008.1 | Uma2 family endonuclease |
| SpaP | WP_065964080.1 | M1 family metallopeptidase |
| SpaR3 | WP_065964009.1 | GntR family transcriptional regulator |
| SpaT4 | WP_065964011.1 | APC family permease |

**Figure S6.** The *lan III-1* cluster and relevant precursors and proteins within the BGC. The precursor peptide sequence is highlighted in the box.

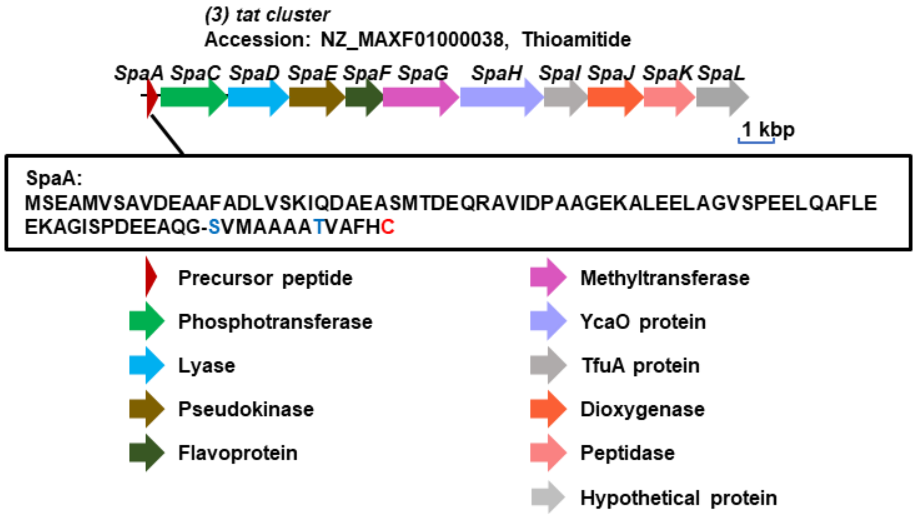

| Relevant precursors and proteins | NCBI Accession | Putative Function |
|---|---|---|
| SpaA | WP_158080408.1 | Thioviridamide family precursor peptide |
| SpaC | WP_065960364.1 | Phosphotransferase |
| SpaD | WP_065960366.1 | Lyase |
| SpaE | WP_065960368.1 | Pseudokinase |
| SpaF | WP_065960370.1 | Flavoprotein |
| SpaG | WP_065960373.1 | 50S ribosomal protein L11 methyltransferase |
| SpaH | WP_065960374.1 | YcaO protein |
| SpaI | WP_065960379.1 | TfuA protein |
| SpaJ | WP_065960381.1 | Phytanoyl-CoA dioxygenase |
| SpaK | WP_065960382.1 | C1 family peptidase |
| SpaL | WP_245737963.1 | Hypothetical protein |

**Figure S7.** The *tat* cluster [3] and relevant precursors and proteins within the BGC. The precursor peptide sequence is highlighted in the box.

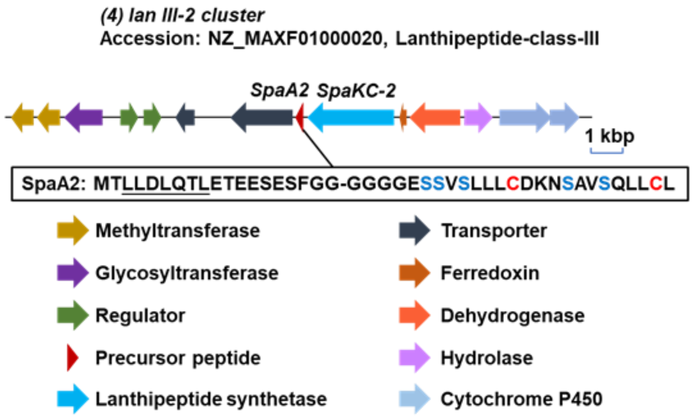

| Relevant precursors and proteins | NCBI Accession | Putative Function |
|---|---|---|
| SpaPT | WP_065958787.1 | Prenyltransferase |
| SpaMT | WP_065958788.1 | Class I SAM-dependent methyltransferase |
| SpaGT | WP_425428765.1 | Glycosyltransferase family 4 protein |
| SpaR4 | WP_065958832.1 | TetR family transcriptional regulator |
| SpaR5 | WP_079150775.1 | Response regulator transcription factor |
| SpaT5 | WP_065958789.1 | ABC transporter |
| SpaT6 | WP_065958790.1 | ABC transporter |
| SpaA2 | WP_104531210.1 | Class III lanthipeptide precursor peptide |
| SpaKC-2 | WP_065958791.1 | Class III lanthipeptide synthetase |
| SpaFR | WP_065958792.1 | Ferredoxin |
| SpaAD | WP_065958793.1 | Aldehyde dehydrogenase |
| SpaMH | WP_065958794.1 | MBL fold metallo-hydrolase |
| SpaP450-1 | WP_245737868.1 | Cytochrome P450 |
| SpaP450-2 | WP_245737876.1 | Cytochrome P450 |

**Figure S8.** The *lan III-2* cluster and relevant precursors and proteins within the BGC. The precursor peptide sequence is highlighted in the box.

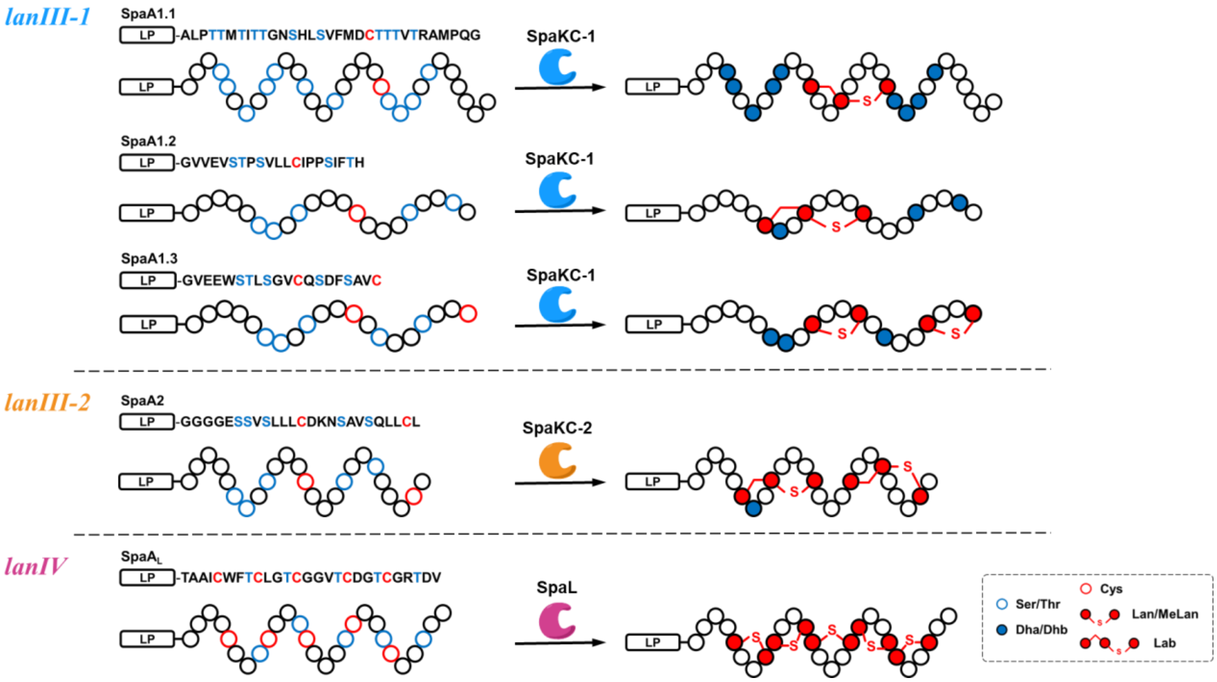

**Figure S9.** Precursor peptides processed by enzymes encoded within cognate BGCs.

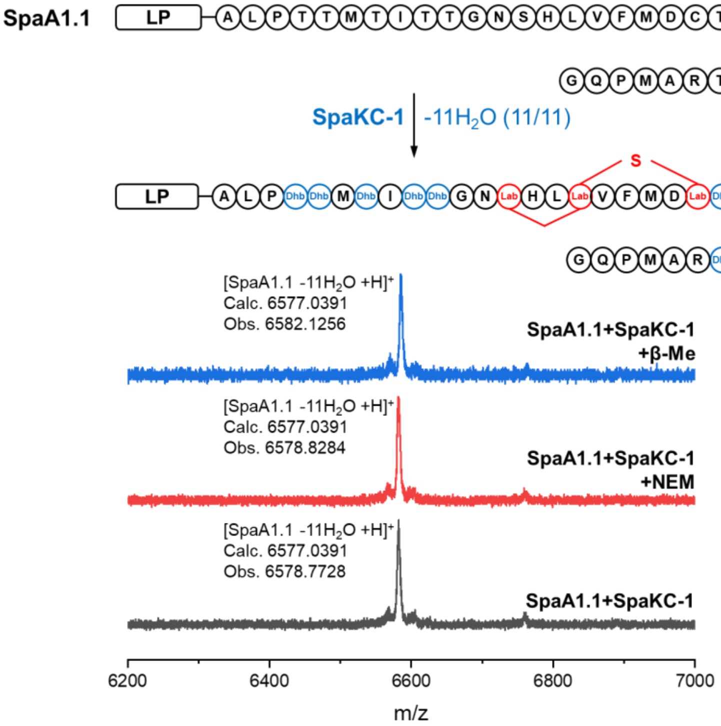

**Figure S10.** MALDI-TOF MS analysis and NEM/β-Me derivatization of product from *in vivo* coexpression of SpaKC-1 and precursor peptide SpaA1.1.

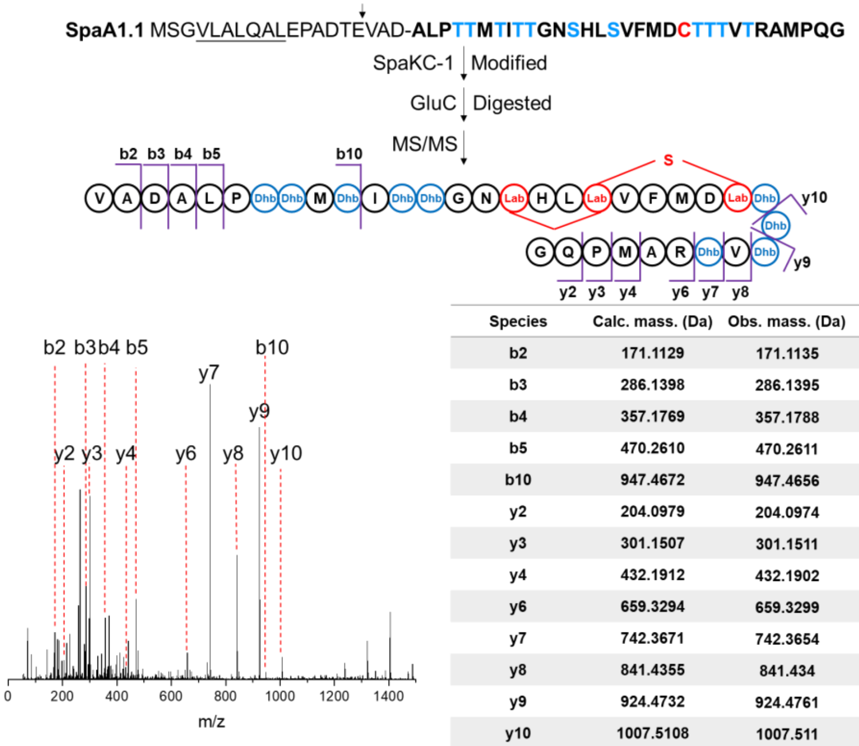

| Species | Calc. mass. (Da) | Obs. mass. (Da) | Err. (ppm) |
|---|---|---|---|
| b2 | 171.1129 | 171.1135 | 3.5 |
| b3 | 286.1398 | 286.1395 | -1.0 |
| b4 | 357.1769 | 357.1788 | 5.3 |
| b5 | 470.2610 | 470.2611 | 0.2 |
| b10 | 947.4672 | 947.4656 | -1.7 |
| y2 | 204.0979 | 204.0974 | -2.4 |
| y3 | 301.1507 | 301.1511 | 1.3 |
| y4 | 432.1912 | 432.1902 | -2.3 |
| y6 | 659.3294 | 659.3299 | 0.8 |
| y7 | 742.3671 | 742.3654 | -2.3 |
| y8 | 841.4355 | 841.434 | -1.8 |
| y9 | 924.4732 | 924.4761 | 3.1 |
| y10 | 1007.5108 | 1007.511 | 0.4 |

**Figure S11.** Structural characterization via MS/MS of the SpaKC-1-modified SpaA1.1 product. Peptide products were digested with GluC before MS/MS to remove the N-terminal segments. The cleavage site is indicated by an arrow in the sequence.

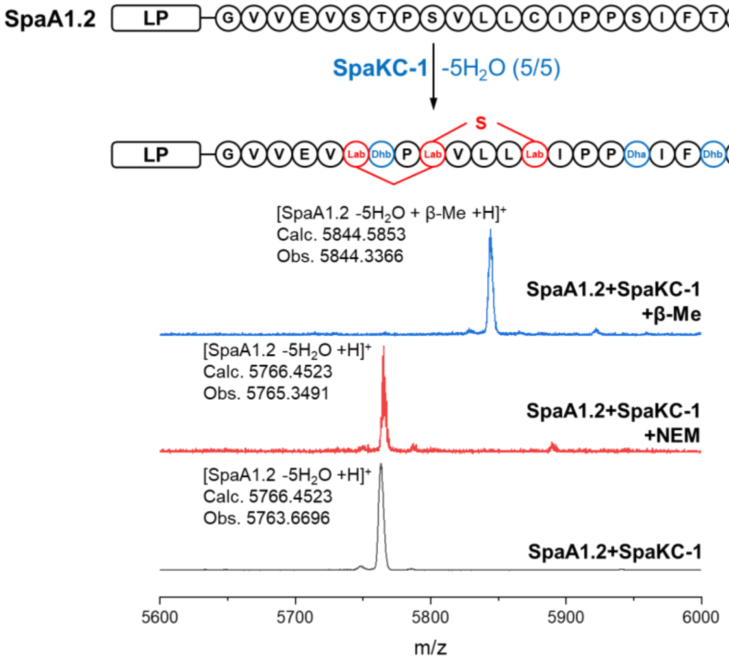

**Figure S12.** MALDI-TOF MS analysis and NEM/β-Me derivatization of product from *in vitro* modification of SpaKC-1 and precursor peptide SpaA1.2.

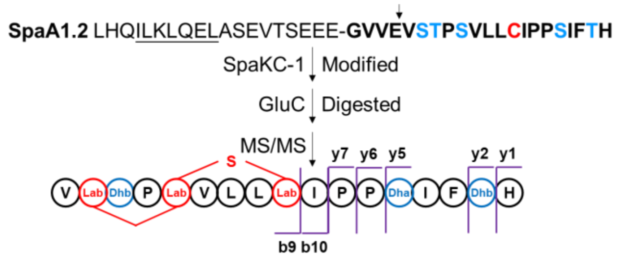

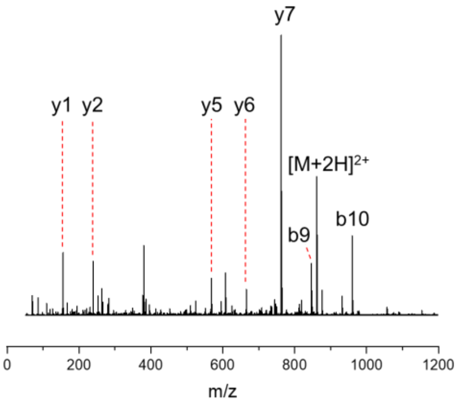

| Species | Calc. mass. (Da) | Obs. mass. (Da) | Err. (ppm) |
|---|---|---|---|
| b9 | 846.4560 | 846.4531 | -3.4 |
| b10 | 959.5400 | 959.5371 | -3.0 |
| y1 | 156.0768 | 156.0771 | 1.9 |
| y2 | 239.1145 | 239.1137 | -3.3 |
| y5 | 568.2890 | 568.2866 | -4.2 |
| y6 | 665.3417 | 665.3398 | -2.9 |
| y7 | 762.3945 | 762.3923 | -2.9 |
| [M+2H]$^{2+}$ | 860.9673 | 860.9663 | -1.2 |

**Figure S13.** Structural characterization via MS/MS of the SpaKC-1-modified SpaA1.2 product.

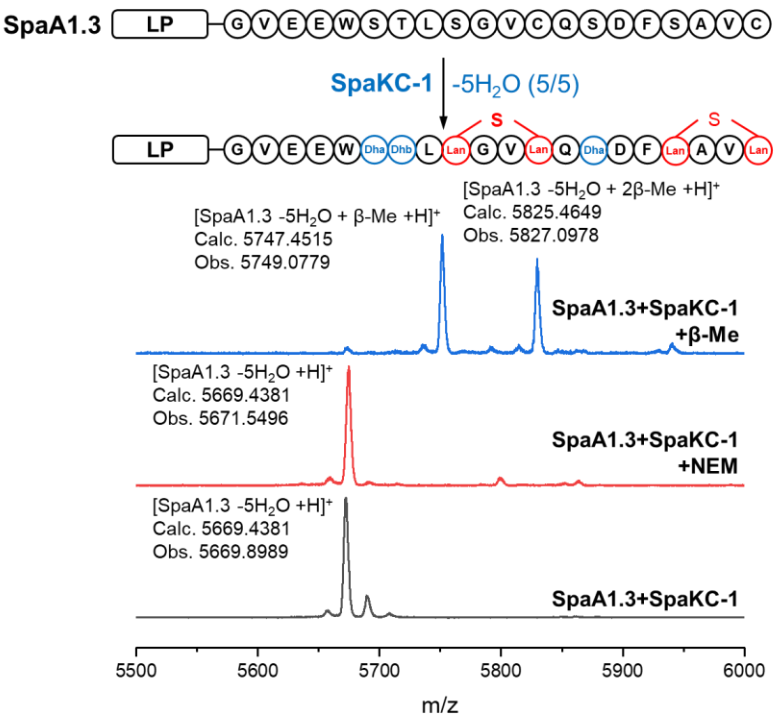

**Figure S14.** MALDI-TOF MS analysis and NEM/β-Me derivatization of product from *in vitro* modification of SpaKC-1 and precursor peptide SpaA1.3.

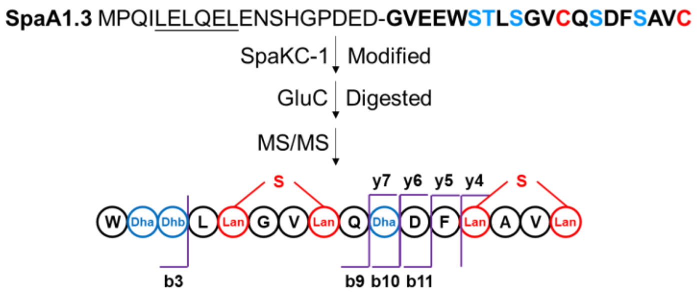

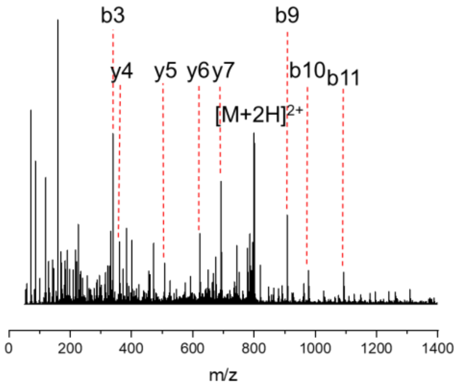

| Species | Calc. mass. (Da) | Obs. mass. (Da) | Err. (ppm) |
|---|---|---|---|
| b3 | 339.1463 | 339.1449 | -4.1 |
| b9 | 908.4101 | 908.4085 | -1.8 |
| b10 | 977.4321 | 977.4325 | 0.4 |
| b11 | 1092.4590 | 1092.4586 | -0.4 |
| y4 | 361.1546 | 361.1528 | -5.0 |
| y5 | 508.2230 | 508.2221 | -1.8 |
| y6 | 623.2500 | 623.2465 | -5.6 |
| y7 | 692.2720 | 692.2711 | -1.3 |
| $[M+2H]^{2+}$ | 800.3410 | 800.3382 | -3.5 |

**Figure S15.** Structural characterization via MS/MS of the SpaKC-1-modified SpaA1.3 product.

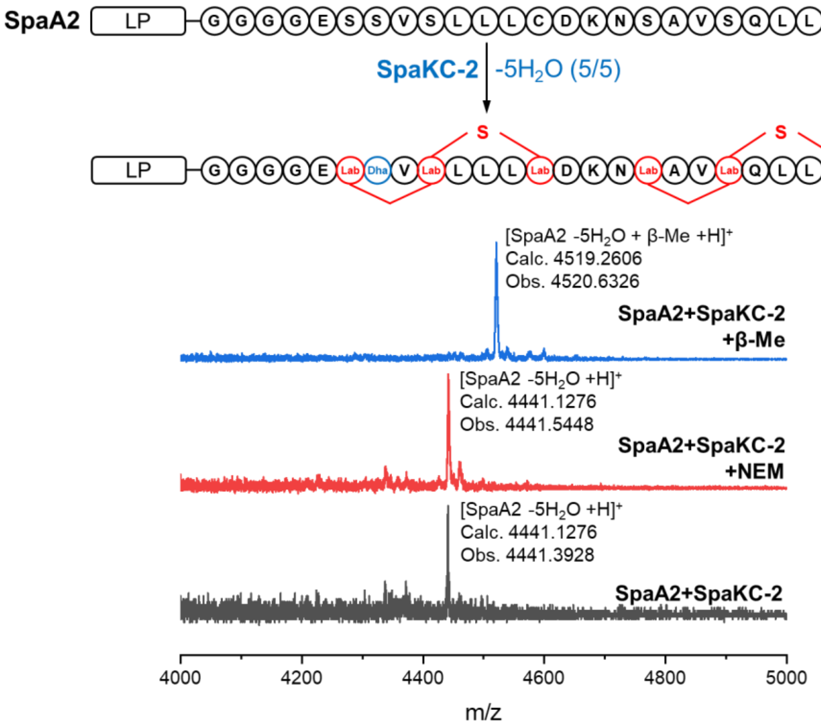

**Figure S16.** MALDI-TOF MS analysis and NEM/β-Me derivatization of product from *in vitro* modification of SpaKC-2 and precursor peptide SpaA2.

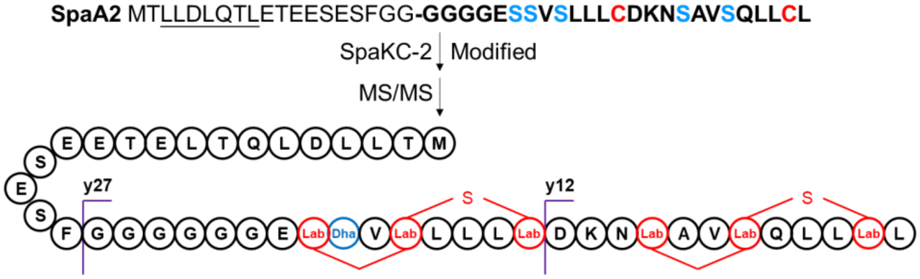

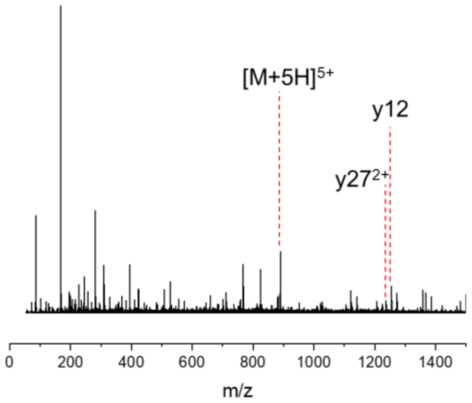

| Species | Calc. mass. (Da) | Obs. mass. (Da) | Err. (ppm) |
|---|---|---|---|
| y12 | 1254.6520 | 1254.6490 | -2.9 |
| $y27^{2+}$ | 1237.6130 | 1237.6140 | 0.6 |
| $[M+5H]^{5+}$ | 889.0314 | 889.0369 | 6.2 |

**Figure S17.** Structural characterization via MS/MS of the SpaKC-2-modified SpaA2 product.

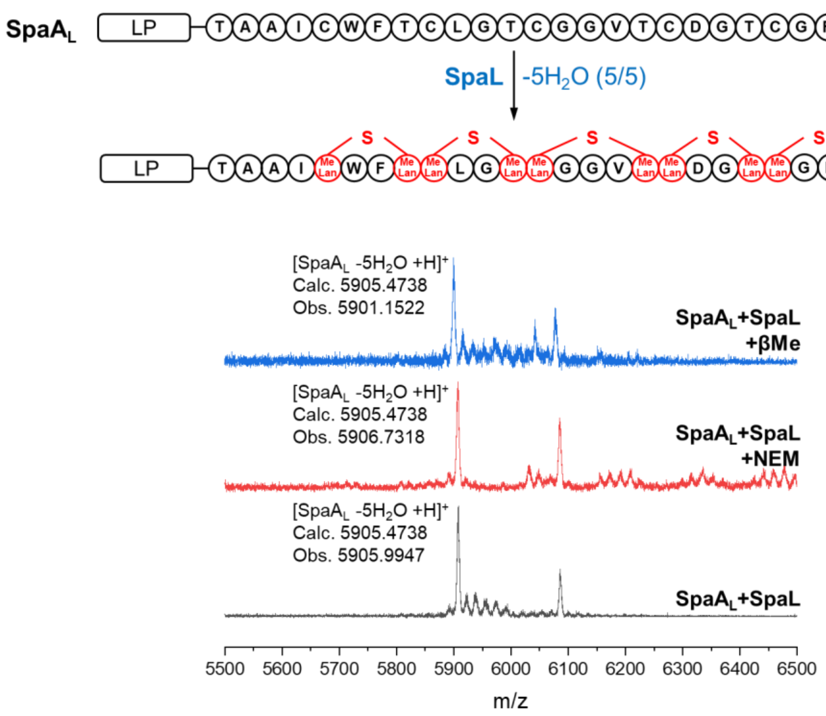

**Figure S18.** MALDI-TOF MS analysis and NEM/β-Me derivatization of products from *in vivo* coexpression of SpaL and precursor peptide SpaA$_L$.

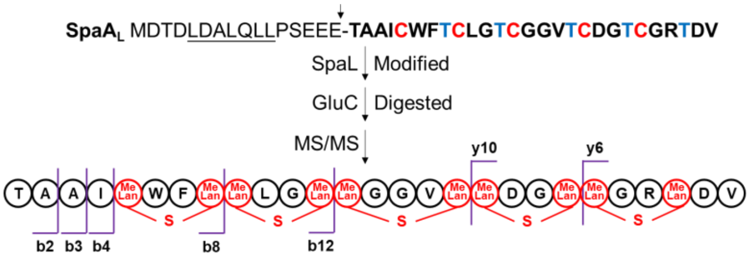

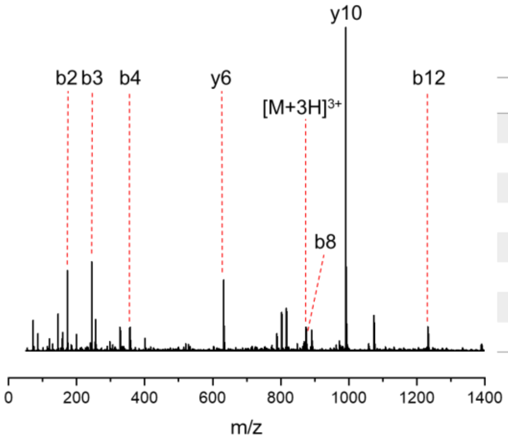

| Species | Calc. mass. (Da) | Obs. mass. (Da) | Err. (ppm) |
|---|---|---|---|
| b2 | 173.0921 | 173.0919 | -1.2 |
| b3 | 244.1292 | 244.1287 | -2.0 |
| b4 | 357.2133 | 357.2116 | -4.8 |
| b8 | 876.4079 | 876.4109 | 3.4 |
| b12 | 1232.5603 | 1232.5583 | -1.6 |
| y6 | 632.2827 | 632.2829 | 0.3 |
| y10 | 990.3780 | 990.3777 | -0.3 |
| [M+3H]$^{3+}$ | 874.3679 | 874.3638 | -4.7 |

**Figure S19.** Structural characterization via MS/MS of the SpaL-modified SpaA$_L$ product.

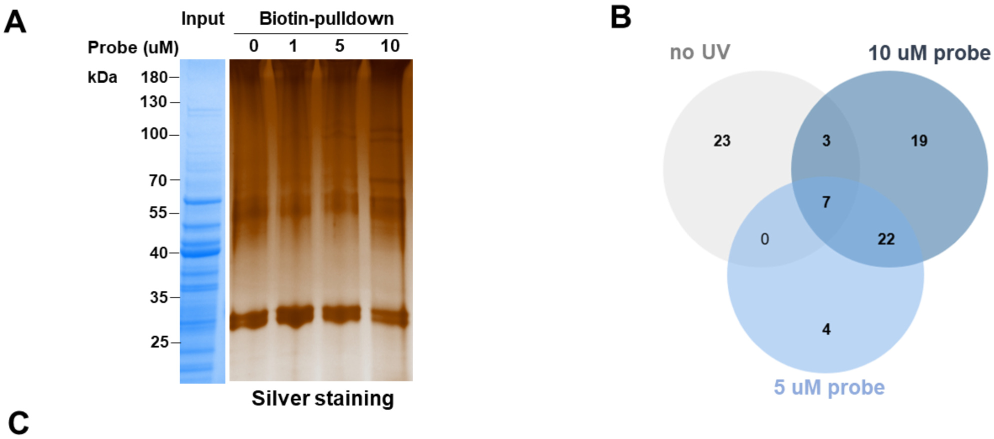

C

| No. | Protein | Protein Accession | MW [kDa] | Description |
|---|---|---|---|---|
| 1 | SpaP1 | A0A1R1SK32 | 94.6 | Aminopeptidase N |
| 2 | SpaDLD | A0A1R1SRQ8 | 49.6 | Dihydrolipoyl dehydrogenase |
| 3 | SpaKC-2 | A0A1R1SIB9 | 100.5 | Lanthipeptide synthetase |
| 4 | SpaP2 | A0A1R1SJF9 | 95.8 | Aminopeptidase N |
| 5 | SpaR | A0A1R1SFE5 | 36.3 | Putative transcriptional regulator-like protein |
| 6 | SpaL | A0A1R1SNM5 | 98.7 | Lanthipeptide synthetase |
| 7 | SpaKC-1 | A0A1R1SP09 | 92.3 | Lanthipeptide synthetase |
| 8 | SpaXDH | A0A1R1S8W6 | 39.8 | Xanthine dehydrogenase |
| 9 | SpaRimK | A0A1R1S5M5 | 31.5 | RimK domain-containing protein |
| 10 | SpaSBP | A0A1R1SBD3 | 50.6 | Family 5 extracellular solute-binding protein |
| 11 | SpaDLD2 | A0A1R1SHN3 | 48.6 | Dihydrolipoyl dehydrogenase |
| 12 | SpaP3-M6 | A0A1R1S661 | 87.1 | Putative secreted protease |
| 13 | SpaMbtB | A0A1R1SC91 | 200.9 | Phenyloxazoline synthase MbtB |
| 14 | SpaADH | A0A1R1SFT7 | 36.3 | Acetoin dehydrogenase |
| 15 | SpaFeS | A0A1R1SHN6 | 12.4 | FeS cluster biogenesis domain-containing protein |
| 16 | SpaPNPT | A0A1R1SQT0 | 80.6 | Polyribonucleotide nucleotidyltransferase |
| 17 | SpaAH | A0A1R1S614 | 52.4 | Adenosylhomocysteinase |
| 18 | SpaBCCP | A0A1R1SF10 | 61.2 | Biotin carboxylase |
| 19 | SpaSI | A0A1R1SF21 | 32.5 | Siderophore-interacting protein |
| 20 | SpaRHA | A0A1R1SI25 | 59.9 | FAD-binding PCMH-type domain-containing protein |
| 21 | SpaDnaK | A0A1R1SLI9 | 66.4 | Chaperone protein DnaK |
| 22 | SpaMMP | A0A1R1SPZ1 | 59.0 | Neutral metalloproteinase |

**Figure S20.** (A) Silver staining of proteins captured by the SpaA1.2LP-P2 probe from *S. sparsogenes* cell lysates. Coomassie Brilliant Blue staining of the total lysate served as an input control. Silver staining revealed two prominent bands at approximately 100 kDa, corresponding to the presence of the target proteins SpaP1, SpaP2, SpaKC-1, SpaKC-2, and SpaL. (B) Venn diagram showing proteins significantly enriched by the photoaffinity probe SpaA1.2LP-P2 in *S. sparsogenes* cell lysates. (C) Chemical proteomics identification of 22 proteins from *S. sparsogenes*.

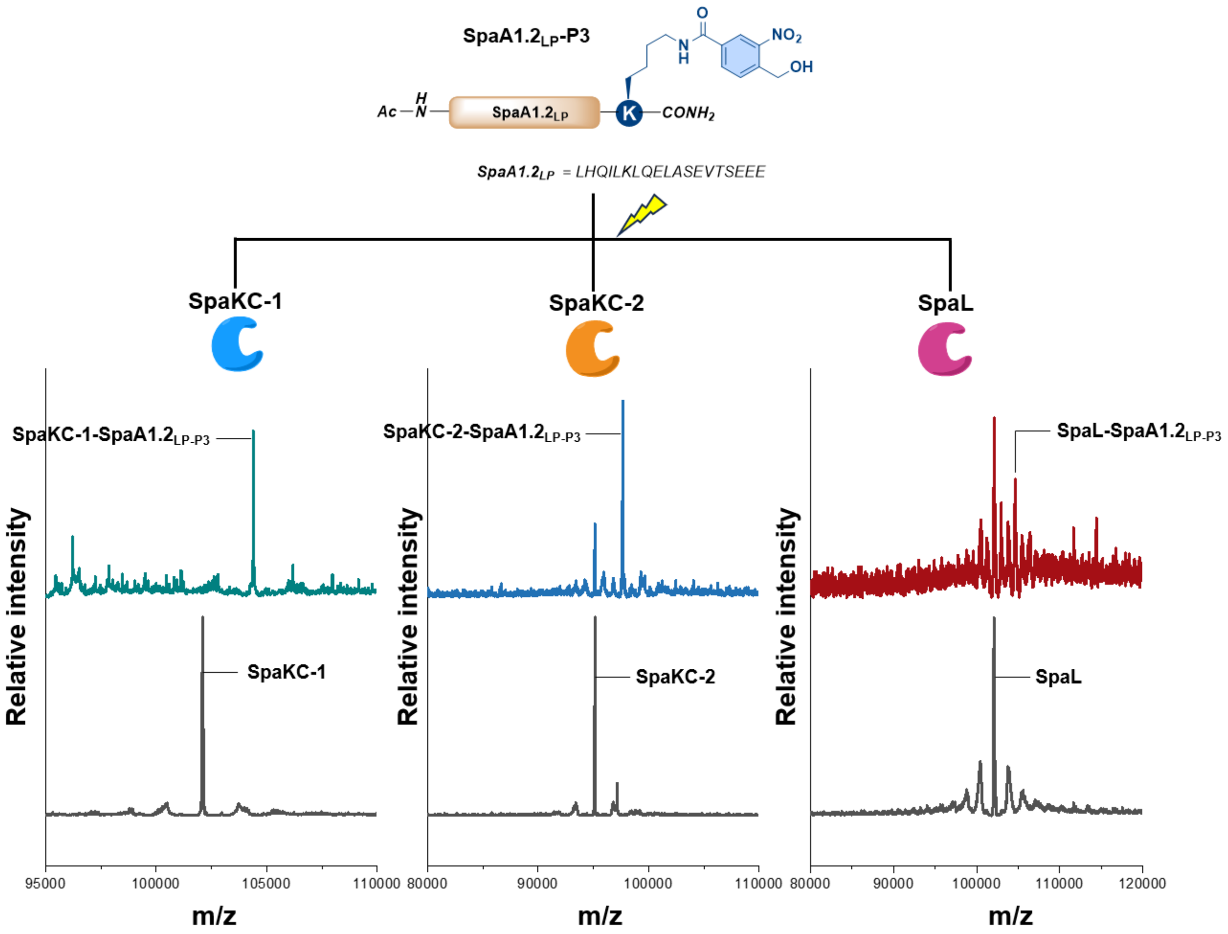

**Figure S21.** HRMS analysis of photo-crosslinking between SpaA1.2LP-P3 and SpaKC-1, SpaKC-2 and SpaL. SpaKC-1: calcd: 95131 Da, obs.: 95132 Da. SpaKC-1-SpaA1.2LP-P3: calcd: 97638 Da, obs.: 97643 Da. SpaKC-2: calcd: 102070 Da, obs.: 102078 Da. SpaKC-2-SpaA1.2LP-P3: calcd: 104577 Da, obs.: 104580 Da. SpaL: calcd: 102115 Da, obs.: 102113 Da. SpaL-SpaA1.2LP-P3: calcd: 104622 Da, obs.: 104624 Da.

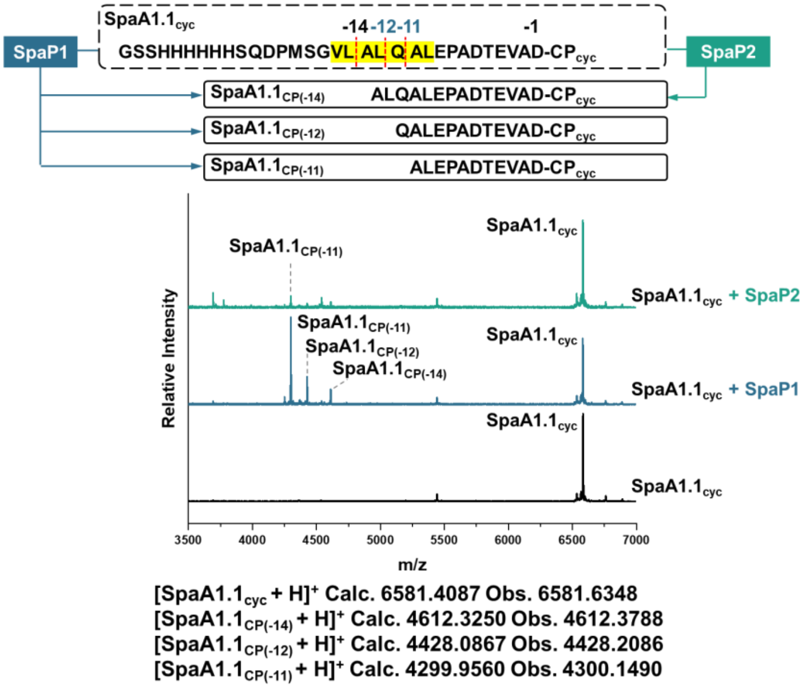

**Figure S22.** MALDI-TOF MS analysis of the endopeptidase activity of SpaP1 and SpaP2 toward SpaA1.1cyc.

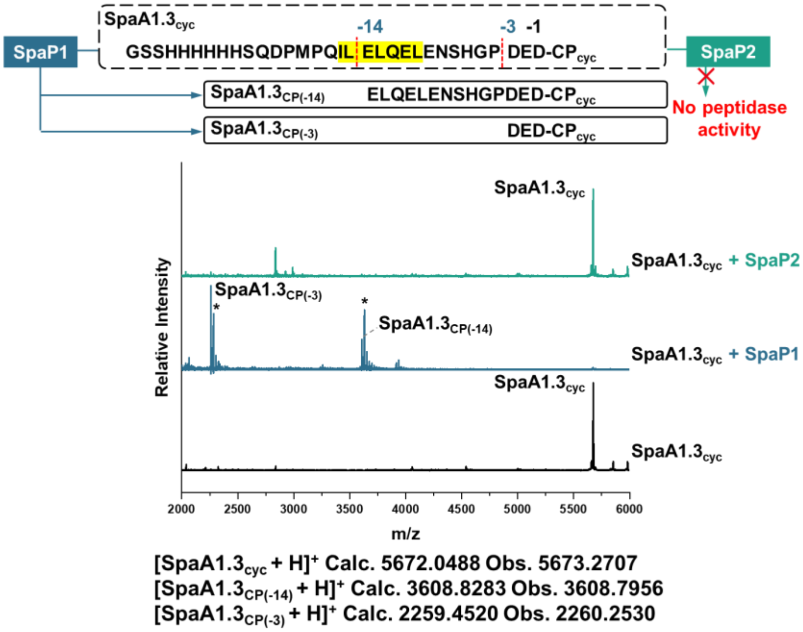

**Figure S23.** MALDI-TOF MS analysis of the endopeptidase activity of SpaP1 and SpaP2 toward SpaA1.3cyc.

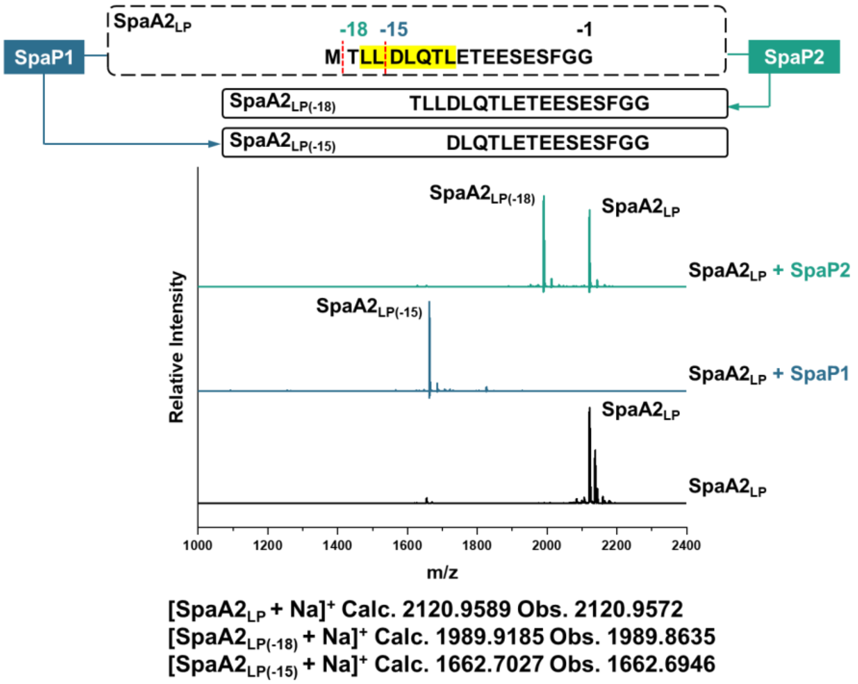

**Figure S24.** MALDI–TOF MS analysis of the endopeptidase activity of SpaP1 and SpaP2 toward $SpaA2_{LP}$.

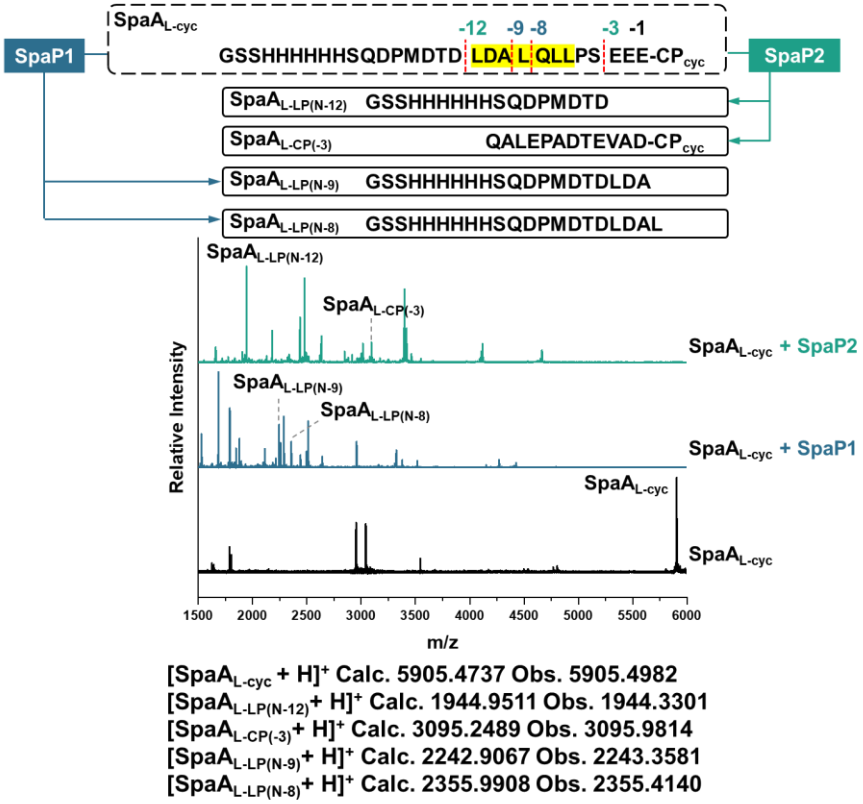

**Figure S25.** MALDI–TOF MS analysis of the endopeptidase activity of SpaP1 and SpaP2 toward $SpaA_{L\text{-}cyc}$.

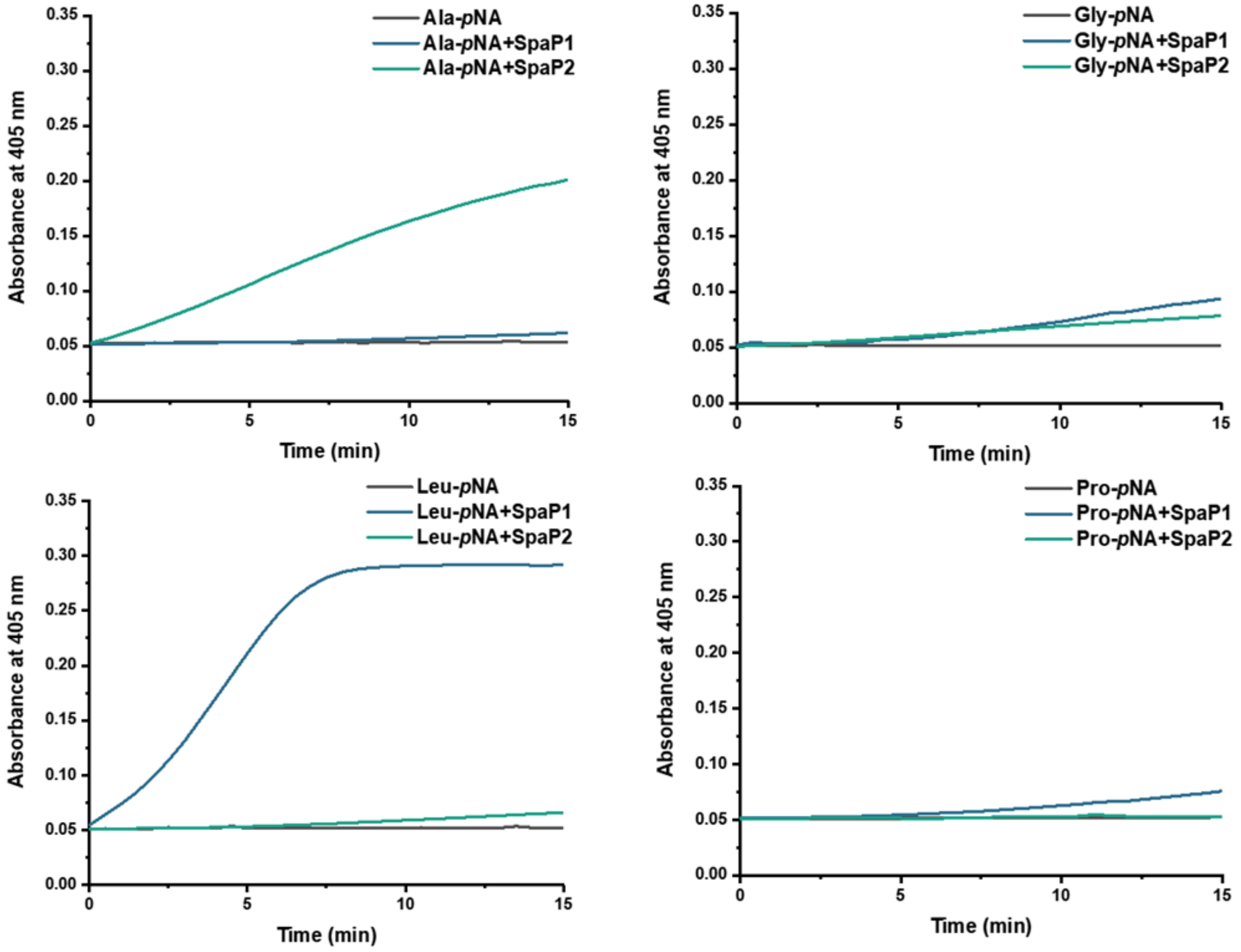

**Figure S26.** The aminopeptidase activity of SpaP1 and SpaP2. Both enzymes exhibited activity against amino acid *p*NA derivatives, albeit with different substrate preferences.

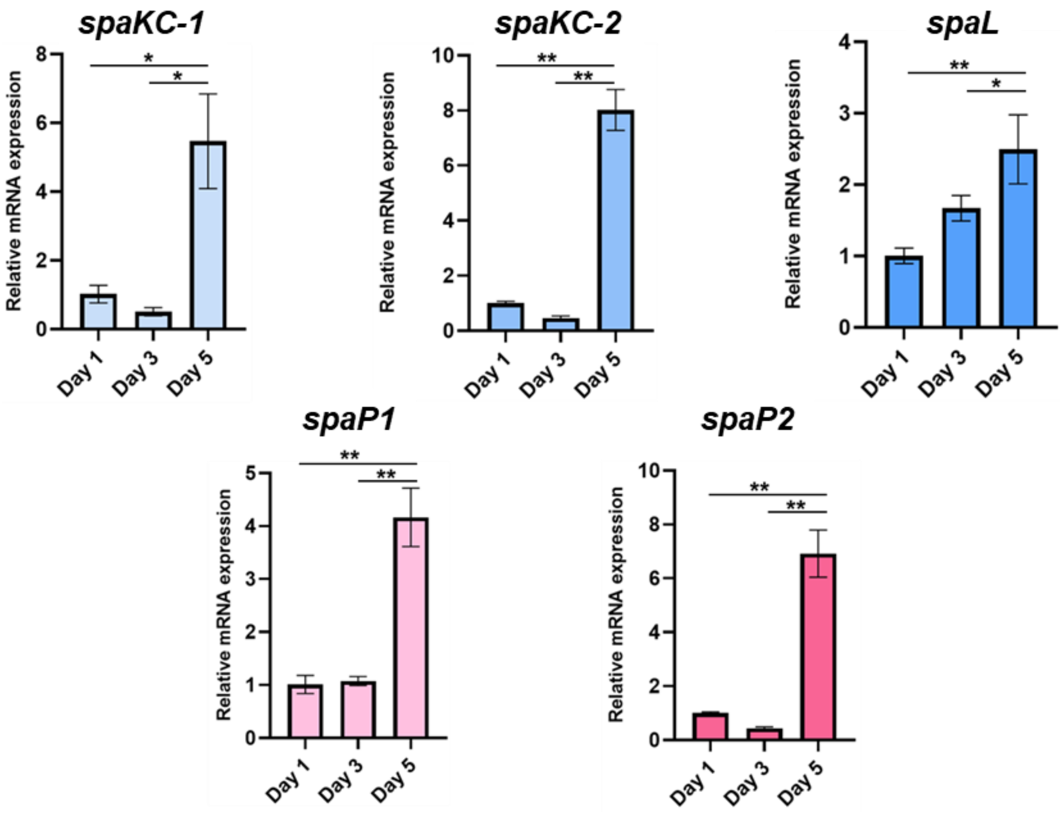

**Figure S27.** qRT-PCR analysis of mRNA expression levels of the five probe-identified target proteins in *Streptomyces sparsogenes* ATCC 25498. mRNA expression levels of the five probe-identified target proteins reached their highest point on the fifth day. Data are shown as means SD (n=3). Statistical analysis was performed using one-way ANOVA, specific P-values are shown in the source data. (ns: not significant, $p > 0.05$, $^*p < 0.05$, $^{**}p < 0.01$, and $^{***}p < 0.001$)

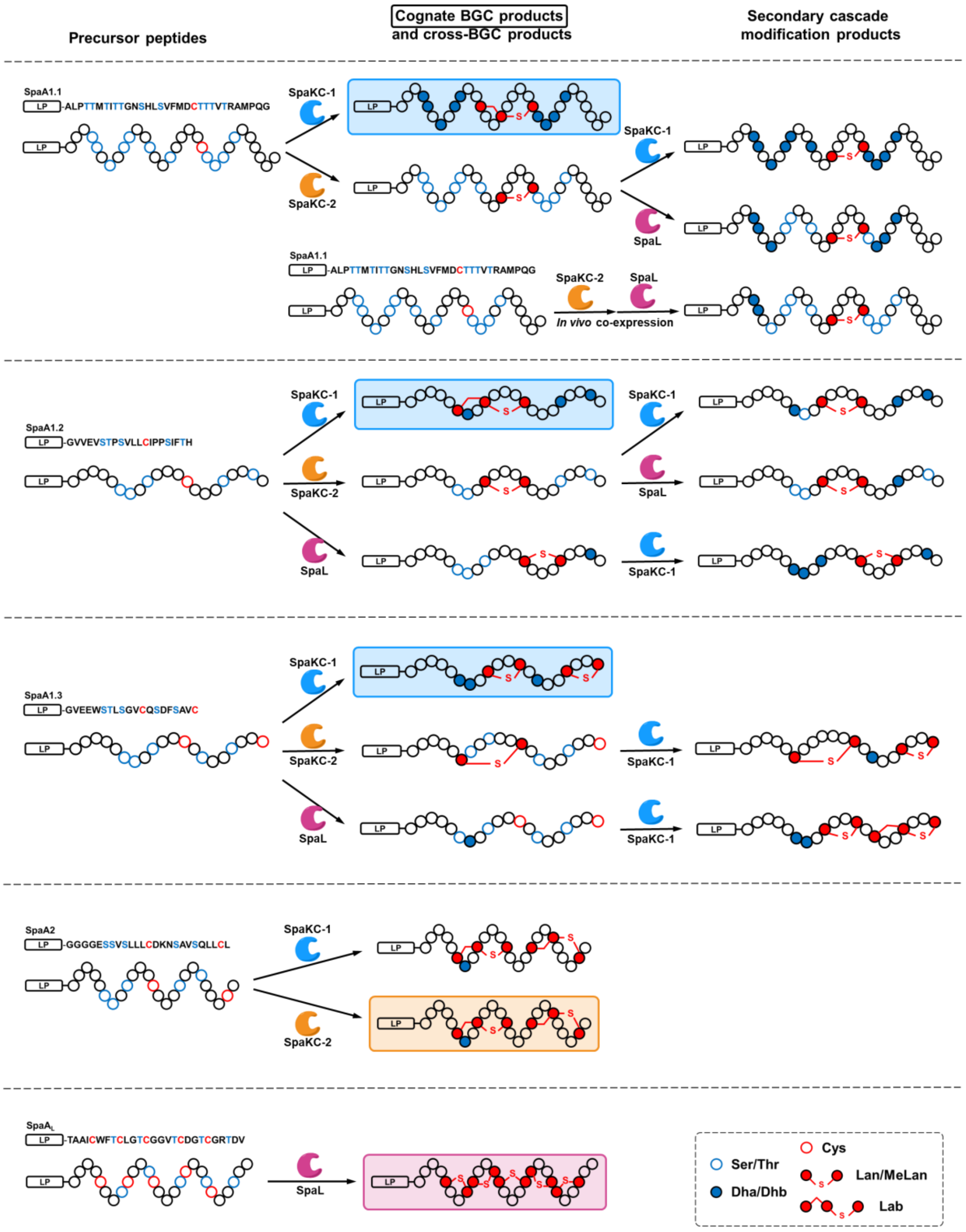

**Figure S28.** Structural comparison of combinatorial biosynthesis products from *lan III-1*, *lan III-2* and *lan IV* BGCs. Cognate BGC products are highlighted in the boxes.

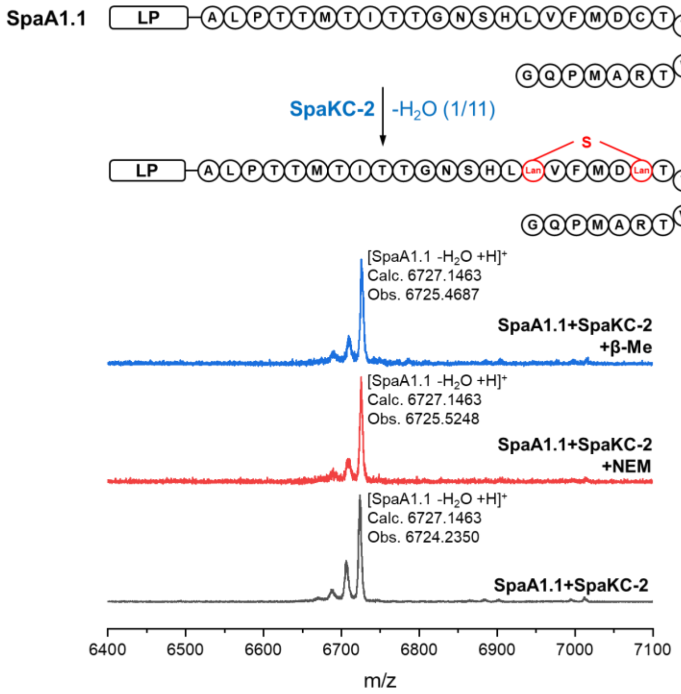

**Figure S29.** MALDI-TOF MS analysis and NEM/β-Me derivatization of product by *in vivo* coexpression of SpaKC-2 from the *lan III-2* cluster with SpaA1.1 from the *lan III-1* cluster.

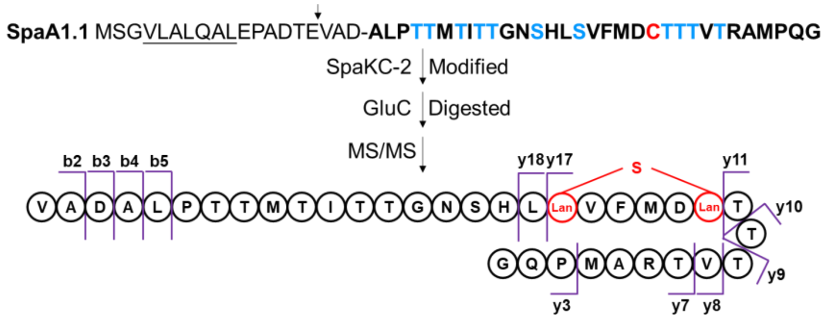

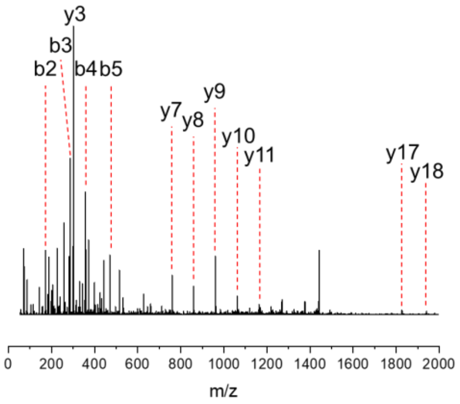

| Species | Calc. mass. (Da) | Obs. mass. (Da) | Err. (ppm) |
|---|---|---|---|
| b2 | 171.1129 | 171.1133 | 2.3 |
| b3 | 286.1398 | 286.1391 | -2.4 |
| b4 | 357.1769 | 357.1763 | -1.7 |
| b5 | 470.2610 | 470.2598 | -2.6 |
| y3 | 301.1507 | 301.1499 | -2.7 |
| y7 | 760.3771 | 760.3763 | -1.1 |
| y8 | 859.4455 | 859.4451 | -0.5 |
| y9 | 960.4932 | 960.4922 | -1.0 |
| y10 | 1061.5408 | 1061.5347 | -5.7 |
| y11 | 1162.5885 | 1162.5853 | -2.8 |
| y17 | 1826.8240 | 1826.8228 | -0.7 |
| y18 | 1939.9081 | 1939.9007 | -3.8 |

**Figure S30.** Structural characterization via MS/MS of the SpaKC-2-modified SpaA1.1 product.

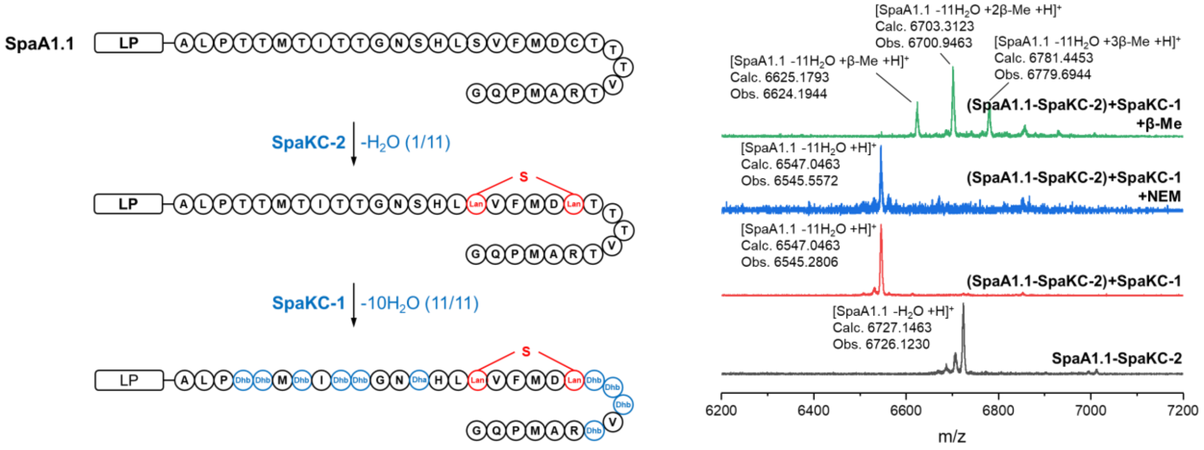

**Figure S31.** MALDI-TOF MS analysis and NEM/β-Me derivatization of the *in vitro* cascade modification product SpaA1.1–SpaKC-2–SpaKC-1.

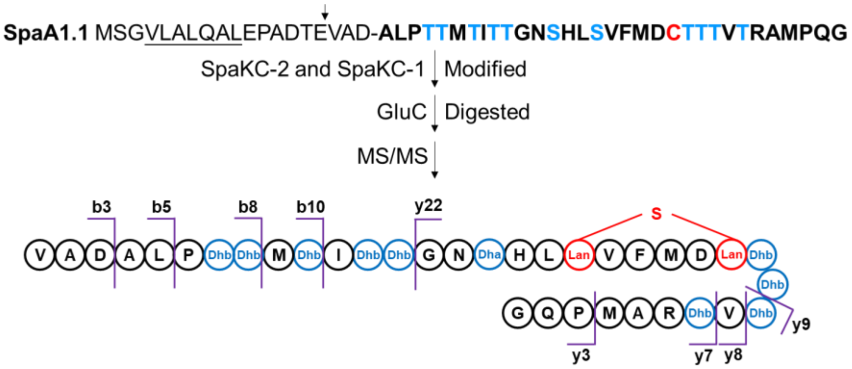

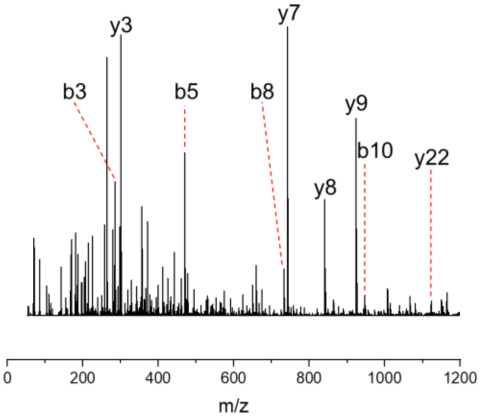

| Species | Calc. mass. (Da) | Obs. mass. (Da) | Err. (ppm) |
|---|---|---|---|
| b3 | 286.1398 | 286.1394 | -1.4 |
| b5 | 470.2610 | 470.2608 | -0.4 |
| b8 | 733.3891 | 733.3902 | 1.5 |
| b10 | 947.4672 | 947.4703 | 3.3 |
| y3 | 301.1507 | 301.1505 | -0.7 |
| y7 | 742.3671 | 742.3681 | 1.3 |
| y8 | 841.4355 | 841.4370 | 1.8 |
| y9 | 924.4732 | 924.4755 | 2.5 |
| $y22^{2+}$ | 1123.0104 | 1123.0101 | -0.3 |

**Figure S32.** Structural characterization via MS/MS of the cascade modification product SpaA1.1–SpaKC-2–SpaKC-1.

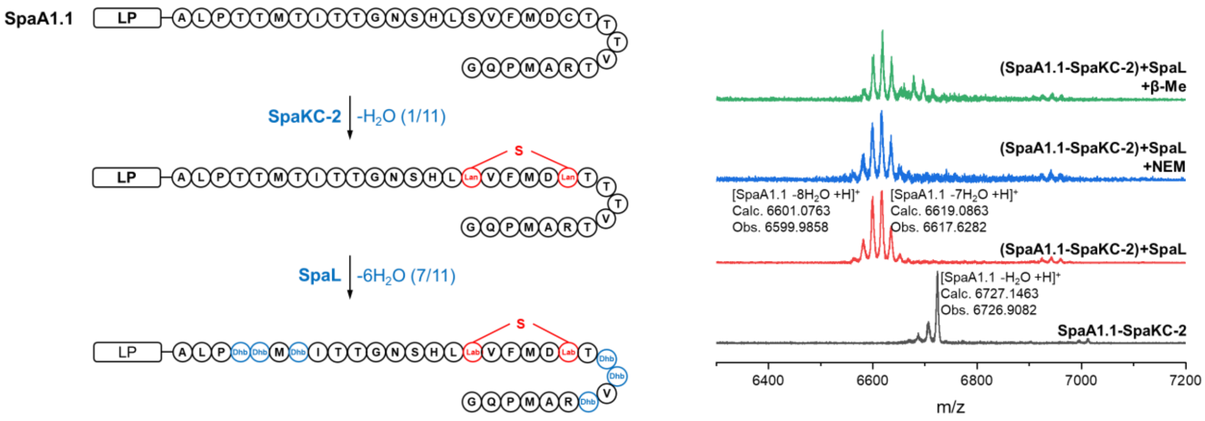

**Figure S33.** MALDI-TOF MS analysis and NEM/β-Me derivatization of the *in vitro* cascade modification product SpaA1.1–SpaKC-2–SpaL.

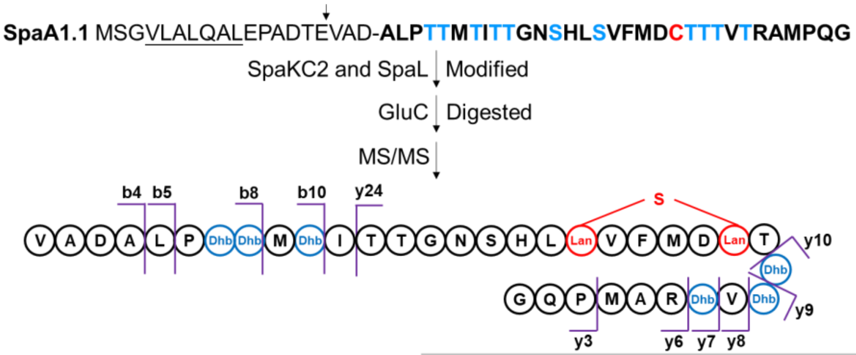

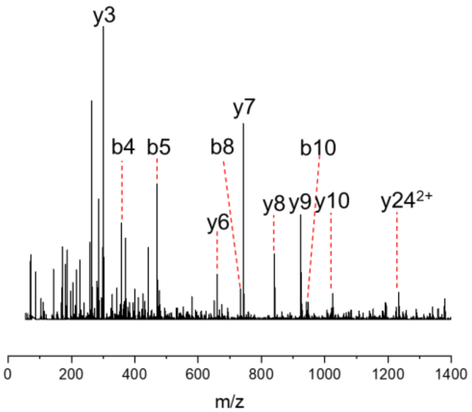

| Species | Calc. mass. (Da) | Obs. mass. (Da) | Err. (ppm) |
|---|---|---|---|
| b4 | 357.1769 | 357.1772 | 0.8 |
| b5 | 470.2610 | 470.2614 | 0.9 |
| b8 | 733.3891 | 733.3878 | -1.8 |
| b10 | 947.4672 | 947.4659 | -1.4 |
| y3 | 301.1507 | 301.1515 | 2.7 |
| y6 | 659.3294 | 659.3308 | 2.1 |
| y7 | 742.3671 | 742.3678 | 0.9 |
| y8 | 841.4355 | 841.4362 | 0.8 |
| y9 | 924.4732 | 924.4726 | -0.6 |
| y10 | 1025.5208 | 1025.5203 | -0.5 |
| $y24^{2+}$ | 1233.0630 | 1233.0656 | 2.1 |

**Figure S34.** Structural characterization via MS/MS of the cascade modification product SpaA1.1–SpaKC-2–SpaL.

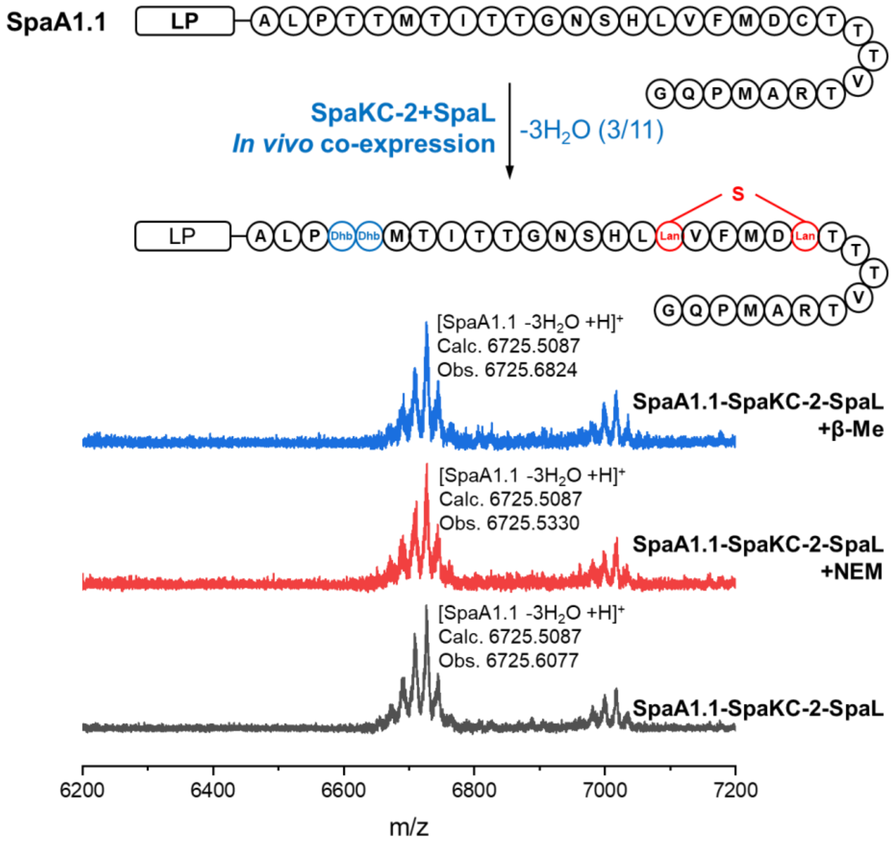

**Figure S35.** MALDI-TOF MS analysis and NEM/β-Me derivatization of product by *in vivo* coexpression of the cascade modification product SpaA1.1–SpaKC-2–SpaL.

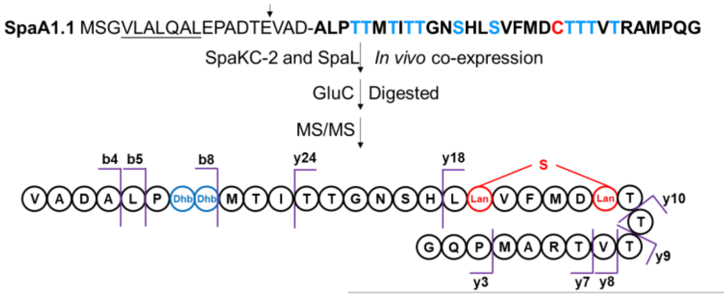

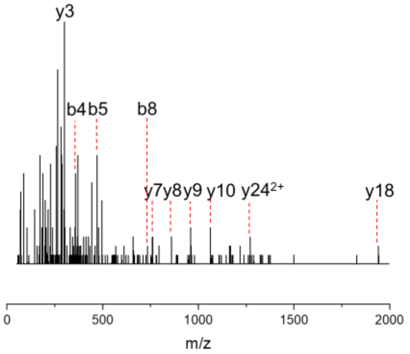

| Species | Calc. mass. (Da) | Obs. mass. (Da) | Err. (ppm) |
|---|---|---|---|
| b4 | 357.1769 | 357.1771 | 0.6 |
| b5 | 470.2610 | 470.2613 | 0.6 |
| b8 | 733.3891 | 733.3885 | -0.8 |
| y3 | 301.1507 | 301.1497 | -3.3 |
| y7 | 760.3771 | 760.3756 | -2.0 |
| y8 | 859.4455 | 859.4498 | 5.0 |
| y9 | 960.4932 | 960.4906 | -2.7 |
| y10 | 1061.5408 | 1061.5418 | 0.9 |
| y18 | 1939.9081 | 1939.9007 | -3.8 |
| y24$^{2+}$ | 1269.0830 | 1269.0786 | -3.5 |

**Figure S36.** Structural characterization via MS/MS of the *in vivo* coexpression cascade modification product SpaA1.1–SpaKC-2–SpaL.

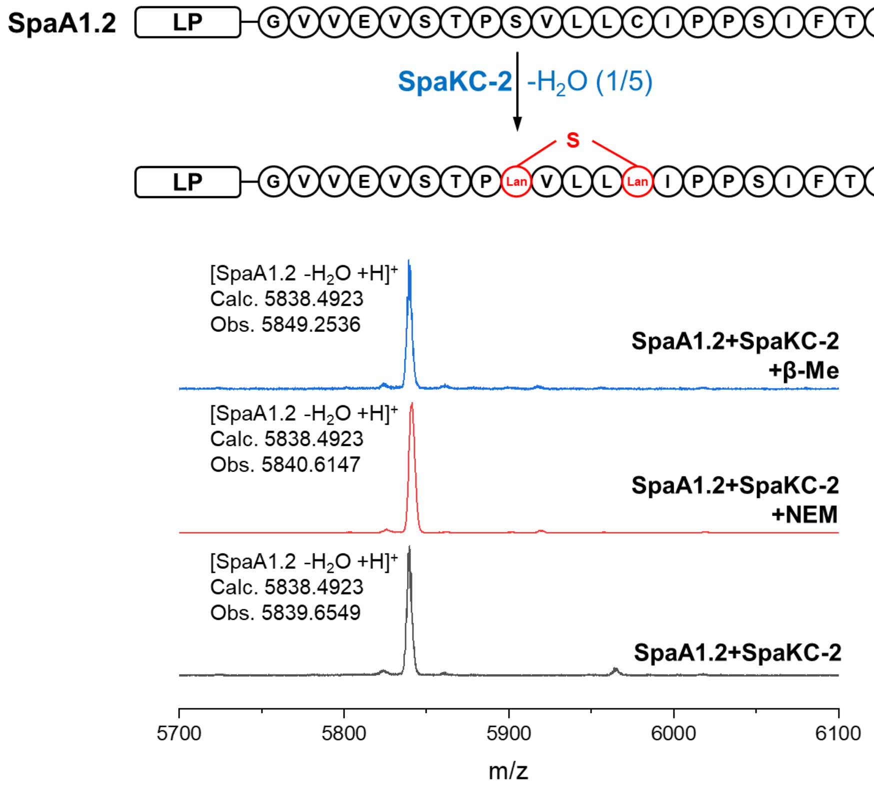

**Figure S37.** MALDI-TOF MS analysis and NEM/β-Me derivatization of product by *in vitro* modification of SpaKC-2 from *lan III-2* and precursor peptide SpaA1.2 from *lan III-1*.

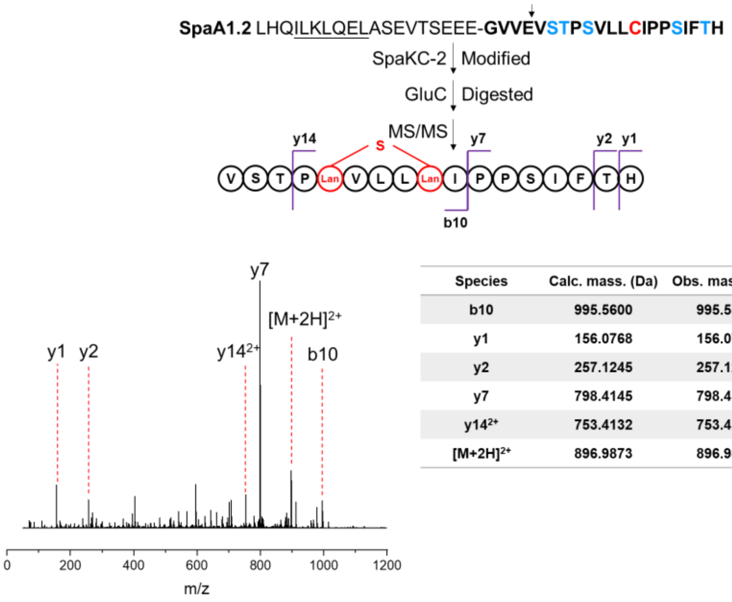

| Species | Calc. mass. (Da) | Obs. mass. (Da) | Err. (ppm) |
|---|---|---|---|
| b10 | 995.5600 | 995.5560 | -4.0 |
| y1 | 156.0768 | 156.0761 | -4.5 |
| y2 | 257.1245 | 257.1233 | -4.7 |
| y7 | 798.4145 | 798.4107 | -4.8 |
| $y14^{2+}$ | 753.4132 | 753.4120 | -1.6 |
| $[M+2H]^{2+}$ | 896.9873 | 896.9851 | -2.5 |

**Figure S38.** Structural characterization via MS/MS of the SpaKC-2-modified SpaA1.2 product.

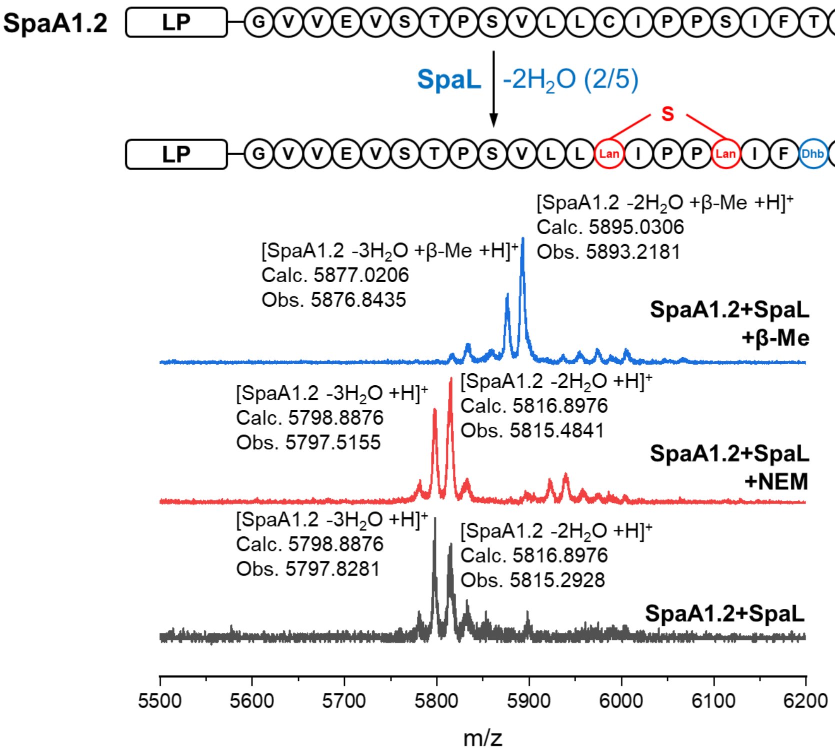

**Figure S39.** MALDI-TOF MS analysis and NEM/β-Me derivatization of product by *in vitro* modification of SpaL from *lan IV* and precursor peptide SpaA1.2 from *lan III-1*.

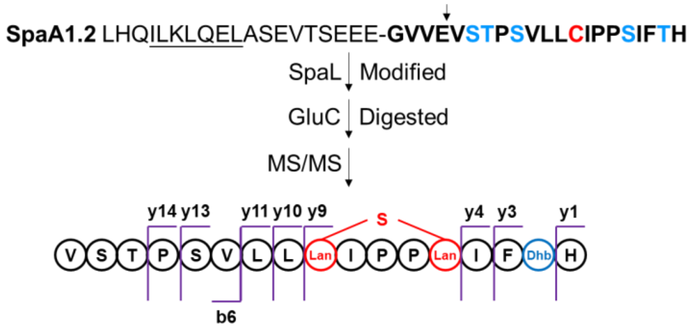

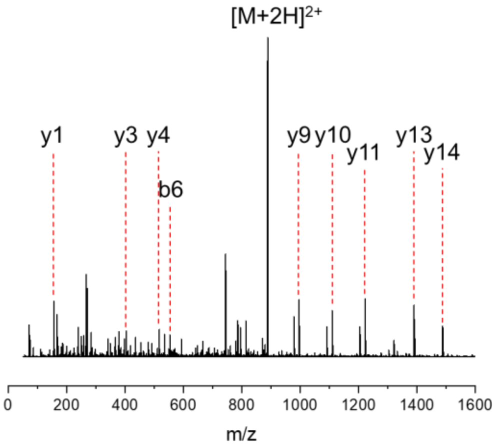

| Species | Calc. mass. (Da) | Obs. mass. (Da) | Err. (ppm) |
|---|---|---|---|
| b6 | 553.2986 | 553.2985 | -0.2 |
| y1 | 156.0768 | 156.0765 | -1.9 |
| y3 | 404.1929 | 404.1931 | 0.5 |
| y4 | 517.2770 | 517.2769 | -0.2 |
| y9 | 996.4978 | 996.5000 | 2.2 |
| y10 | 1109.5818 | 1109.5842 | 2.2 |
| y11 | 1222.6659 | 1222.6688 | 2.4 |
| y13 | 1390.7563 | 1390.7596 | 2.4 |
| y14 | 1487.8091 | 1487.8140 | 3.3 |
| $[M+2H]^{2+}$ | 887.9823 | 887.9856 | 3.7 |

**Figure S40.** Structural characterization via MS/MS of the SpaL-modified SpaA1.2 product.

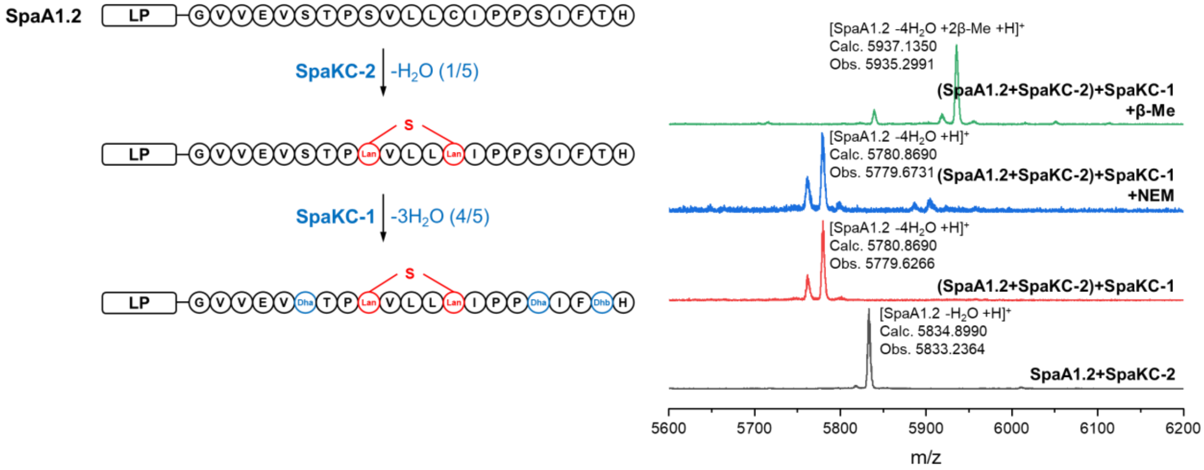

**Figure S41.** MALDI-TOF MS analysis and NEM/β-Me derivatization of the *in vitro* cascade modification product SpaA1.2–SpaKC-2–SpaKC-1.

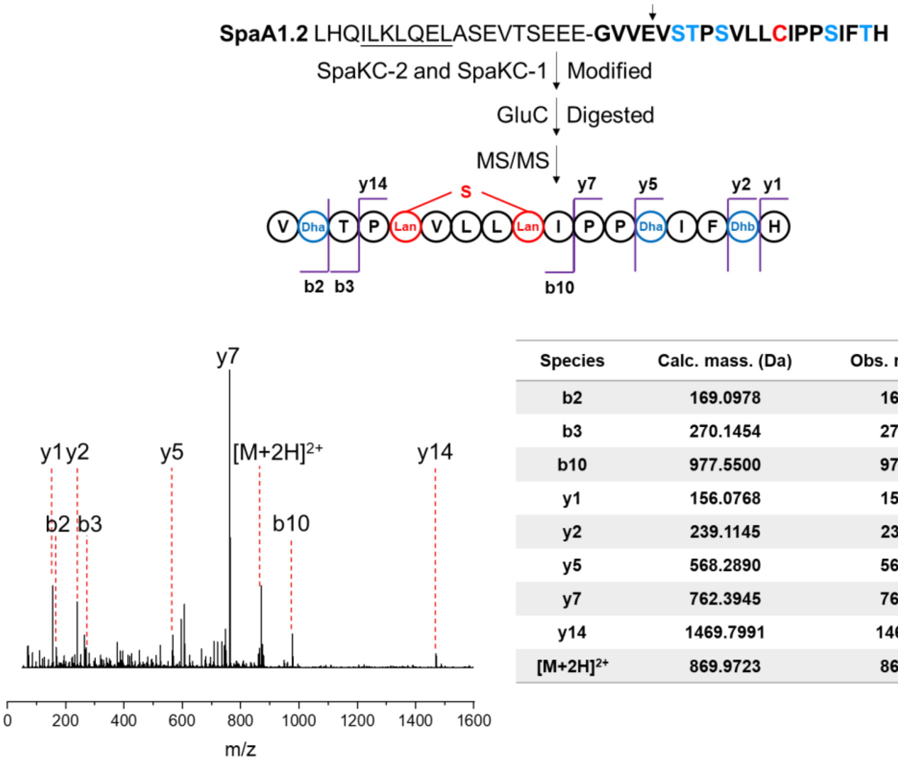

| Species | Calc. mass. (Da) | Obs. mass. (Da) | Err. (ppm) |
|---|---|---|---|
| b2 | 169.0978 | 169.0974 | -2.4 |
| b3 | 270.1454 | 270.1455 | 0.4 |
| b10 | 977.5500 | 977.5527 | 2.8 |
| y1 | 156.0768 | 156.0770 | 1.3 |
| y2 | 239.1145 | 239.1146 | 0.4 |
| y5 | 568.2890 | 568.2890 | 0.0 |
| y7 | 762.3945 | 762.3955 | 1.3 |
| y14 | 1469.7991 | 1469.8026 | 2.4 |
| $[M+2H]^{2+}$ | 869.9723 | 869.9747 | 2.8 |

**Figure S42.** Structural characterization via MS/MS of the cascade modification product SpaA1.2–SpaKC-2–SpaKC-1.

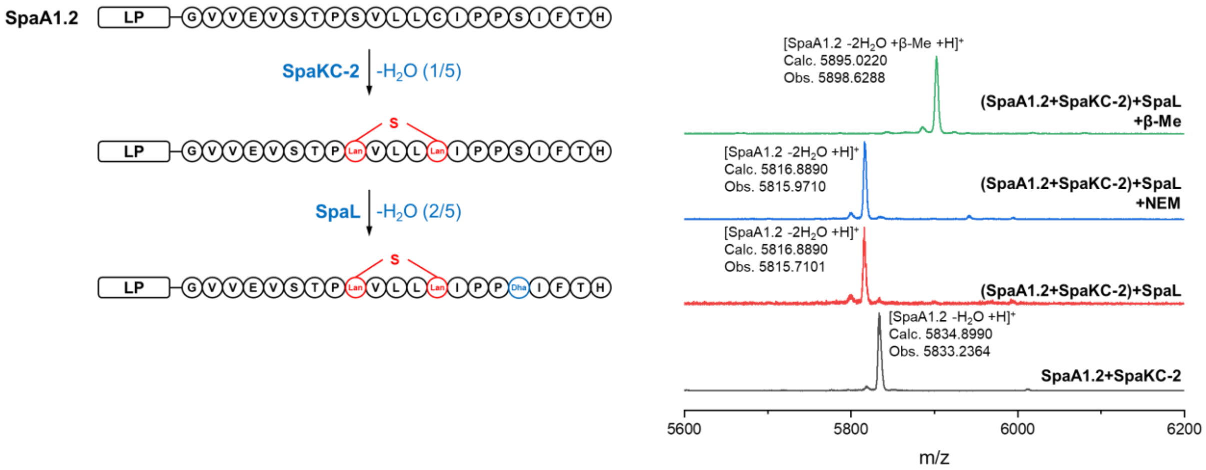

**Figure S43.** MALDI-TOF MS analysis and NEM/β-Me derivatization of the *in vitro* cascade modification product SpaA1.2–SpaKC-2–SpaL.

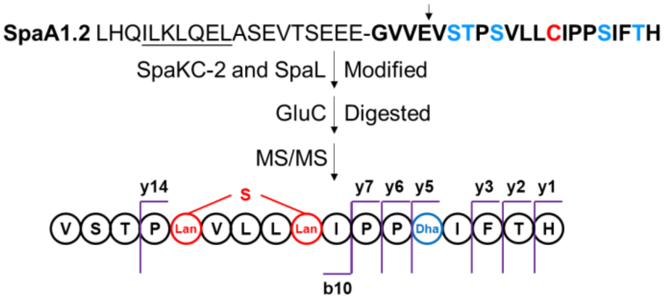

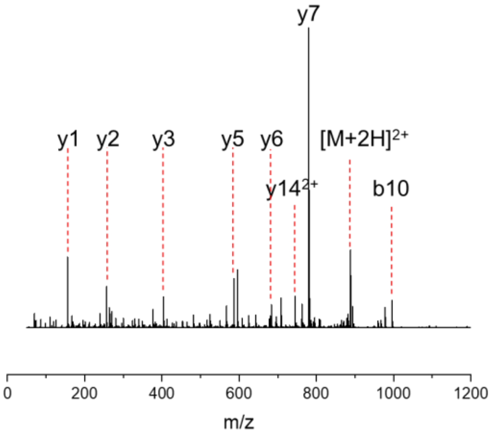

| Species | Calc. mass. (Da) | Obs. mass. (Da) | Err. (ppm) |
|---|---|---|---|
| b10 | 995.5600 | 995.5608 | 0.8 |
| y1 | 156.0768 | 156.0767 | -0.6 |
| y2 | 257.1245 | 257.1248 | 1.2 |
| y3 | 404.1929 | 404.1926 | -0.7 |
| y5 | 586.2989 | 586.2981 | -1.4 |
| y6 | 683.3517 | 683.3512 | -0.7 |
| y7 | 780.4045 | 780.4033 | -1.5 |
| $y14^{2+}$ | 744.4082 | 744.4076 | -0.8 |
| $[M+2H]^{2+}$ | 887.9823 | 887.9821 | -0.2 |

**Figure S44.** Structural characterization via MS/MS of the cascade modification product SpaA1.2–SpaKC-2–SpaL.

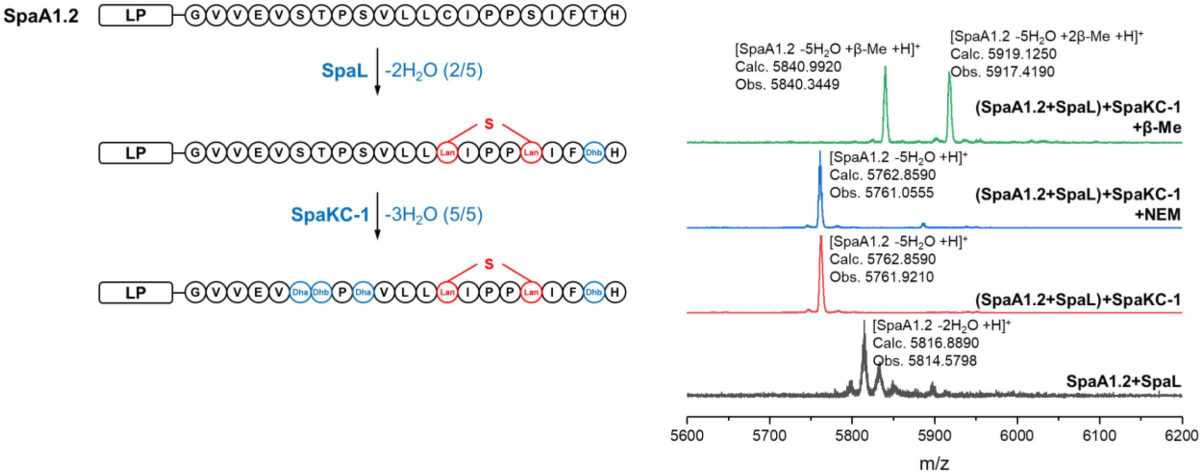

**Figure S45.** MALDI-TOF MS analysis and NEM/β-Me derivatization of the *in vitro* cascade modification product SpaA1.2–SpaL–SpaKC-1.

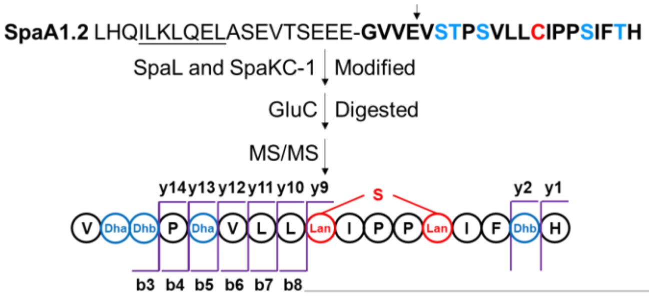

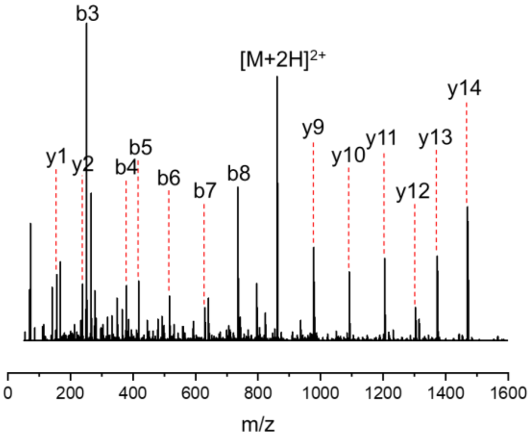

| Species | Calc. mass. (Da) | Obs. mass. (Da) | Err. (ppm) |
|---|---|---|---|
| b3 | 252.1354 | 252.1344 | -4.0 |
| b4 | 349.1882 | 349.1869 | -3.7 |
| b5 | 418.2102 | 418.2082 | -4.8 |
| b6 | 517.2786 | 517.2766 | -3.9 |
| b7 | 630.3627 | 630.3609 | -2.9 |
| b8 | 743.4468 | 743.4449 | -2.6 |
| y1 | 156.0768 | 156.0766 | -1.3 |
| y2 | 239.1145 | 239.1141 | -1.7 |
| y9 | 978.4878 | 978.4875 | -0.3 |
| y10 | 1091.5718 | 1091.5725 | 0.6 |
| y11 | 1204.6559 | 1204.6569 | 0.8 |
| y12 | 1303.7243 | 1303.7247 | 0.3 |
| y13 | 1372.7463 | 1372.7462 | -0.1 |
| y14 | 1469.7991 | 1469.7994 | 0.2 |
| [M+2H]$^{2+}$ | 860.9673 | 860.9666 | -0.8 |

**Figure S46.** Structural characterization via MS/MS of the cascade modification product SpaA1.2–SpaL–SpaKC-1.

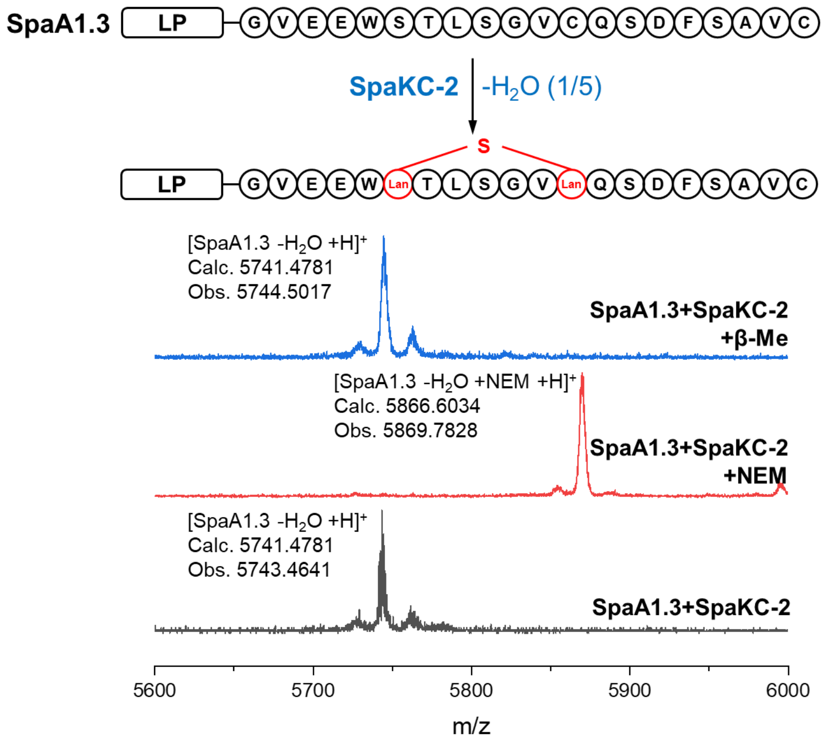

**Figure S47.** MALDI-TOF MS analysis and NEM/β-Me derivatization of product by *in vitro* modification of SpaKC-2 from *lan III-2* and precursor peptide SpaA1.3 from *lan III-1*.

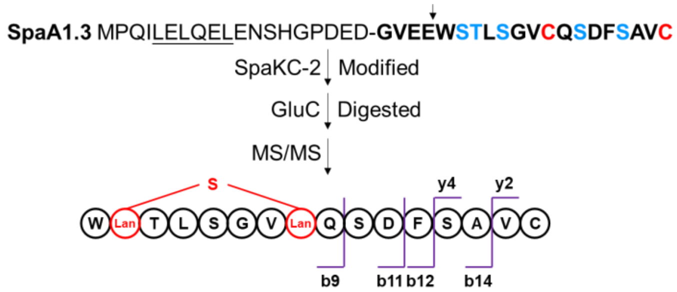

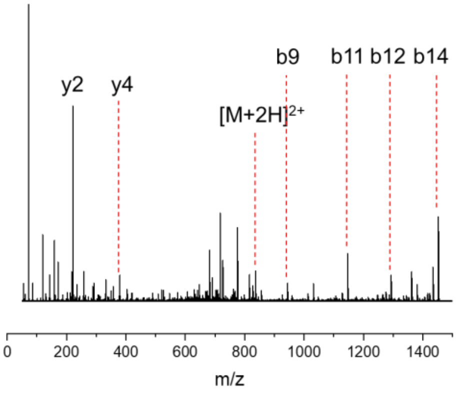

| Species | Calc. mass. (Da) | Obs. mass. (Da) | Err. (ppm) |
|---|---|---|---|
| b9 | 944.4301 | 944.4312 | 1.2 |
| b11 | 1146.4890 | 1146.4921 | 2.7 |
| b12 | 1293.5575 | 1293.5599 | 1.9 |
| b14 | 1451.6266 | 1451.6272 | 0.4 |
| y2 | 221.0955 | 221.0958 | 1.4 |
| y4 | 379.1646 | 379.1649 | 0.8 |
| $[M+2H]^{2+}$ | 836.3610 | 836.3659 | 5.9 |

**Figure S48.** Structural characterization via MS/MS of the SpaKC-2-modified SpaA1.3 product.

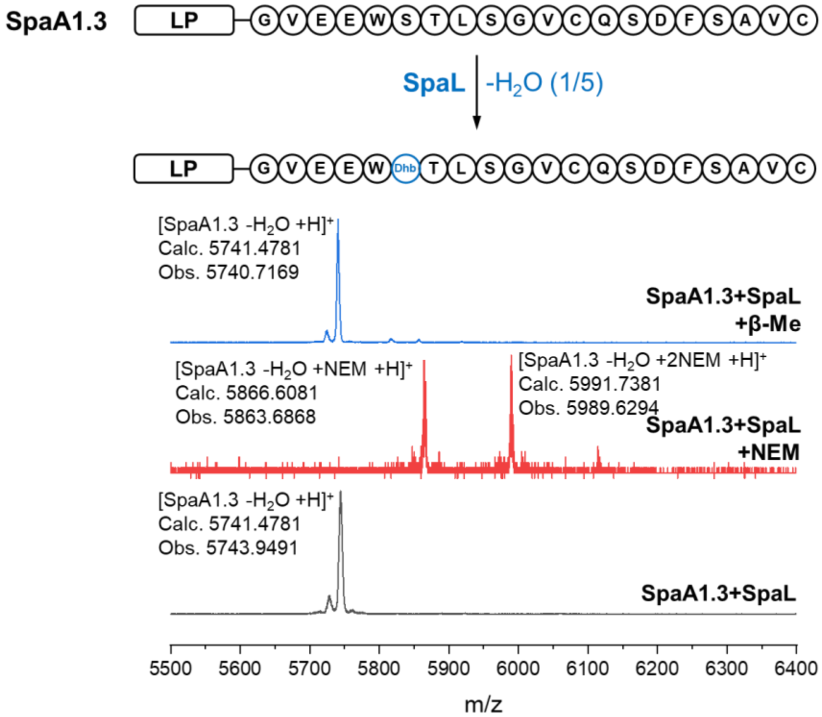

**Figure S49.** MALDI-TOF MS analysis and NEM/β-Me derivatization of product by *in vitro* modification of SpaL from *lan IV* and precursor peptide SpaA1.3 from *lan III-1*.

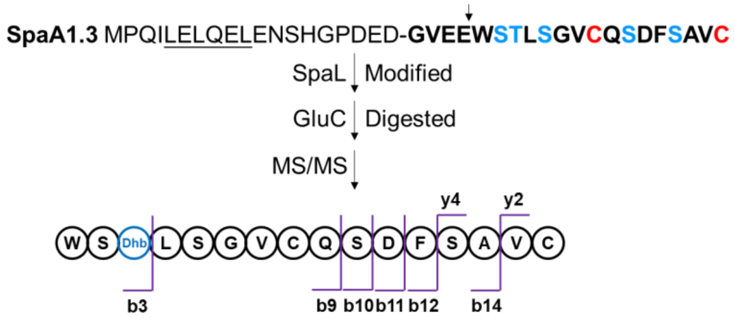

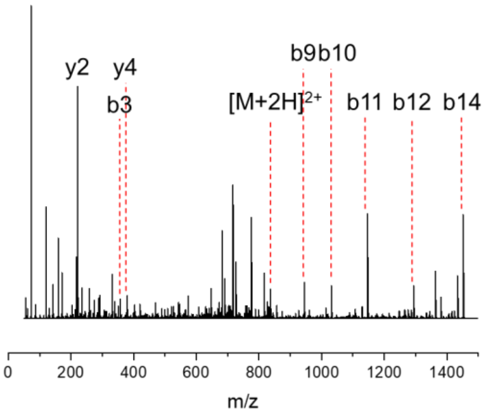

| Species | Calc. mass. (Da) | Obs. mass. (Da) | Err. (ppm) |
|---|---|---|---|
| b3 | 357.1563 | 357.1567 | 1.1 |
| b9 | 944.4301 | 944.4306 | 0.5 |
| b10 | 1031.4621 | 1031.4636 | 1.5 |
| b11 | 1146.4890 | 1146.4904 | 1.2 |
| b12 | 1293.5575 | 1293.5599 | 1.9 |
| b14 | 1451.6266 | 1451.6280 | 1.0 |
| y2 | 221.0955 | 221.0949 | -2.7 |
| y4 | 379.1646 | 379.1639 | -1.8 |
| $[M+2H]^{2+}$ | 836.3610 | 836.3635 | 3.0 |

**Figure S50.** Structural characterization via MS/MS of the SpaL-modified SpaA1.3 product.

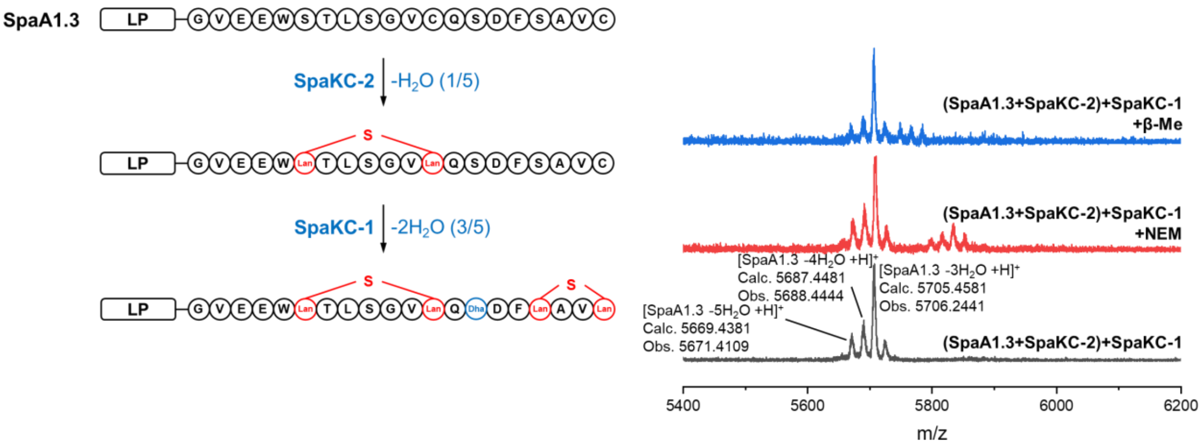

**Figure S51.** MALDI-TOF MS analysis and NEM/β-Me derivatization of the *in vitro* cascade modification product SpaA1.3–SpaKC-2–SpaKC-1.

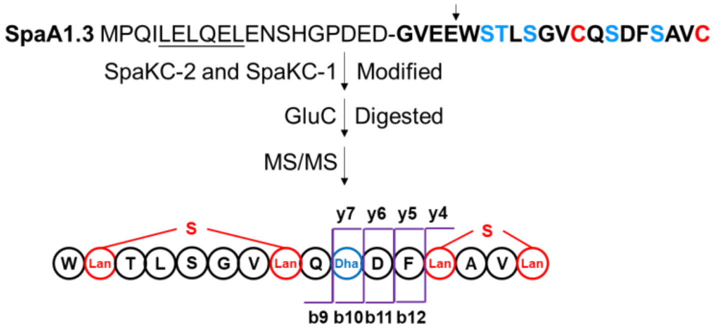

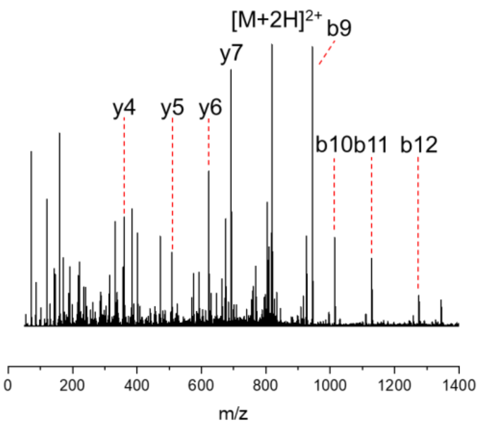

| Species | Calc. mass. (Da) | Obs. mass. (Da) | Err. (ppm) |
|---|---|---|---|
| b9 | 944.4301 | 944.4326 | 2.6 |
| b10 | 1013.4521 | 1013.4544 | 2.3 |
| b11 | 1128.4790 | 1128.4811 | 1.9 |
| b12 | 1275.5475 | 1275.5477 | 0.2 |
| y4 | 361.1546 | 361.1541 | -1.4 |
| y5 | 508.2230 | 508.2229 | -0.2 |
| y6 | 623.2500 | 623.2506 | 1.0 |
| y7 | 692.2720 | 692.2724 | 0.6 |
| [M+2H]$^{2+}$ | 818.3510 | 818.3528 | 2.2 |

**Figure S52.** Structural characterization via MS/MS of the cascade modification product SpaA1.3–SpaKC-2–SpaKC-1.

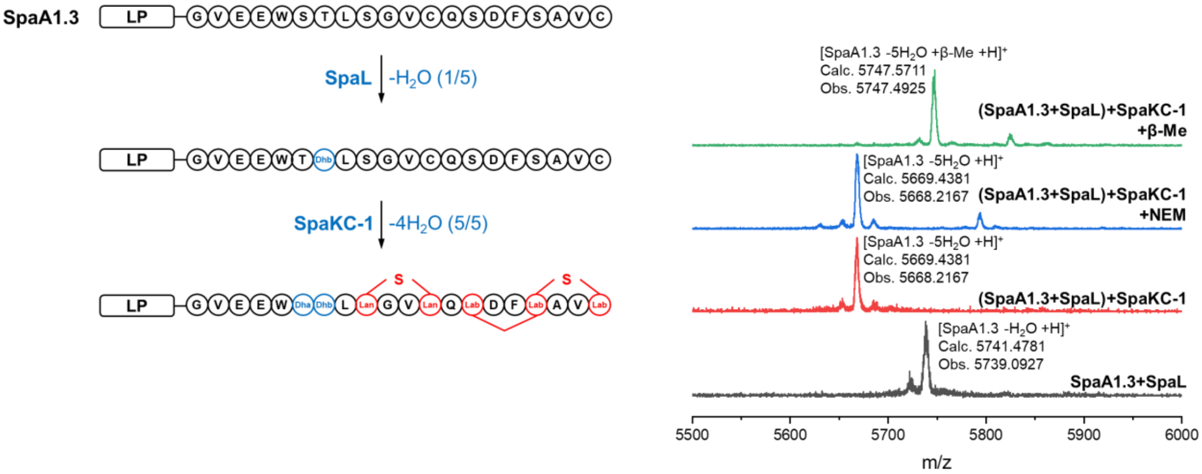

**Figure S53.** MALDI-TOF MS analysis and NEM/β-Me derivatization of the *in vitro* cascade modification product SpaA1.3–SpaL–SpaKC-1.

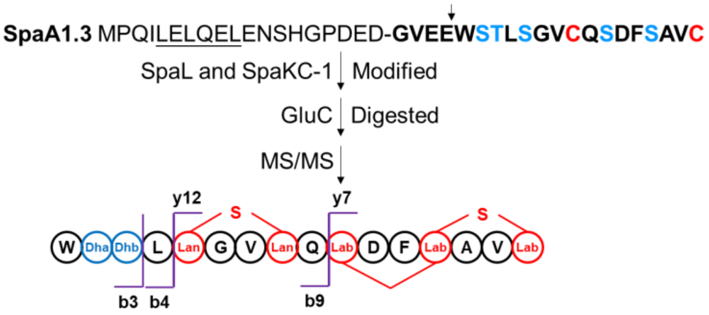

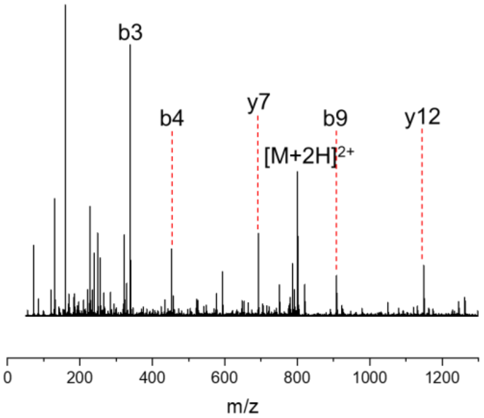

| Species | Calc. mass. (Da) | Obs. mass. (Da) | Err. (ppm) |
|---|---|---|---|
| b3 | 339.1463 | 339.1454 | -2.7 |
| b4 | 452.2304 | 452.2295 | -2.0 |
| b9 | 908.4101 | 908.4095 | -0.7 |
| y7 | 692.2720 | 692.2726 | 0.9 |
| y12 | 1148.4517 | 1148.4543 | 2.3 |
| $[M+2H]^{2+}$ | 800.3410 | 800.3417 | 0.9 |

**Figure S54.** Structural characterization via MS/MS of the cascade modification product SpaA1.3–SpaL–SpaKC-1.

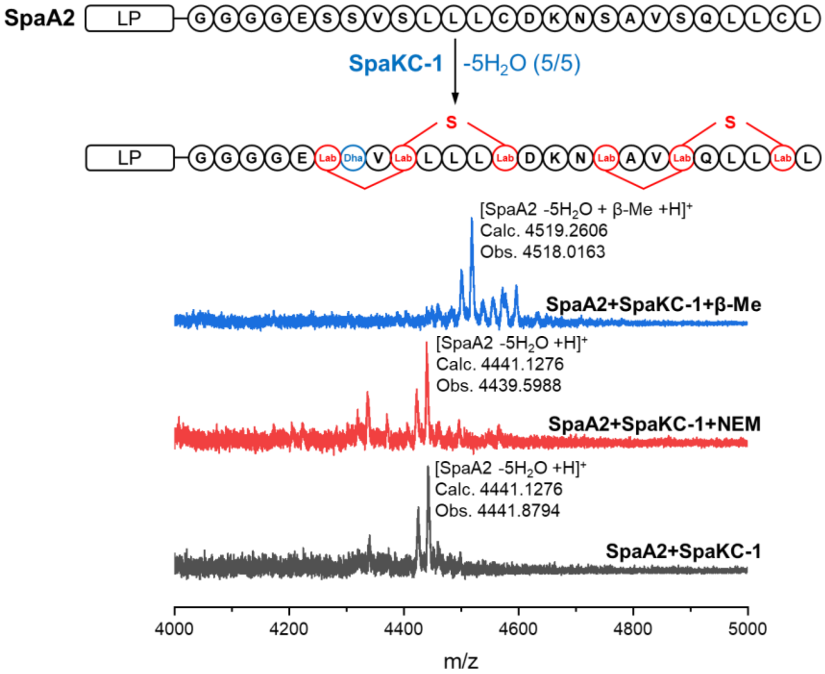

**Figure S55.** MALDI-TOF MS analysis and NEM/β-Me derivatization of product by *in vitro* modification of SpaKC-1 from *lan III-1* and precursor peptide SpaA2 from *lan III-2*.

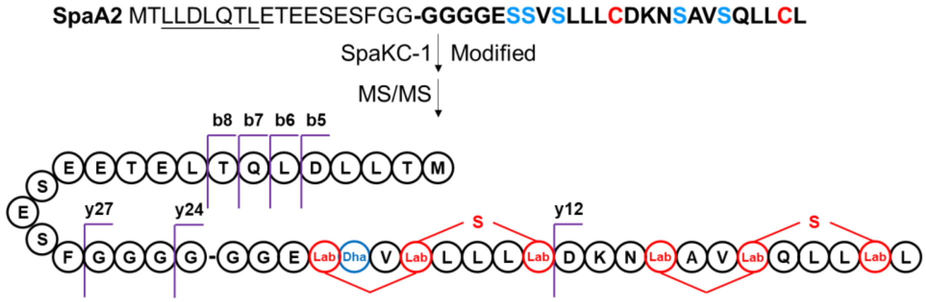

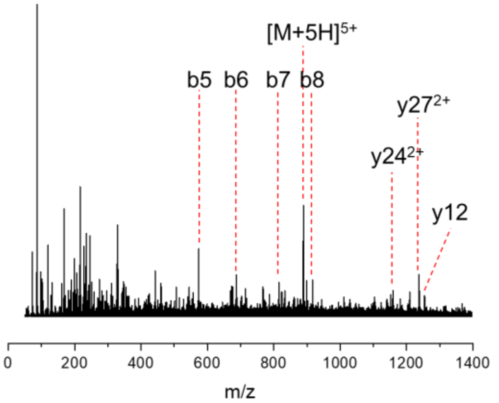

| Species | Calc. mass. (Da) | Obs. mass. (Da) | Err. (ppm) |
|---|---|---|---|
| b5 | 574.2906 | 574.2926 | 3.5 |
| b6 | 687.3746 | 687.3759 | 1.9 |
| b7 | 815.4332 | 815.4337 | 0.6 |
| b8 | 916.4809 | 916.4859 | 5.5 |
| y12 | 1254.6523 | 1254.6591 | 5.4 |
| $y24^{2+}$ | 1152.0812 | 1152.0866 | 4.7 |
| $y27^{2+}$ | 1237.6134 | 1237.6176 | 3.4 |
| $[M+5H]^{5+}$ | 889.0314 | 889.0349 | 3.9 |

**Figure S56.** Structural characterization via MS/MS of the SpaKC-1-modified SpaA2 product.

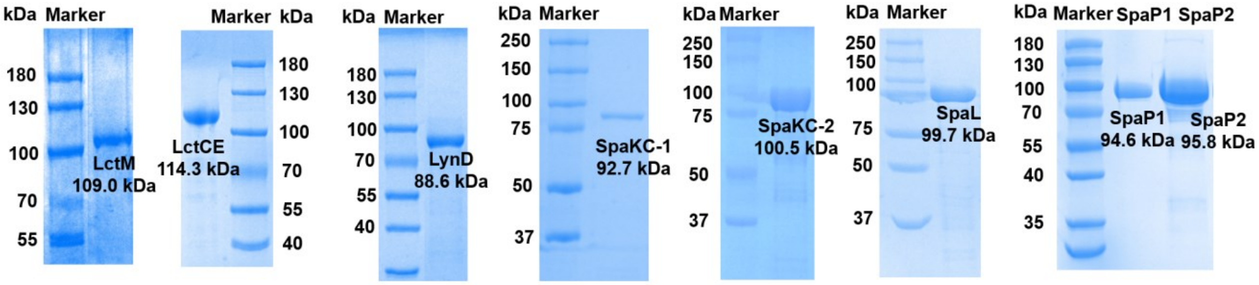

**Figure S57.** SDS-PAGE analysis of purified enzymes used in this work (LctM, LctCE[4], LynD, SpaKC-1, SpaKC-2, SpaL, SpaP1 and SpaP2). The gels were stained with Coomassie Blue.

### LC–MS analysis for leader peptide probes

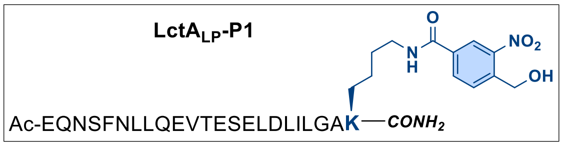

**LctA$_{LP}$-P1** was prepared as a white powder from Rink MBHA Amide following the **Pathway 1**.

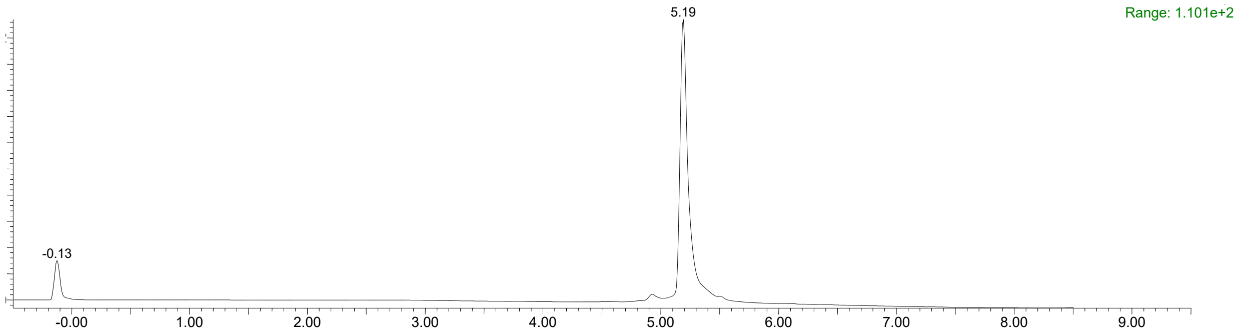

$t_R$ = 5.19 min, 10%-90% B for 8 min, then 90% B 8-10 min. **Purity**: 94.9%.

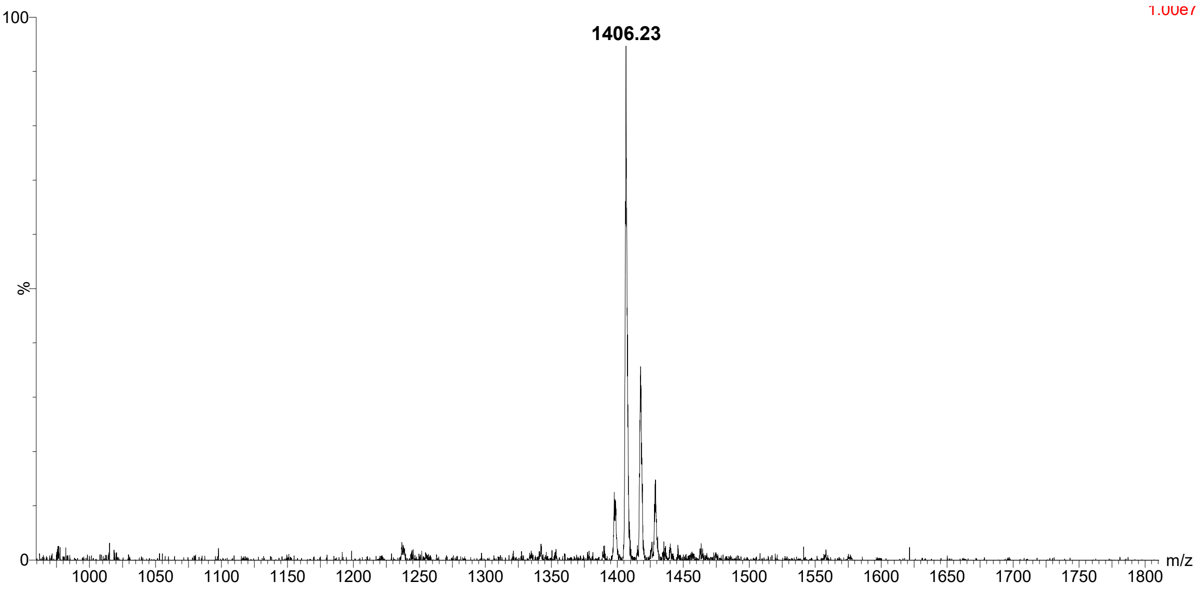

**ESI-MS:** Calculated for $C_{123}H_{192}N_{30}O_{45}$ $[M+2H]^{2+}$: 1406.19, found: 1406.23.

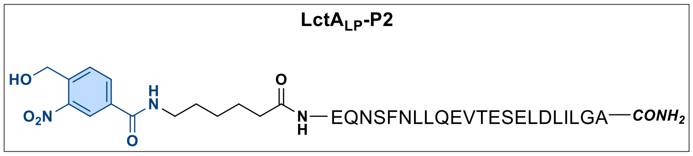

**LctA$_{LP}$-P2** was prepared as a white powder from Rink MBHA Amide following the **Pathway 2**.

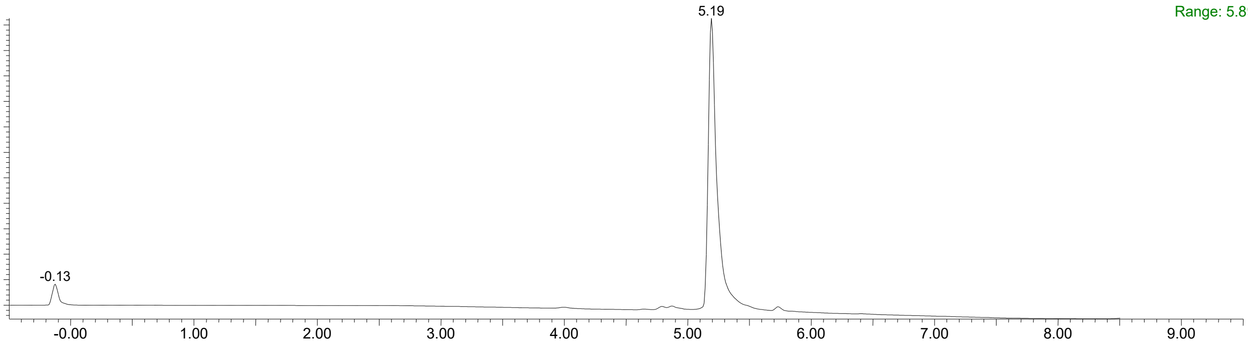

$t_R$ = 5.19 min, 10%-90% B for 8 min, then 90% B 8-10 min. **Purity**: > 98%.

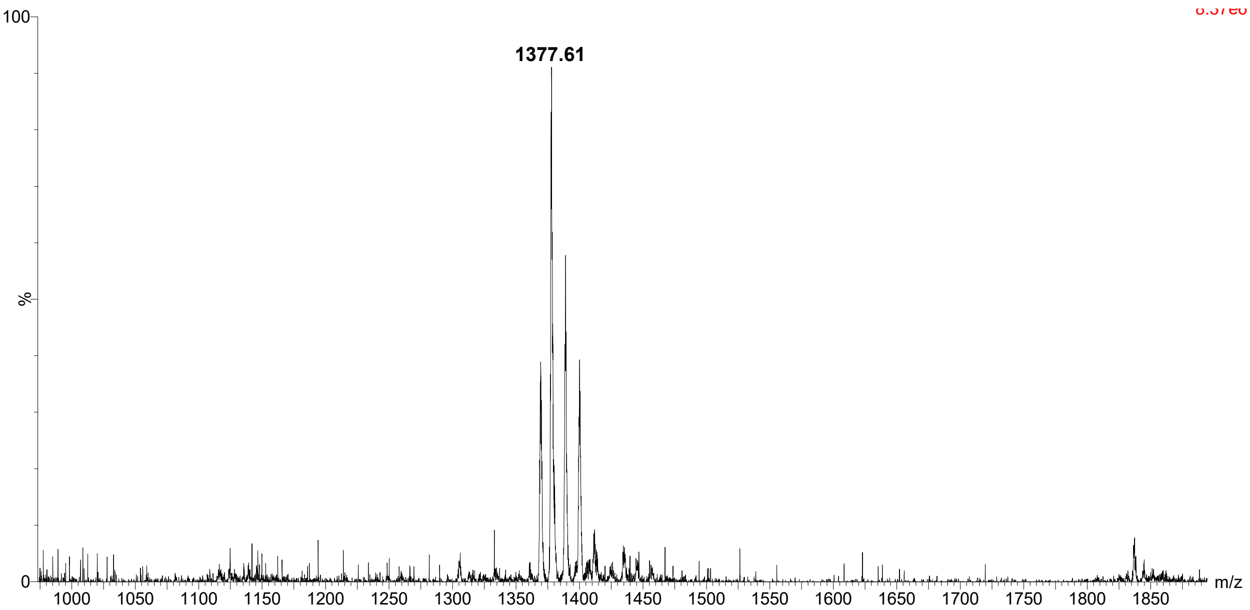

**ESI-MS:** Calculated for $C_{121}H_{189}N_{29}O_{44}$ $[M+2H]^{2+}$: 1377.68, found: 1377.61.

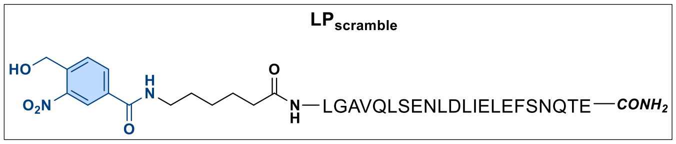

**LP$_{scramble}$** was prepared as a white powder from Rink MBHA Amide following the **Pathway 2.**

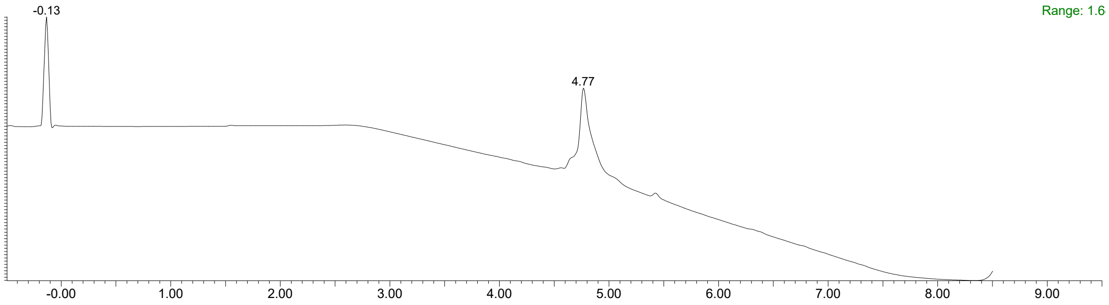

$t_R$ = 4.77 min, 10%-90% B for 8 min, then 90% B 8-10 min. **Purity**: 80%.

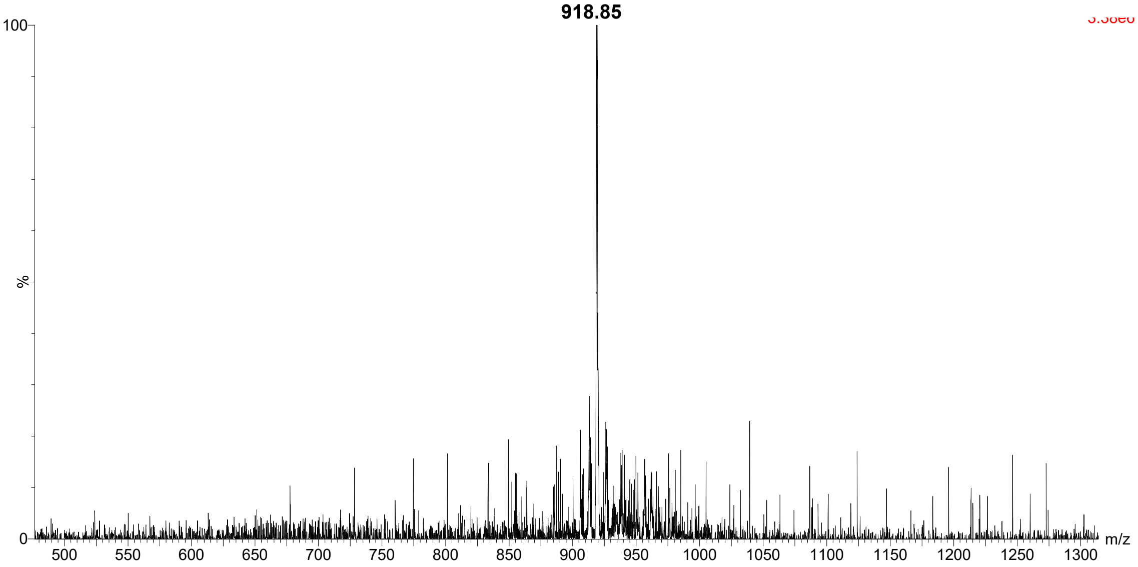

**ESI-MS:** Calculated for $C_{121}H_{189}N_{29}O_{44}$ $[M+3H]^{3+}$: 918.79, found: 918.85.

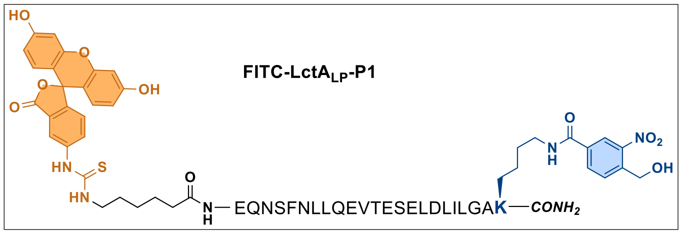

**FITC-LctA$_{LP}$-P1** was prepared as a yellow powder from Rink MBHA Amide following the **Pathway 3.**

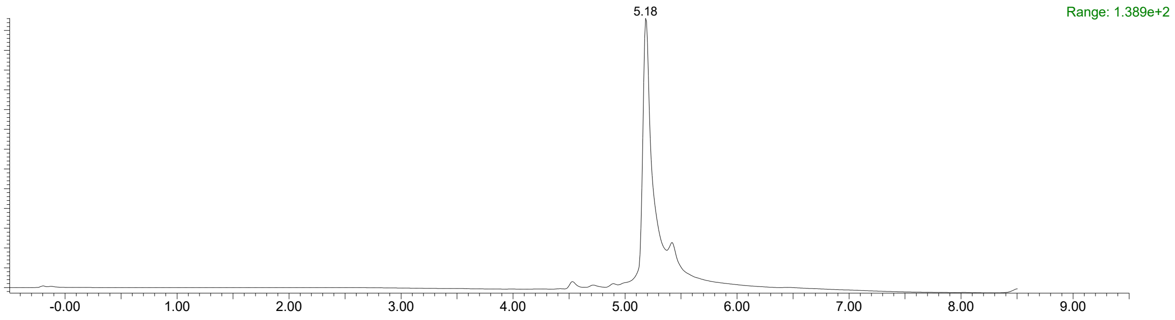

$t_R$ = 5.18 min, 10%-90% B for 8 min, then 90% B 8-10 min. **Purity**: 87%.

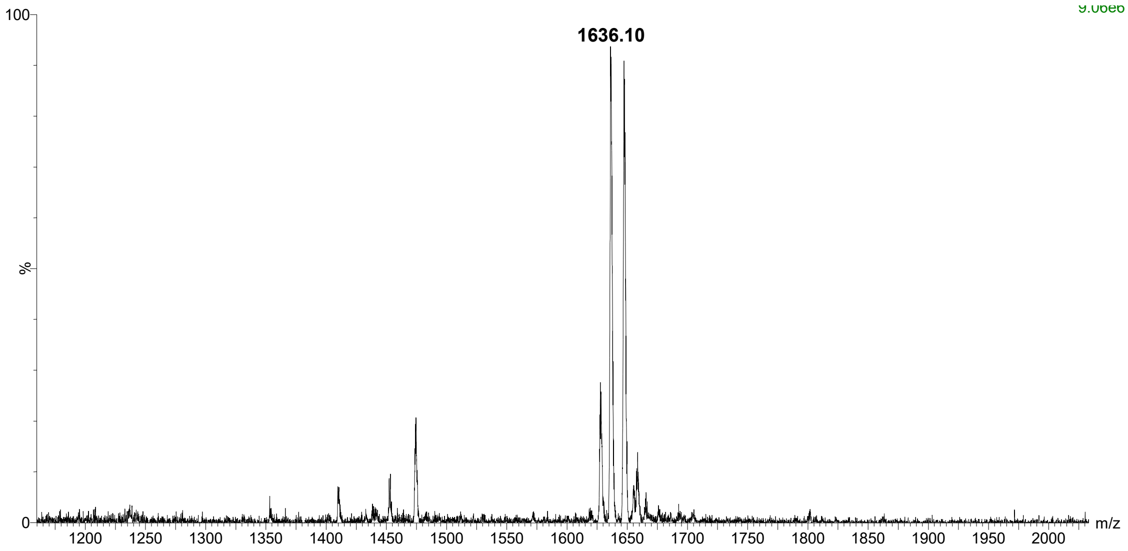

**ESI-MS:** Calculated for $C_{148}H_{212}N_{32}O_{50}S$ $[M+2H]^{2+}$: 1636.25, found: 1636.10.

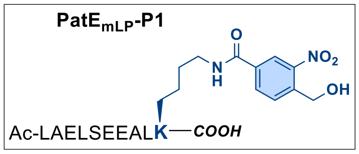

**PatE$_{mLP}$-P1** was prepared as a white powder from Rink MBHA Amide following the **Pathway 1**.

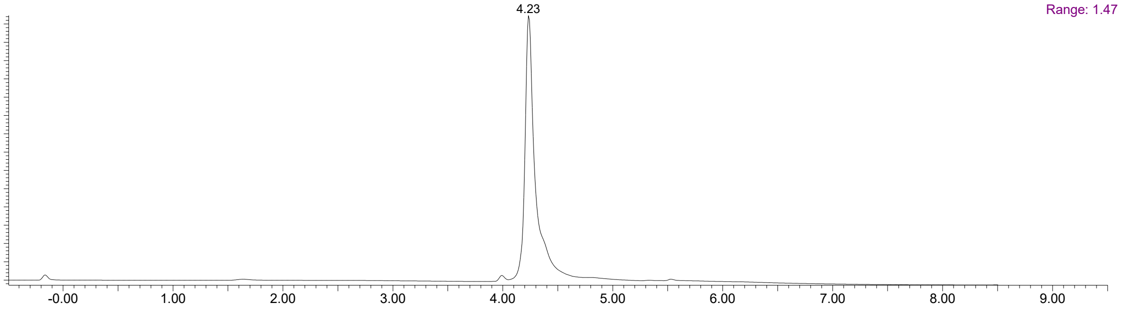

$t_R$ = 4.23 min, 10%-90% B for 8 min, then 90% B 8-10 min. **Purity**: 97%.

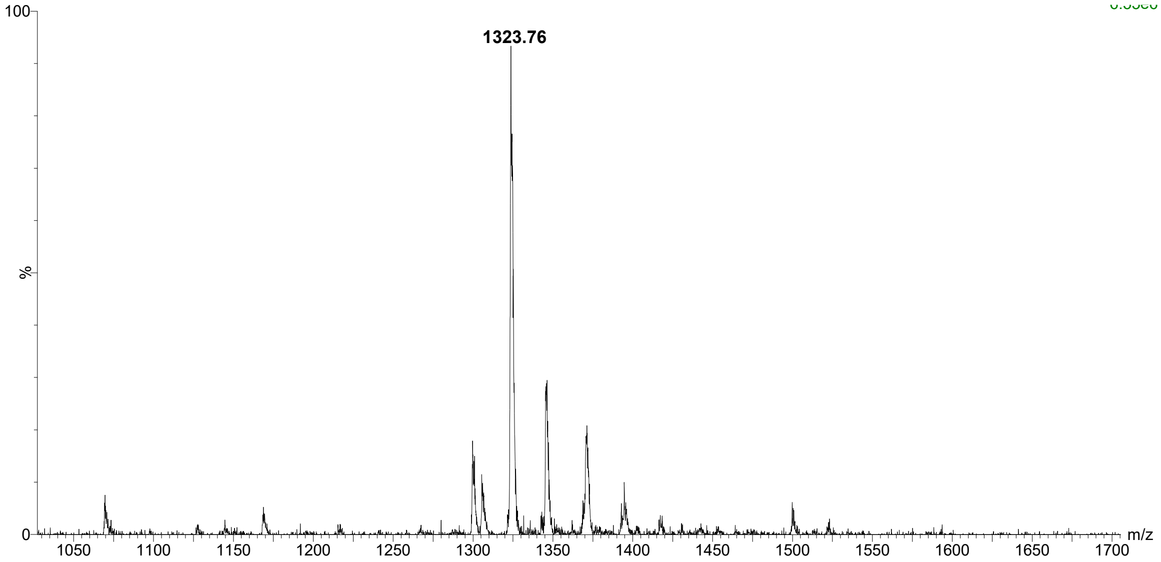

**ESI-MS:** Calculated for $C_{58}H_{90}N_{12}O_{23}$ $[M+H]^+$: 1323.63, found: 1323.76.

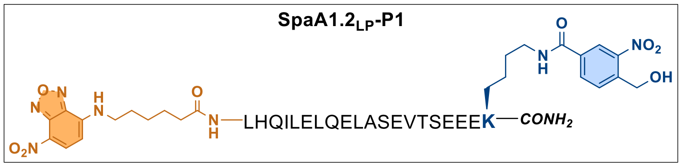

**SpaA1.2LP-P1** was prepared as a yellow powder from Rink MBHA Amide following the **Pathway 4.**

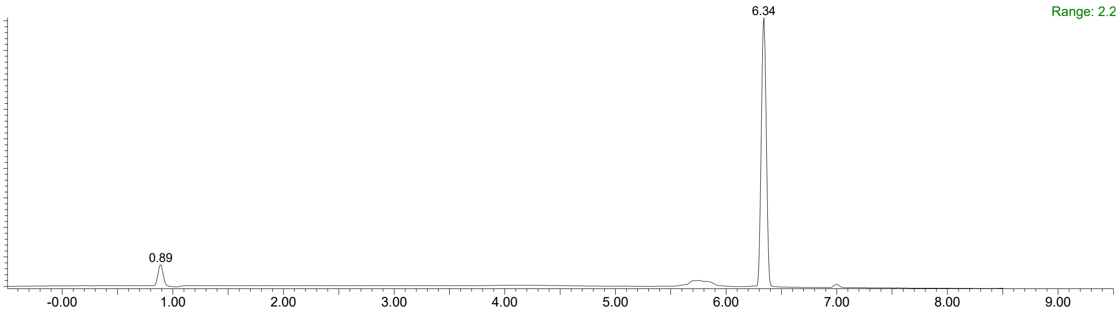

$t_R$ = 6.34 min, 10%-90% B for 8 min, then 90% B 8-10 min. **Purity**: 91%.

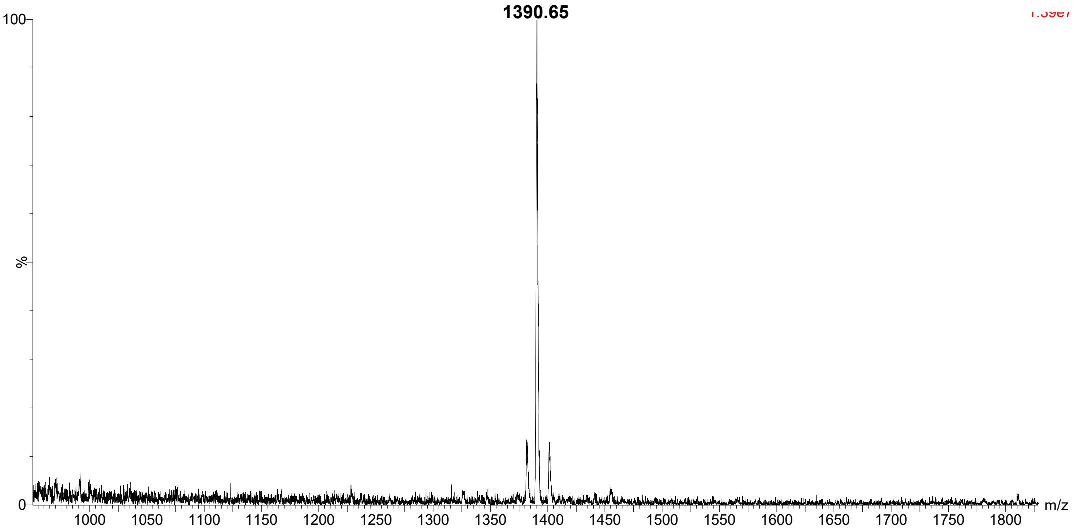

**ESI-MS:** Calculated for $C_{120}H_{183}N_{31}O_{45}$ $[M+2H]^{2+}$: 1390.66, found: 1390.65.

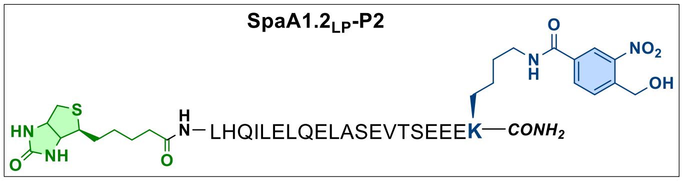

**SpaA1.2$_{LP}$-P2** was prepared as a white powder from Rink MBHA Amide following the **Pathway 5.**

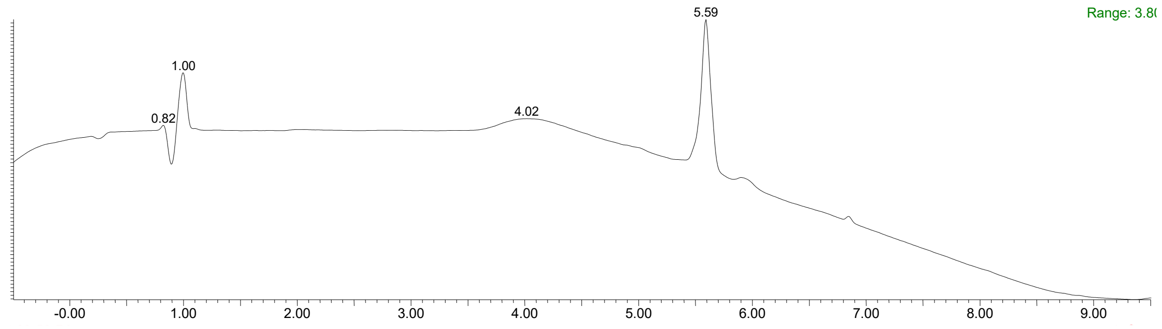

$t_R$ = 5.59 min, 10%-90% B for 8 min, then 90% B 8-10 min. **Purity**: 95%.

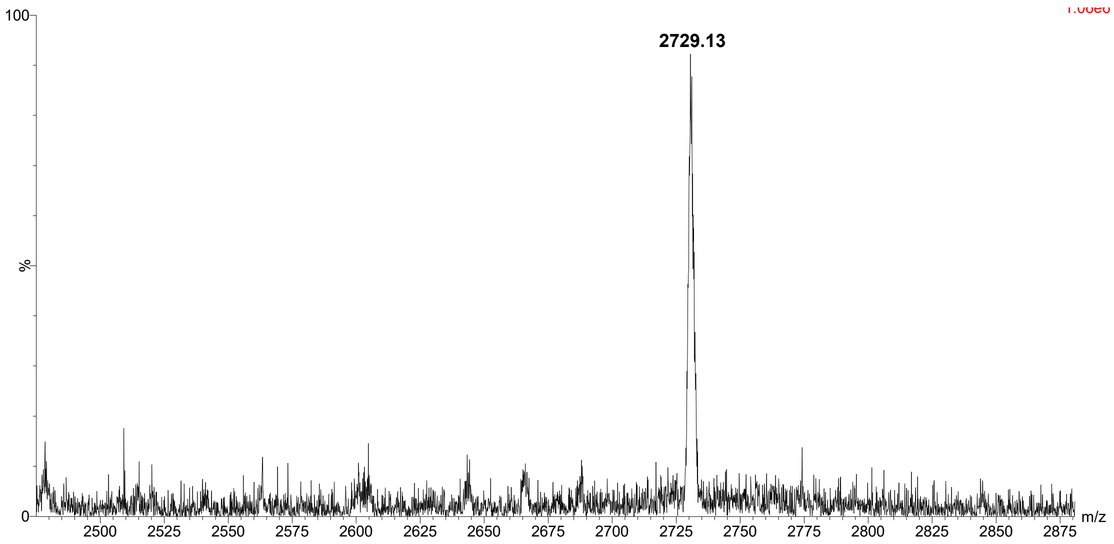

**ESI-MS:** Calculated for $C_{118}H_{185}N_{29}O_{43}S$ $[M+H]^+$: 2729.29, found: 2729.13.

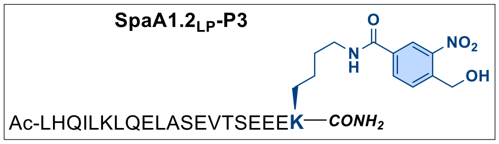

**SpaA1.2LP-P3** was prepared as a white powder from Rink MBHA Amide following the **Pathway 1**.

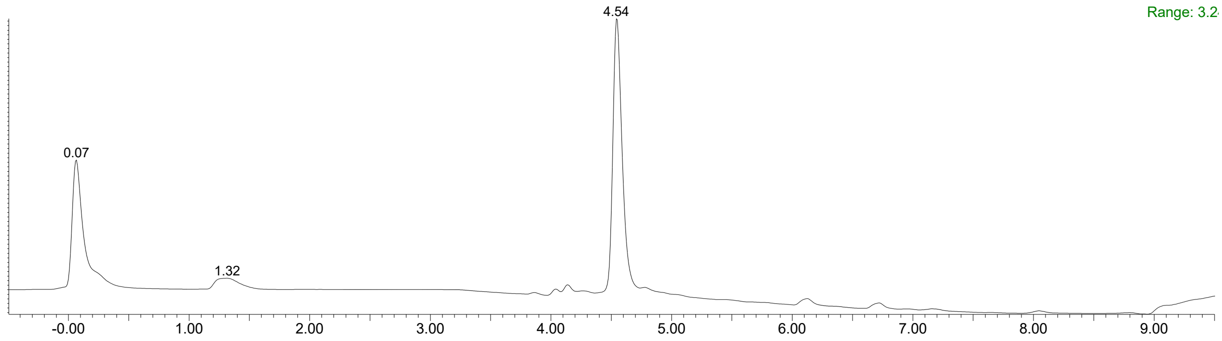

$t_R$ = 4.54 min, 10%-90% B for 8 min, then 90% B 8-10 min. **Purity**: 96%.

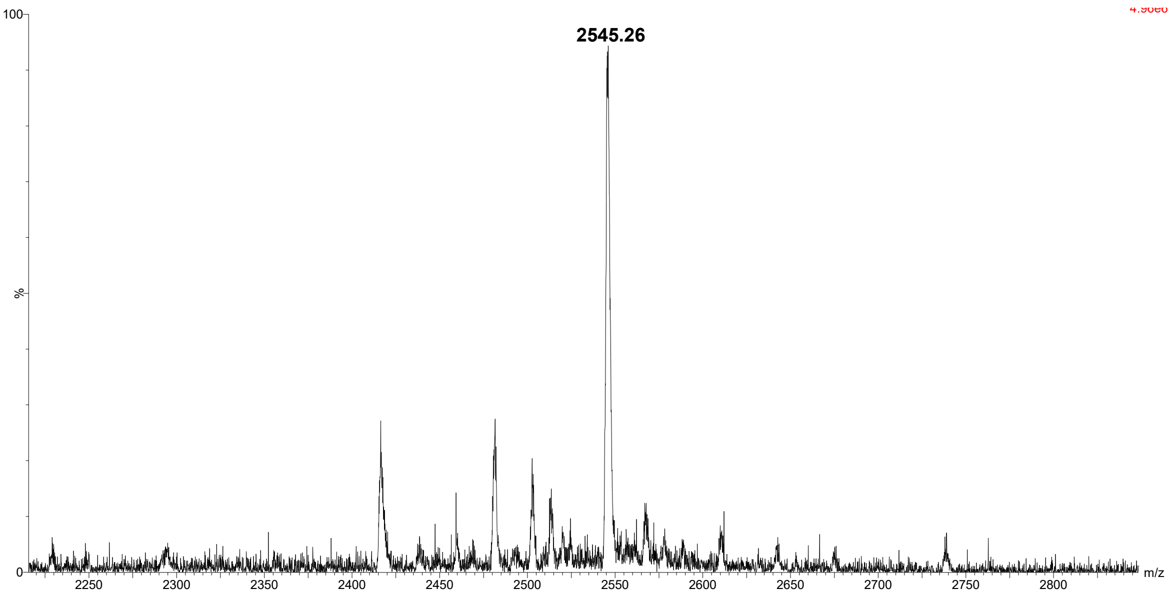

**ESI-MS:** Calculated for $C_{111}H_{178}N_{28}O_{40}$ $[M+H]^+$: 2545.28, found: 2545.26.
